# Accelerating Virtual Directed Evolution of Proteins via Reinforcement Learning

**DOI:** 10.1101/2025.06.25.661516

**Authors:** Tianyu Mi, Yuxiang Wang, Jingyu Zhao, Wanze Wang, Yunhao Shen, Nan Xiao, Ligong Chen, Guo-Qiang Chen, Shuyi Zhang, Wen-Bin Zhang, Haipeng Gong

## Abstract

With the advancement of machine learning methods, the protein fitness landscape can be predicted, providing reliable guidance in the selection of advantageous mutations for the directed evolution of proteins. However, the potential multiple mutational variants derived from the simple combination of a limited number of advantageous single mutations may not represent superior choices. Moreover, the exploration and selection of the astronomical number of multiple mutational variants remain a highly challenging task. In this study, we introduce a virtual directed evolution pipeline, RelaVDEP, for the rapid identification of mutational variants with explicit enhancement in the desired property of the target protein. By adapting and fine-tuning a pre-trained fitness predictor to improve sequence-based protein functional prediction and by designing a model-based reinforcement learning framework to efficiently explore the vast combinatorial space of protein mutations, this pipeline is capable of effectively accelerating the directed evolution process for a broad spectrum of proteins with versatile functional profiles. According to a series of experimental validations, the diversified mutational variants identified by our method exhibit notable improvements in desirable protein functional properties. In particular, by integrating RelaVDEP with active learning, we successfully optimized the sequence of a PETase, enhancing its catalytic activity through previously unknown mutations.

## Introduction

Protein engineering tailors protein properties by modifying existing proteins to meet specific needs in industry, medicine and biotechnology. As a foundational approach, directed evolution^1^ employs iterative cycles of mutagenesis and selection to accumulate beneficial mutations in proteins for property optimization. This strategy mimics Darwinian evolution through mutant library creation and artificial selection pressures, steering proteins toward improved functionality^2,3^.

Recent advances in directed evolution involve the integration of machine learning in the prediction of mutation-induced fitness changes. Unsupervised methods make predictions by learning acceptable transitions between amino acids across homologous sequences. EVmutation^4^ and DeepSequence^5^ model natural transitions using Markov random fields and variational autoencoders, respectively. ESM-1v^6^, ESM-2^7^ and MSA Transformer^8^ employ protein language models to learn and predict the probabilities of various amino acids at individual positions. Supervised methods, on the other hand, leverage deep mutational scanning (DMS) data for supervised model training, as exemplified by ECNet^9^, SESNet^10^ and GVP-MSA^11^. ProteinNPT^12^ effectively combines unsupervised predictions with supervised learning, achieving the leading position on ProteinGym^13^ benchmarks. SPIRED-Fitness^14^ stands out for its speed, accuracy and adaptability across zero-shot/supervised settings, by integrating single-sequence-based structure prediction and supervised fitness prediction in an end-to-end framework.

Furthermore, machine learning has been substantiated as effective in guiding and accelerating the directed evolution process. Examples include, but are not limited to, the Gaussian-process-based approaches adopted in GP-UCB^15^, MLDE^16^ and ALDE^17^, the clustering strategies used in CLADE^18^ and CLADE2.0^19^, the adaptive greedy search applied in AdaLead^20^, and the graph-based neural networks utilized in ProLGN^21^. Factors beyond fitness are also taken into account for the practical directed evolution guidance, as exemplified by PEX^22^ that prioritizes high-fitness mutants with minimal mutations and MODIFY^23^ that optimizes both fitness and diversity in mutant libraries. EVOLVEpro^24^ achieves single-sequence-based prediction of variable protein activity metrics in the low-*N* scenario by integrating the protein language model with a regression model, enabling iterative optimization of protein functions through active learning.

Despite these remarkable advances, existing methods still struggle to navigate the vast combinatorial space of protein sequences. DeepMind addressed a similar challenge in Go by combining Monte Carlo tree search (MCTS) with deep reinforcement learning (RL), leading to AlphaGo^25^ and its successors, AlphaGo Zero^26^, AlphaZero^27^ and MuZero^28^. Similar approaches have been applied in protein engineering but with limited success. DyNA-PPO^29^ engages modelbased RL but suffers from sparse rewards and the risk of misfolded sequences. EvoPlay^30^, inspired by AlphaZero, suffers from inefficient exploration due to its single-threaded sampling/training paradigm as well as poor generalizability owing to its rudimentary fitness predictor. KnowRLM^31^ relies on known experimental data, weakening its effectiveness in few-shot or zero-shot settings. Moreover, these prior attempts focus on amino acid sequences for functional optimization, overlooking critical inter-residue relationships within the three-dimensional protein structures.

In this study, we present RelaVDEP (<u>Re</u>inforcement learning <u>a</u>ssisted <u>V</u>irtual <u>D</u>irected <u>E</u>volution for <u>P</u>roteins), a model-based RL framework specifically designed to optimize protein functions through a virtual directed evolution process. The framework integrates a high-precision pre-trained protein fitness predictor as its reward function and employs a graph neural network (GNN) architecture to explicitly encode the structure-aware inter-residue relationships. Built with a distributed computational architecture, RelaVDEP supports a parallelized training process. Additionally, a multi-objective optimization strategy is designed to construct the mutant library that systematically balances functional fitness and sequence diversity. Experimental results demonstrate that RelaVDEP achieves accurate identification of dominant mutation sites in target proteins and enables combinatorial generation of numerous beneficial mutants, lead-ing to the derivation of diversified multi-mutational sequences with marked improvement in various desired functional properties including fluorescence intensity, abundance, transcription repression, and enzyme activity.

## Results

### Overview of RelaVDEP

The RelaVDEP pipeline is shown schematically in Figure 1. In this pipeline, the varying sequence of the target protein is modeled as a state representation, with the initial state corresponding to the starting sequence and the following states reflecting the continuous mutations during the virtual directed evolution (Figure 1a). Specifically, each sequence state is represented by an individual graph, where the node and edge features are extracted from the sequence embedding of the language model ESM-2^7^ and the protein structure predicted by the single-sequence-based structure predictor SPIRED^14^. On the other hand, SPIRED-Fitness^14^ pre-trained on DMS data is chosen as a reward model. The zero-shot version of this model is adapted to allow the prediction of multiple mutational effects and then fine-tuned upon the available experimental data of the target protein, aiming to reliably score the state transitions elicited by mutations in our framework. In the RL environment, the virtual directed evolution process is modeled as a Markov Decision Process, in which each step corresponds to a transition between two different protein sequence states by a mutation action. Specifically, at step *t*, the action *a_t_* is chosen based on the policy *π_t_*. Once the action is received by the environment, a directed mutation is applied to the specific single residue, generating a new sequence, with the reward *u_t_*inferred from the reward model. A complete mutation trajectory concludes when the number of mutated residues reaches a pre-defined maximum mutation count. This process could be simulated by the MCTS, where each mutation trajectory corresponds to a tree-search path, and the simulations in return help to determine the policy *π* at individual mutation steps. Moreover, mutation trajectories are recorded in the replay buffer, allowing training and intermittent update of a neural network model, the core component of the model-based RL framework that guides the agent to carry out MCTS operations in a differentiable manner (Figure 1b). This neural network model consists of a representation function *h*(*θ*) that converts the graph-based protein sequence state into a hidden state, a prediction function *f* (*θ*) that predicts the policy and value for each hidden state, and a dynamics function *g*(*θ*) that performs a step of transition from the current hidden state based on the action and predicts the corresponding reward and the next hidden state. Overall, the entire model enables end-to-end training, effectively accelerating the exploration of protein sequence space. Finally, candidate mutational sequences suggested by the RL framework are further selected to compose an optimized mutant library, by comprehensively considering predicted fitness and sequence diversity in a low-dimensional subspace formed by DHR^32^ embedding and t-SNE^33^ clustering (Figure 1c). Computational models including SPIRED-Stab^14^, ESMFold^21^ and AlphaFold2^34^ are employed as additional filters to extract sequences with sufficient stability and foldability from the mutant library for subsequent wet-lab experimental verification.

**Figure 1.**
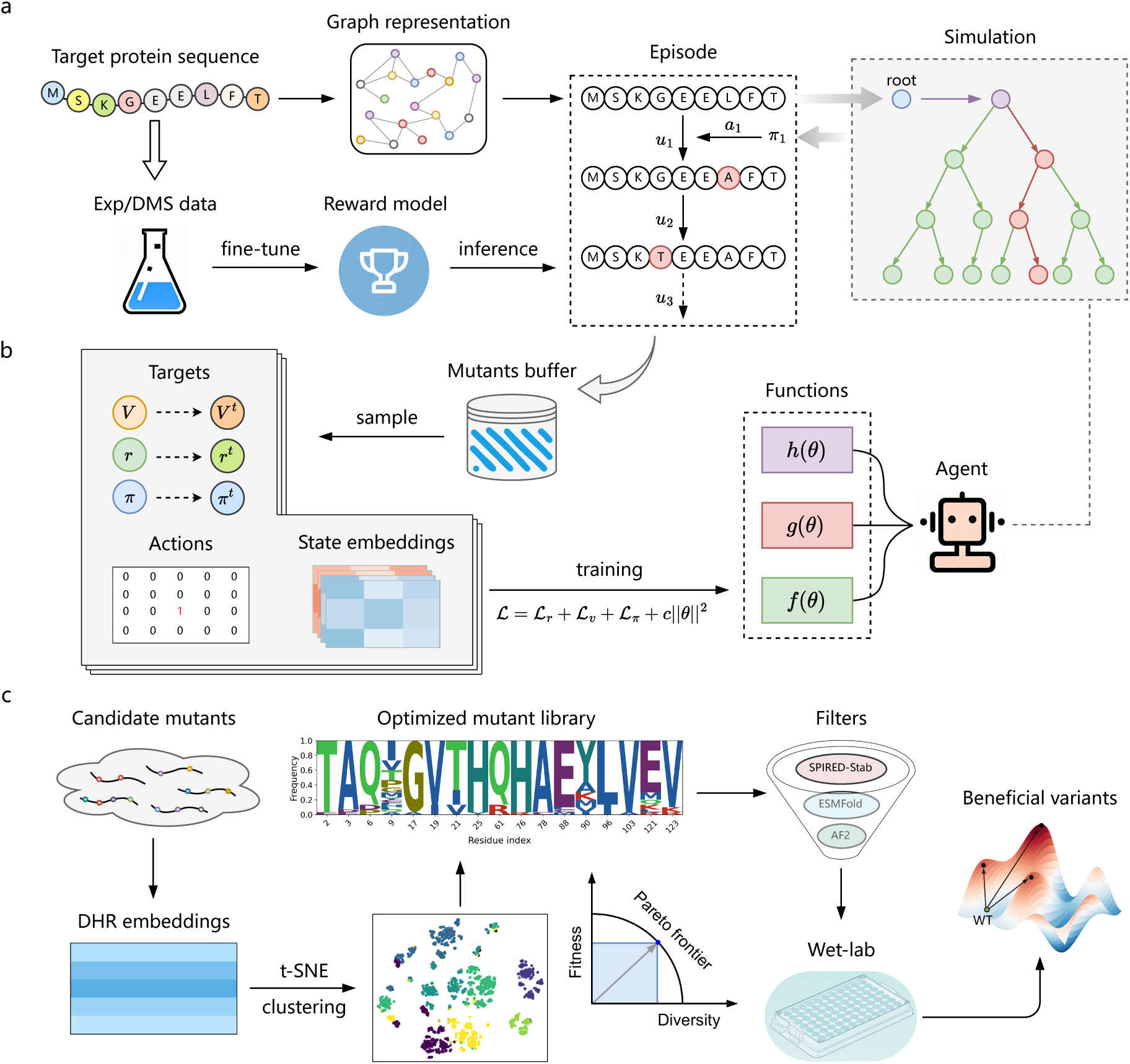
Schematic illustration of the RelaVDEP pipeline. **a)** In the virtual directed evolution environment, the protein sequence state is represented as a graph and experiences sequential mutations. At each step, the mutation action (*a*) is selected based on the policy (*π*), which is derived from hundreds of steps of MCTS simulations, and the resulting variant is endowed with a reward (*u*) by a reward model that has been optimized based on available DMS/experiment data collected for the target protein. **b)** In the RL framework, MCTS operations are guided by a neural network model during the generation of mutation trajectories. This model consists of a representation function *h*, a prediction function *f* and a dynamics function *g*, with learnable parameters (*θ*) optimized based on the sampled sequence states and their corresponding data. **c)** To obtain beneficial variants corresponding to versatile optimal peaks of the fitness landscape, candidate mutants proposed by the RL framework are selected based on both predicted fitness and sequence diversity, followed by computational filtering to ensure foldability and stability before proceeding to experimental validation.

### Evaluation of the framework on GFP

In general RL tasks, an immediate reward is obtained based on the inherent rules. Inspired by a previous study^35^, the virtual fitness reward is provided by a pre-trained predictor in our RL-assisted directed evolution task, as opposed to empirical experimental outcomes. In previous methods such as EvoPlay^30^, the reward model is constructed with convolutional neural network (CNN), a conventional architecture in RL frameworks due to its simplicity and efficiency. Here, we adapted the zero-shot version of SPIRED-Fitness^14^ to construct a graph-based reward model, which processes wild-type and mutant sequences through the inherited graph-attention (GAT) module, infers their fitness values through additional multi-layer perceptrons (MLPs), and outputs their difference as the predicted fitness change (Figure 2a). Unlike the original SPIRED-Fitness that relies on single and double mutational data on the wild-type sequence for training, our modified version allows for optimization using variants carrying arbitrary mutations and is therefore capable of providing rapid and reliable prediction of multiple mutational effects.

**Figure 2.**
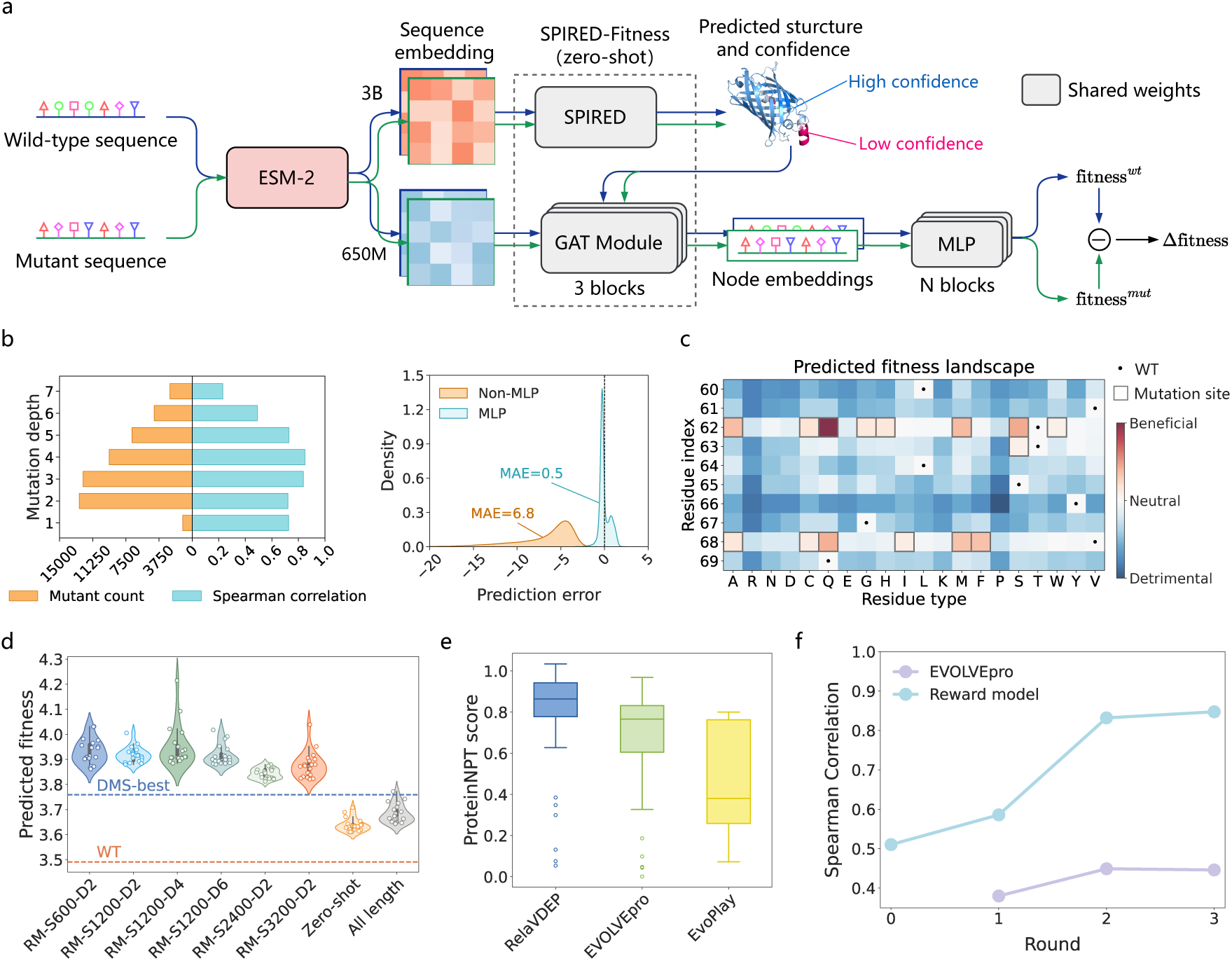
Proof of principle of RelaVDEP on GFP. **a)** The overall architecture of the reward model. Specifically, modules enclosed in the dashed box are directly inherited from the zero-shot version of SPIRED-Fitness, and additional MLP layers are appended to enable model fine-tuning on available DMS or experimental data. Both wild-type and mutant sequences are fed into the network with shared weights, and the difference in their outputs is used to predict the fitness change through a linear layer. **b)** Performance of the reward model on available DMS data about the fluorescence intensity of GFP. In the left panel, the Spearman correlation coefficient (colored in cyan) between prediction and DMS labels is evaluated for variants carrying various numbers of mutations (with the corresponding mutant count colored in orange). In the right panel, the prediction error is further evaluated by MAE for models with (cyan) and without (orange) the downstream MLP layers, respectively. **c)** Schematic diagram of the mutation restriction protocol adopted by RelaVDEP. Mutations are only allowed for beneficial ones predicted based on the fine-tuned reward model. **d)** Fitness distribution of the mutants generated by RelaVDEP is compared for different hyperparameter selections, where the best DMS variant and the wild-type one are labeled as blue and orange dashed lines, respectively. In “RM-SXXX-DY”, RM refers to the fine-tuned reward model, XXX represents the number of MCTS simulations performed before each mutation step, while Y denotes the time interval (in seconds) for updating model parameters. “Zero-shot” means using the zero-shot SPIRED-Fitness to estimate rewards. “All lengths” indicates allowing mutations at all sites, unlike the default constraints on beneficial mutations in panel (**c**). **e)** Comparison of the evolutionary results obtained by different methods. Mutants generated by RelaVDEP (blue), EVOLVEpro (green) and EvoPlay (yellow) are evaluated by ProteinNPT, and the results are compared in boxplots (*n* = 120 samples), where the box limits correspond to upper and lower quartiles and the whiskers extend to 1.5 inter-quartile range. **f)** An active-learning-based evolution is performed to mimic the EVOVLEpro iterative approach, with fitness prediction undertaken by the RelaVDEP reward model and the EVOLVEpro model, respectively. Here, the model performance is evaluated by the Spearman correlation coefficient between model predictions and true DMS data. Rounds 0, 1, 2 and 3 correspond to the cumulative prediction powers on single, double, triple and quadruple mutations, respectively. Notably, EVOLVEpro is not evaluated at the beginning (*i.e.* round 0) due to the lack of zero-shot prediction capability.

In this work, we tested our method on the green fluorescent protein (GFP) from *Aequorea victoria* (avGFP) as a proof of principle. We first fine-tuned the reward model using DMS data of fluorescence intensity in ProteinGym^13^. When trained with varying amounts of available data, this model maintains a robust prediction power (Supplementary Figure 1), supporting its generalizability. In particular, our model presents an evident advantage over the CNN-based reward predictor of EvoPlay when confronted with limited training data, a few-shot scenario frequently encountered in practical protein engineering.

We further assessed the fully trained reward model on data of varying mutation depths (Figure 2b, left). The model retains a highly consistent performance for variants carrying fewer than five mutations, with the Spearman correlation coefficient (Equation 2) *ρ* > 0.7, but gradually loses its power when the number of mutations exceeds five. We determined the maximum mutation count in our RelaVDEP framework based on such analyses. In addition, the MLP layers in our reward model effectively rectify the flaw in the original SPIRED-Fitness that focuses on relative ranking but neglects absolute magnitudes, suppressing the mean absolute error (MAE) of prediction from 6.8 to 0.5 (Figure 2b, right).

With the reward model in hand, exploration of the vast protein sequence space remains a challenge. For instance, in a 100-residue protein, the number of possible sequences carrying up to 3 mutations already exceeds one billion, negating the possibility of exhaustive enumeration. To address this problem, we used the reward model to infer the fitness landscape of all single mutations, identified beneficial mutations as potential candidates, and then restricted sequence sampling to possible combinations of these candidate mutations in the RL process (Figure 2c). As shown in Figure 2d, upon various choices of hyperparameters, the RL framework demonstrates a robust capacity in identifying new variants with predicted fitness values evidently superior to all known DMS data, as evaluated by the reward model. Interestingly, when the restriction on candidate mutations (Figure 2c) is released to allow exploration of the entire sequence space, the efficiency of the RL framework drops significantly, which further confirms the positive contribution of the mutation restriction scheme adopted here.

Next, we evaluated the fitness values of the variants identified by RelaVDEP and compared them with those found by the standard EVOLVEpro and EvoPlay pipelines (Figure 2e) using ProteinNPT^12^, a state-of-the-art fitness predictor in the supervised setting of ProteinGym. Specifically, the ProteinNPT model has been sufficiently optimized on the corresponding DMS data following the standard protocol to ensure fair evaluation. Clearly, the variants identified by our RelaVDEP framework exhibit significantly higher ProteinNPT scores than those generated by the other two pipelines, supporting the advantage of our method in facilitating practical directed evolution. Alternatively, we investigated the potential of our reward model in the active-learning-based incremental sequence optimization scheme proposed by EVOLVEpro^24^. Here, we mimicked this process as follows: 1) take all single mutational variants from the DMS dataset to train the reward model and use it to predict all double mutations in the dataset; 2) take all single and double mutational variants from the DMS dataset as experimental feedback to fine-tune the reward model for subsequent suggestions on triple mutations; 3) continue this procedure for higher-order mutations. We evaluated the model predictions by their Spearman correlation coefficient with the corresponding DMS labels at each individual stage (Figure 2f). In contrast to the gradual gain of prediction power in the EVOLVEpro model, our reward model shows a remarkably faster learning ability, with the Spearman correlation coefficient rapidly rising from 0.5 of the zero-shot version (*i.e.* round 0) to nearly 0.9 in the third iteration.

Subsequently, we collected the variants sampled by RelaVDEP and selected candidates with predicted improvement to compose a mutant library by comprehensively considering fitness and sequence diversity (Figure 1c). Here, fitness and diversity of selected variants are quantified as the mean predicted fitness across all variants and the sum of information entropies at all mutable positions (Equations 11 & 12), respectively. As shown in Figure 3a, sequences selected based on the conventional top-*N* approach (*i.e.* simply choosing *N* sequences with the highest predicted fitness scores) exhibit a clustered distribution in the 2D t-SNE subspace of DHR^32^ sequence embedding, potentially leading to redundant sequence evolution and thus increasing the follow-up experimental costs. As a contrast, the optimized mutant library generated by our approach achieves a compromised balance between fitness and diversity, surpassing both the random selection and the top-*N* approach.

**Figure 3.**
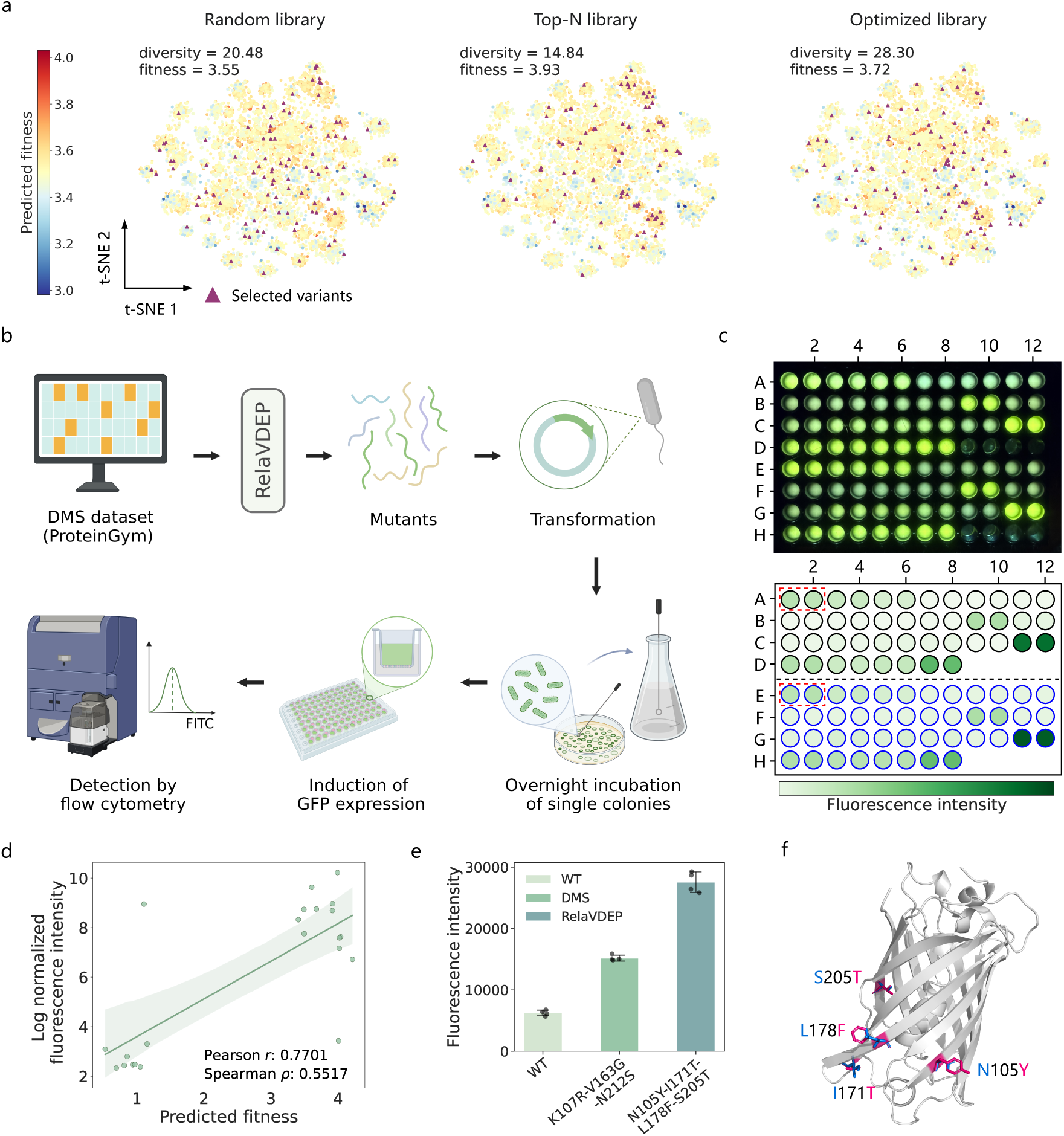
Experimental validation on the identified GFP variants. **a)** Comparison of the mutant libraries established by different approaches. Potential variants are selected from all sequences collected in the RL process, by random picking (left panel), based on the top-*N* approach focusing on predicted fitness only (middle panel), or following the approach adopted in this study (Figure 1c) to balance fitness and sequence diversity (right panel). Selected variants are projected onto the 2D t-SNE subspace of all sequences, highlighted as purple triangles. **b)** Schematic representation of the pipeline for experimental validation. **c)** Experimental results of fluorescence intensity for GFP mutants. Circles in the lower panel correspond to the fluorescent wells of the same 96-well plate in the upper panel. Each pair of adjacent circles in the horizontal direction denotes biological replicates of the same sample, with the inner color indicating fluorescence intensity. Boxes of red dashed lines mark the results of the wild-type sequence. The same column in rows A-D (encircled in black) and rows E-H (encircled in blue) represent results of the same variant with induced expression at 0.1 mM and 0.5 mM IPTG, respectively. **d)** Correlation between predicted fitness and experimentally measured fluorescence intensity. The horizontal axis represents fitness scores predicted by the fine-tuned reward model, while the vertical axis shows natural log normalized fluorescence intensity measured experimentally. **e)** Comparison of the best mutant among existing DMS data and the best variant identified by RelaVDEP against the wild-type (WT) sequence (*n* = 4 repeats). **f)** Structural characterization of the top-performing GFP quadruple mutant generated via a single round of RelaVDEP. Mutated side chains are shown in red, with the corresponding wild-type residues colored in blue.

Finally, we filtered candidate variants from this optimized mutant library using computational models to ensure foldability and stability (Figure 1c), and examined their performance against sequences filtered similarly but with low predicted fitness by wet-lab experimental verification. Specifically, all mutants were transformed into the recombinant strain of *E. coli* for Isopropyl-*β*-D-thiogalactopyranoside (IPTG) inducible expression, and the fluorescence intensity of the corresponding cells was detected by flow cytometry (Figure 3b). As shown in Figure 3c and Supplementary Table 1, duplicate measurements for the same mutant expressed at the induction levels of 0.1 mM and 0.5 mM IPTG exhibit highly consistent intra-group and across-group behaviors, allowing the use of the mean of four independent measurements to represent the fluorescence intensity for each mutant. After normalization, the experimental fluorescence intensity shows a considerable correlation (Pearson correlation coefficient *r* = 0.77) with the fitness predicted by the reward model (Figure 3d), further confirming the effectiveness of our reward model in the evaluation of multiple mutational effects. Noticeably, the best-performing variant identified by our framework (*i.e.* N105Y-I171T-L178F-S205T in Supplementary Table 1) carries a previously unknown quadruple mutation (see Supplementary Information 1.1.3 for a detailed discussion), and demonstrates an improvement in the fluorescence intensity of four-fold over the wild-type avGFP and of nearly two-fold over the best variant among the DMS data that were used to train our reward model, respectively (Figure 3e). Interestingly, when projected onto the structure of avGFP (Figure 3f), the four mutations are scattered sparsely, showing a uniform tendency to increase the size of side chains, which potentially enhances structural rigidity and thus suppresses unnecessary vibration modes relevant to fluorescence quenching.

### Enhancing protein’s cellular abundance through RelaVDEP

Nudix Hydrolase 15 (NUDT15), a member of the Nudix hydrolase superfamily, participates in the metabolism of purine-based therapeutic agents through catalytic hydrolysis of thioguanine triphosphate (TGTP). Variations in NUDT15’s cellular abundance directly influence the dosage requirements during clinical interventions using medications such as azathioprine (AZA) and mercaptopurine (6-MP)^36^. Vitamin K Epoxide Reductase Complex Subunit 1 (VKOR1), the pivotal enzyme that governs vitamin K recycling, serves as an inhibitory target of the widely used anticoagulant warfarin^37^. The cellular abundance of VKOR1 is thus of importance for the regulation of coagulation factor activation in medicine. Through high-throughput mutational analysis, Suiter et al.^38^ and Chiasson et al.^39^ systematically quantified possible single mutations for NUDT15 and VKOR1, respectively, along with their corresponding effects on cellular abundance. While these investigations enabled the preliminary construction of fitness landscapes based on single mutational variants, the combinatorial effects of multiple mutations on protein’s cellular abundance remain unexplored.

In this work, we employed these reported DMS datasets to fine-tune the reward model and implemented RelaVDEP to virtually evolve NUDT15 and VKOR1, respectively, with the optimization explicitly towards abundance augmentation. The identified mutants were then fused with the enhanced green fluorescent protein (eGFP) and transformed into HEK293T cells (*i.e.* a derivative of the human embryonic kidney 293 cell strain) for expression, with mCherry (*i.e.* a red fluorescent protein variant) co-transformed as the background (Figure 4a). Hence, the ratio of eGFP and mCherry signals in flow cytometry could be taken as the experimental abundance values of the variants.

**Figure 4.**
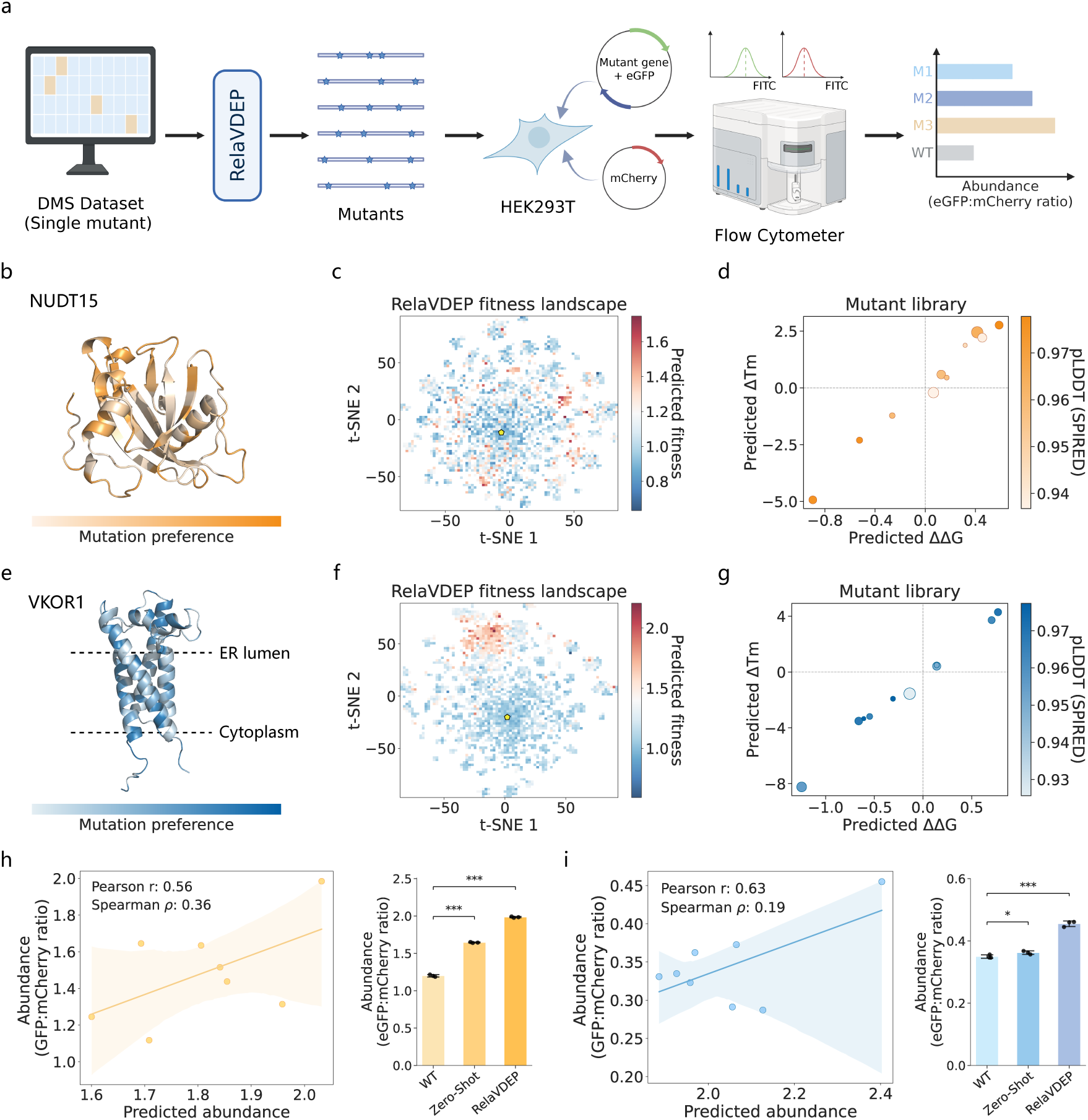
Optimization of protein’s cellular abundance for NUDT15 and VKOR1. **a)** Schematic description of the directed evolution strategy for engineering protein abundance with RelaVDEP. In experimental validation, the ratio of eGFP and mCherry signals is used to represent the cellular abundance of the target protein. **b)** Structure of the wildtype NUDT15 protein colored by site-specific mutation preferences in RelaVDEP. Darker colors indicate higher mutation frequencies. **c)** Projection of NUDT15 sequences sampled by RelaVDEP in the 2D t-SNE subspace. All samples are colored based on fitness predicted by the reward model. The yellow pentagon denotes the wild-type sequence. **d)** The optimized low-*N* mutant library of NUDT15 constructed via RelaVDEP. Each circle represents a variant, with the horizontal and vertical axes corresponding to ΔΔ*G* and Δ*T_m_* predicted by SPIRED-Stab, respectively, and the color denoting foldability (pLDDT) predicted by SPIRED. Circle size reflects the fitness predicted by the reward model. **e)** Structure of the wild-type VKOR1 protein colored by site-specific mutation preferences in RelaVDEP. Darker colors indicate higher mutation frequencies. Horizontal dashed lines indicate boundaries of the endoplasmic reticulum (ER) membrane. **f)** Projection of VKOR1 sequences sampled by RelaVDEP in the 2D t-SNE subspace, following the same conventions as in panel (**c**). **g)** The optimized low-*N* mutant library of VKOR1 constructed via RelaVDEP, following the same conventions as in panel (**d**). **h)** The left panel shows the correlation between experimental abundance and prediction of the reward model for NUDT15, while the right panel compares the abundance of the top variant identified by the default RelaVDEP pipeline against the wild-type (WT) protein (*n* = 3 repeats). “Zero-Shot” means that the allowable actions in the RL system are provided by the zero-shot SPIRED-Fitness model instead. “***” and “*” indicate statistical significance levels in the Student’s t test, with *p*-value < 0.001 and 0.05, respectively. **i)** Validation results for VKOR1 mutants, following the same conventions as in panel (**h**).

Given that the DMS data used to train the reward model consist exclusively of single mutations, we restricted the maximum mutation count to three substitutions to mitigate the impacts of neglecting epistatic effects. Following the virtual directed evolution process, we systematically analyzed mutational preferences (Equation 13) for improved variants with predicted fitness exceeding that of the wild-type sequence (Figure 4b&e). Notably, dominant mutation sites in the identified triple mutational variants are highly consistent with the advantageous single mutations in the DMS datasets (Supplementary Figures 3 & 4), laterally validating the faithful and unbiased sampling by RelaVDEP.

We reconstructed the fitness landscapes using all variants (up to triple mutations) generated by RelaVDEP for NUDT15 (Figure 4c) and VKOR1 (Figure 4f), respectively. The expanded fitness landscapes of both proteins reveal a similar pattern: high-fitness mutants cluster in distinct topological regions in the 2D t-SNE subspace. Following the approach in Figure 1c to balance fitness and diversity, we built optimized mutant libraries for NUDT15 (Figure 4d) and VKOR1 (Figure 4g), respectively. Using SPIRED-Stab^14^ as a multi-parametric filter, we prioritized candidate mutants for experimental verification, based on predicted foldability (by pLDDT), predicted changes in stability (by ΔΔ*G* and Δ*T_m_*) and predicted fitness scores. By this means, we obtained five variants with predicted improvement in cellular abundance for both NUDT15 and VKOR1. Alternatively, we used the zero-shot version of SPIRED-Fitness instead of the fine-tuned reward model to predict beneficial mutations as allowable mutation actions (Figure 2c) but kept exactly the same protocol in the follow-up works. In this way, we obtained three additional variants for each of the two protein targets. For NUDT15, the experimental measurement on the total of eight variants (Supplementary Table 2) is well correlated (Pearson correlation coefficient *r* = 0.56) with prediction of the reward model (Figure 4h, left), and the top variant selected by RelaVDEP (*i.e.* S4P-V14I-Q61I in Supplementary Table 2) shows a significant improvement of approximately 65% over the wild-type protein (Figure 4h, right). Regarding VKOR1, despite the robust Pearson correlation, the weak Spearman correlation indicates less satisfactory performance of the reward model in ranking estimation (Figure 4i, left), which negatively affects both the selection of permissible actions and the evaluation of the variants sampled. Not unexpectedly, the variant identified using the action space defined based on the zero-shot model only shows a marginal improvement with weak statistical significance (Figure 4i, right). Nevertheless, the top variant suggested by the standard RelaVDEP pipeline (*i.e.* R33D-R35Y-E155F in Supplementary Table 3) still exhibits a significant enhancement of approximately 30% in cellular abundance.

Hence, a single round of RelaVDEP-based sequence evolution has already enabled the identification of previously unknown triple mutational variants with significantly improved cellular abundance for both proteins, expediting the expansion and exploration of their fitness landscapes.

### Tuning regulatory transcription factors through RelaVDEP

AmeR, a transcriptional regulator of the TetR family, plays a pivotal role in synthetic biology by mediating the transcriptional regulation of downstream genes through interactions with specified DNA sequences and small-molecule ligands^40^. In a recent work named EvoAI^41^, the authors explored the fitness landscape of this protein through sequence space compression and repression efficiency (*i.e.* fold repression) evaluation across 82 mutational variants. More importantly, using the GeoFitness model to predict mutational effects, they successfully improved the repression power of AmeR through stochastic mutagenesis in the combinatorial space of 52 “legal” mutations.

In comparison to GeoFitness^42^ that relies on the slow, computationally demanding AlphaFold2^34^ and FoldX^43^ to extract structural features, the reward model in RelaVDEP is adapted from the high-throughput SPIRED-Fitness^14^ model that attains similar or superior performance within negligible inference time. Moreover, the RL framework is inherently advantageous in exploring the vast combinatorial space over the random mutagenesis approach. Hence, we implemented RelaVDEP to optimize the repression power of AmeR and verified the identified mutants using an experimental procedure similar to that of EvoAI^41^. Specifically, a candidate variant is transformed within a plasmid composed of an AmeR regulatory operon, and its fold repression is therefore obtainable from the expression level of the reporter gene, which is the yellow fluorescent protein (YFP) here (Figure 5a).

**Figure 5.**
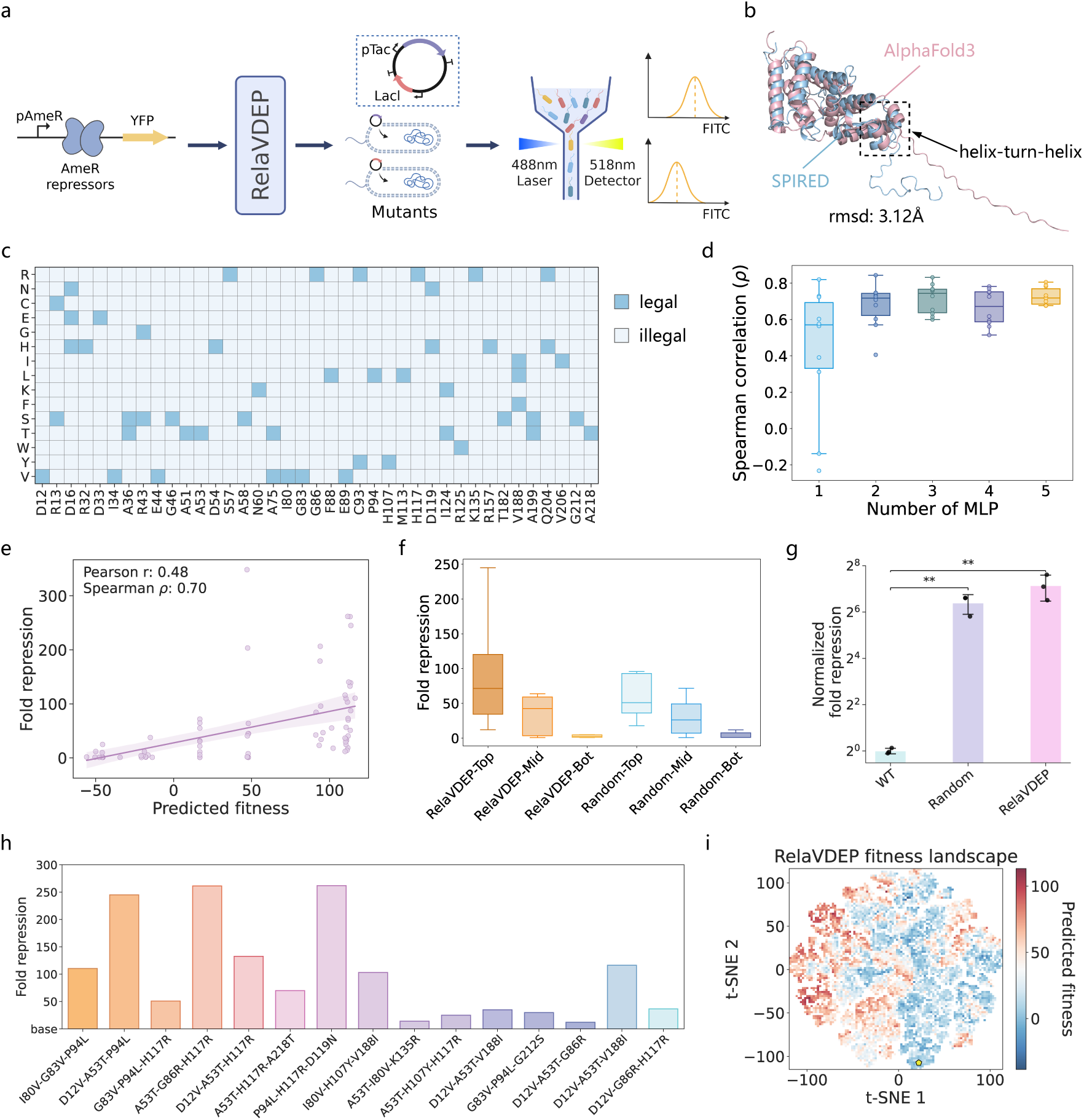
Optimization of repression power for AmeR. **a)** Schematic diagram of the RelaVDEP-mediated directed evolution strategy for optimizing the repression efficiency of transcription factor AmeR. **b)** Comparative structural analysis between SPIRED and AlphaFold3 predictions. The region in the dashed box highlights the typical helix-turn-helix (HTH) domain of TetR family proteins. **c)** Permissible mutation space mapping of AmeR. The “legal” (dark blue) and “illegal” (light blue) labels define allowed and disallowed mutational options in the RL system. **d)** 10-fold crossvalidation of the reward model with a varying number of MLP layers. The Spearman correlation coefficient between predictions and experimental values is chosen as the evaluation metric here. In the boxplots (*n* = 10 samples), the box limits correspond to upper and lower quartiles and the whiskers extend to 1.5 inter-quartile range. **e)** Correlation between predicted fitness and experimentally measured fold repression. **f)** Comparative analysis of transcriptional repression efficiency between variants explored by RelaVDEP and those generated by random evolution, stratified by predicted fitness tertiles (Top: high; Mid: medium; Bot: low). In the boxplots (*n* = 24 samples for RelaVDEP-Top, *n* = 10 samples for the others), the box limits correspond to upper and lower quartiles and the whiskers extend to 1.5 inter-quartile range. **g)** Benchmark on the fold repression of the top-performing variant explored by RelaVDEP vs. that generated by random evolution (*n* = 3 repeats). The original fold repression is normalized by the wild-type (WT) value in the vertical axis. “**” indicates statistical significance (*p*-value < 0.01) in the Student’s t test. **h)** Epistatic analysis of RelaVDEP-suggested high-performing variants. The “base” in the vertical axis represents seven common mutations (D33E, I34V, A51T, S57R, A75T, C93R & Q204R) shared across all 15 decuple mutational variants. **i)** Projection of sequence sampled by RelaVDEP in the 2D t-SNE subspace. All samples are colored based on fitness predicted by the reward model. The yellow pentagon denotes the wild-type sequence.

In the absence of experimental AmeR structures, the agreement between structures predicted by SPIRED and the state-of-the-art AlphaFold3^44^ (Figure 5b) supports the proper extraction of structural features in our reward model. Following EvoAI^41^, we restricted the action space to the combination of 52 “legal” mutations distributed over 39 sites (Figure 5c). Moreover, after fine-tuning on the limited amount of data released from EvoAI, the reward model exhibits a strong prediction power, with the Spearman correlation coefficient approaching 0.7 for a varying number of MLP layers in the 10-fold cross-validation (Figure 5d).

After a single round of the RelaVDEP-based sequence evolution, we stratified all sampled variants into three tiers of different predicted fitness values (high, medium and low) and applied computational filters to choose candidate mutants from them for experimental verification. As shown in Figure 5e, fitness prediction and experimental measurement show a moderate correlation in magnitude (Pearson correlation coefficient *r* = 0.48) and a stronger correlation in ranking (Spearman correlation coefficient *ρ* = 0.70), supporting the robustness of our reward model even in the presence of limited training data (82 pieces of data here). In the benchmark against mutants generated through the random mutagenesis approach (Figure 5f), RelaVDEP exhibits advantages in exploration efficacy across the fitness landscape, effectively identifying both high-fitness gain-of-function mutants (Supplementary Table 4) and low-fitness loss-of-function variants (Supplementary Table 6). Moreover, the top-performing variant derived through RelaVDEP achieves an increase of ∼141-fold in repression power over the wild-type protein, surpassing that obtained through random mutagenesis (∼84-fold) by 68% (Figure 5g).

Further inspection of the mutants sampled by RelaVDEP with high predicted fitness also reveals a conserved motif composed of seven mutations shared by a total of 15 decuple mutational variants (Supplementary Table 4). Taking this motif as the baseline, we analyzed the effects caused by the other variant-specific triple mutations (Figure 5h). Intriguingly, two distinct triple mutational groups (A53T-G86R-H117R and P49L-H117R-D119N) exhibit comparable functional gains over the base motif (261.5 vs. 261.8), whereas two similar variants (D12V-A53T-P94L and D12V-A53T-G86R) demonstrate marked phenotypic divergence (245.0 vs. 109.9) attributable to variation in a single site. This observation indicates the presence of position-dependent epistatic effects in the sequence evolution of AmeR. Finally, the fitness landscape of AmeR presents multiple discrete sequence islands of high predicted fitness in the 2D t-SNE subspace (Figure 5i), demonstrating that RelaVDEP can achieve functional improvement through divergent evolutionary trajectories from the same origin (*i.e.* wild-type).

### Improving enzyme activity through RelaVDEP

Polyethylene terephthalate (PET), a polymer of the polyester family, has emerged as a predominant plastic material. Over decades, substantial research efforts have focused on identifying and isolating PETases as well as engineering their catalytic activity and thermostability for efficient PET degradation under industrial conditions. Suliman et al.^45^ discovered a leaf-branch compost cutinase (LCC) exhibiting exceptional thermostability and high-temperature hydrolytic activity. Subsequent protein engineering by Tournier et al.^46^ yielded an optimized quadruple mutational variant (F243I-D238C-S283C-Y127G) named LCC-ICCG, which is currently recognized as one of the most efficient PETases. As a highly engineered variant with near optimal performance, this enzyme represents an even greater challenge for further improvement in protein engineering.

Informed by the general esterolytic properties of LCC-ICCG (Figure 6a, left), we employed RelaVDEP to facilitate the evolution of this enzyme, aiming for optimizing its hydrolytic activity towards a non-canonical small-molecule ester substrate, *p*-nitrophenol acetate (*p*NP-C2, abbreviated as *p*NP, see Supplementary Figure 5a), without sacrificing its thermostability. In addition, given the absence of prior activity data for *p*NP hydrolysis by LCC, we strategically designated all residues within a 12 Å radius of the catalytic triad (H207/D175/S130) for PET depolymerization as mutation sites permissible in the virtual directed evolution (Figure 6a, right).

**Figure 6.**
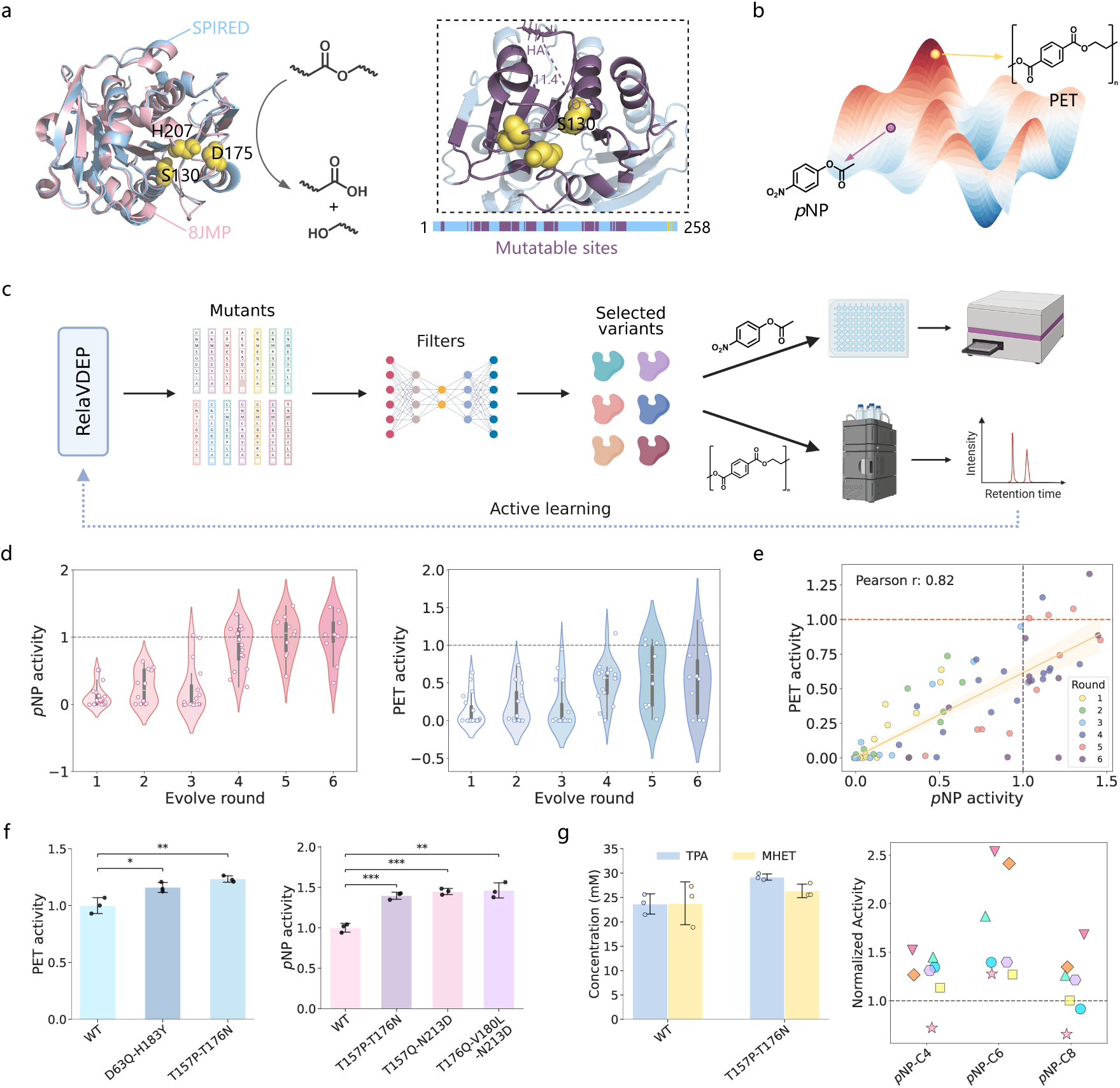
Optimization of enzyme activity for LCC-ICCG. **a)** Structural illustration of LCC-ICCG. The left panel compares the SPIRED-predicted protein structure (blue) against that determined experimentally (pink), with the three residues critical to ester hydrolysis depicted as yellow spheres. The right panel illustrates RelaVDEP-identified mutable sites (purple), encompassing all residues within a radius of 12 Å from the active site (composed of the three key residues). **b)** Hypothetical functional activity landscape of LCC-ICCG. The yellow dot denotes hydrolytic activity toward the macromolecular substrate PET, while the purple marker quantifies hydrolysis capability for the non-canonical small-molecule substrate *p*NP. **c)** Schematic illustration of the strategy for improving activity and thermostability for the LCC-ICCG enzyme with RelaVDEP. In this iterative approach, the reward model is continuously fine-tuned on the data of experimental feedback, improving its guiding capacity for the RL-assisted sequence exploration. **d)** Performance trajectory of LCC-ICCG through six rounds of RelaVDEP-based sequence evolution, with the activity evaluated for *p*NP hydrolysis (left) and PET depolymerization (right), respectively. **e)** Correlation analysis between PET and *p*NP hydrolytic activities for LCC-ICCG mutants derived from six rounds of RelaVDEP-based sequence evolution. The activities for PET depolymerization (vertical) and *p*NP hydrolysis (horizontal) have been normalized by the corresponding values of the wild-type LCC-ICCG (dashed lines). **f)** Comparison of variants with marked improvement in their catalytic activities for PET depolymerization (left) and *p*NP hydrolysis (right) against the wild-type (WT) LCC-ICCG (*n* = 3 repeats). “*”, “**” and “***” indicate increasing levels of statistical significance in the Student’s t test, corresponding to *p*-value < 0.05, 0.01 and 0.001, respectively. **g)** The left panel compares the proportions of TPA and MHET in the products of PET depolymerization catalyzed by the T157P-T176N variant and the wild-type (WT) LCC-ICCG (*n* = 3 repeats). The right panel shows the catalytic results of identified variants (each labeled by the same symbol and the same color) on *p*NP esters of varying lengths, where values in the vertical axis have been normalized to the wild-type (dashed line). In the formula “*p*NP-CX”, X refers to the number of carbons in the acyl group. Detailed data are available in Supplementary Tables 23, 24 & 25.

Here, we propose the concept of the general activity landscape of LCC-ICCG: this enzyme maintains generalized esterase activity, approaching a well-optimized peak performance for PET but preserving only a suboptimal catalytic power for atypical small-molecule substrates such as *p*NP (Figure 6b). Based on this perspective, we adopted an activelearning-like approach, integrating RelaVDEP-based sequence exploration and experimental validation to iteratively optimize *p*NP hydrolytic activity and thermostability for LCC-ICCG (Figure 6c). Considering the plausible correlation between the hydrolysis of PET and *p*NP, we adopted data on PET depolymerization activity and thermostability of two PETase homologs collected in a previous research^47^ to initiate supervised fine-tuning of the reward model, using it to coarsely guide the first round of enzyme evolution. In the subsequent three iterations, the experimental evaluation of *p*NP activity and thermostability on variants suggested by the previous rounds of virtual directed evolution enabled continual fine-tuning of the reward model, gradually steering RelaVDEP-based sequence exploration towards desired directions. Starting from the fifth round, PET depolymerization activity was also included in the experimental feedback for reward model fine-tuning, with the purpose of finding an LCC-ICCG variant having improved activity in both *p*NP hydrolysis and PET depolymerization as well as acceptable thermostability (*i.e. T_m_* > 65 ^◦^C, the optimal temperature for PET depolymerization).

Consistent with the variation of experimental schemes across iterations, the number of MLP layers in the reward model also changes dynamically (Supplementary Figure 6). Assisted by the continually improved reward model, the variants recommended by RelaVDEP show gradually elevated activities on *p*NP hydrolysis along the iterative evolution (Figure 6d, left). In addition, the catalytic performance on PET depolymerization begins to increase in the fourth round (Figure 6d, right). These trends could be observed more clearly in the correlation plot (Figure 6e). Noticeably, our method has shown the capacity to identify LCC-ICCG variants with simultaneous improvements in *p*NP hydrolysis and PET depolymerization since the fourth round.

Eventually, we obtained a number of LCC-ICCG mutants with evident amelioration in the hydrolytic activities of both *p*NP and PET (Figure 6f) as well as satisfactory thermostability (Supplementary Table 22) at the same time. Notably, most of these variants involve mutations at sites that have not been reported in previous studies (*e.g.*, D63, T157, T176, V180, H183, etc.), which reinforces that our method achieves significant functional improvement by exploring previously unvisited sequence subspaces. The most prominent mutant among these newly identified LCC-ICCG variants is a sequence carrying the double mutation of T157P-T176N (Supplementary Tables 15 & 21), which improves the catalytic activity by 23.4% for PET depolymerization and by 39.7% for *p*NP hydrolysis, respectively, at the cost of moderately reduced thermostability (*T_m_* = 80 ^◦^C). In the analysis of product components (Supplementary Figure 5b), the increased ratio of terephthalic acid (TPA, final product) to mono-(2-hydroxyethyl) terephthalate (MHET, intermediate product) within the PET hydrolytic products of this variant confirms the positive contribution of the corresponding double mutation in improving the depolymerization efficiency of LCC-ICCG (Figure 6g, left). Evaluation of *p*NP hydrolysis on *p*NP esters with increasing lengths in the acyl group (*e.g.*, *p*NP-C4, *p*NP-C6 and *p*NP-C8, see Supplementary Figure 5c) also supports the robust performance of identified enzyme variants (Figure 6g, right). In addition to T157P-T176N, we found two interesting double mutational variants with enhanced activity and acceptable thermostability. The variant A174K-N204R (Supplementary Tables 14 & 20) exhibits increases of 7.8% and 30.1% in the hydrolytic activities of PET and *p*NP, respectively, and a well-sustained thermostability (*T_m_* = 91 ^◦^C). The variant D63Q-H183Y (Supplementary Tables 13 & 19) presents a balanced improvement in both activities (16.0% for PET and 11.7% for *p*NP) as well as an acceptable *T_m_* of 74 ^◦^C. Overall, the LCC-ICCG variants identified in this study provide alternative starting points for the future engineering of PETases.

### Time consumption of RelaVDEP

The RelaVDEP pipeline is computationally efficient, attributable to our specific design in the distributed framework and parallel computing (Supplementary Figure 7). In Supplementary Table 26, we summarize the time consumption for RelaVDEP-based virtual directed evolution (including reward model optimization, RL-assisted sequence space exploration, and mutant library construction) on four NVIDIA RTX 2080Ti (24GB) GPUs in all the optimization projects mentioned above. Clearly, a single round of RelaVDEP typically costs no more than 2 days, which is acceptable considering the significant amounts of time saved in the lengthy process of generic wet-lab experimental verification in biology.

## Discussion

In this study, we have developed a virtual directed evolution framework, RelaVDEP, which adopts RL to facilitate the optimization and engineering of functional proteins. As the central component of an RL framework, the reward model should, in principle, have an ultra-fast inference speed to allow repeated function call in the RL system and a robust performance to closely align with the optimization objectives, two characteristics that can hardly be satisfied simultaneously in previous RL-based sequence optimization attempts. In RelaVDEP, we provide a solution by leveraging the high-throughput SPIRED-Fitness model that has been pre-trained on a plethora of DMS data to decipher the general protein sequence-fitness relationship. After architecture modification and model fine-tuning with specific DMS/experimental data, this model can provide instant and reliable predictions for the multiple mutational effects on specified protein properties, effectively guiding the agent’s learning trajectory in the RL-assisted virtual directed evolution. On the other hand, the efficiency of protein sequence space exploration is obviously determined by the underlying RL algorithms. Inspired by MuZero^28^, we design a model-based RL framework with MCTS incorporated. Unlike the policy optimization methods (*e.g.*, PPO^48^ and TRPO^49^) that rely on substantial real-time agent-environment interaction data for direct policy updating, MCTS systematically explores and evaluates a large number of diverse paths, effectively integrating statistics from simulated future trajectories. Within the RelaVDEP framework, MCTS simulations are particularly conducive to revealing the correlations between mutations occurring at different sites, by sampling the combinatorial mutational events from complete mutation trajectories. Furthermore, by leveraging a learnable neural network model to emulate MCTS planning, the hybrid strategy in RelaVDEP remarkably alleviates the computational overhead associated with the intensive simulation inherent in traditional MCTS algorithms, achieving remarkable enhancement of computational efficiency while preserving the exploratory advantages of MCTS.

Attributed to these designs, RelaVDEP excels in identifying variants with marked improvements in desired properties by efficiently exploring the combinatorial space of permissible mutations. When abundant DMS/experimental data are available to ensure sufficient training of the reward model (*e.g.*, in the cases of GFP, NUDT15 and VKOR1), or when the reward model presents a satisfactory prediction power after fine-tuning on the limited available data (*e.g.*, in the case of AmeR), a single round of RelaVDEP is typically sufficient for identifying well-performing variants, with evident advantages over the random mutagenesis approach. In the tasks of optimizing previously unexplored properties (*e.g.*, in the case of LCC-ICCG), the lack of experimental data prohibits reliable training of the reward model, and an active learning approach could be adopted to enable iterative fine-tuning of the reward model based on continued accumulation of experimental feedback. In such scenarios, RL is still superior to the conventional random generation approach (*e.g.*, adopted by ALDE^17^ and EVOLVEpro^24^) in recommending candidate variant sequences within the vast combinatorial space for model evaluation and experimental validation. Accordingly, by combining RelaVDEP with the active learning strategy, we successfully optimized the sequence of the PETase LCC-ICCG, acquiring previously unknown variants with marked improvements in the hydrolytic activity as well as acceptable thermostability.

In addition to the above achievements, due to the restriction on the maximum mutation count, a single round of RelaVDEP is constrained to explore the local sequence subspace around the starting sequence. In order to achieve remarkable sequence variation from the origin, iterative rounds of RelaVDEP are frequently required, which is costly and time-consuming. To address this problem, sequence generative models like ProteinMPNN^50^ or multimodal protein language models like ESM-3^51^ could be employed to generate a number of highly diversified sequences that are compatible with the structure or function of the target protein. After rapid experimental verification, multiple RelaVDEP trajectories could be initiated in parallel from these different valid seed sequences to explore the protein sequence subspaces located distantly from the starting sequence.

## Methods

### Graph representation of protein sequence state

Unlike the one-hot encoding used in traditional directed evolution tasks, the advancement in protein language models and protein structure prediction methods allows for an improved graph-based representation of the target protein, with nodes representing residues and edges describing inter-residue interactions. Before initializing the RL environment, we utilize the single-sequence-based structure predictor SPIRED^14^ to derive the protein structure given the initial sequence and extract node and edge features from it in a similar manner to the graph vector perceptron (GVP)^52^. Specifically, for a residue node, the vectors that point from its C*_α_* atom toward the C*_α_* atoms of its two neighboring residues are taken as the vector feature of this node. To derive the edge features, for each center residue of index *i*, we first select 12 spatially closest residues as its neighbors based on the inter-residue C*_α_* distances. The sequence separation and spatial distance between the neighboring residue *j* and the center residue *i* are taken as the scalar feature of the directional edge between the nodes *i* and *j*, while the unit vector pointing from the C*_α_* atom of residue *i* toward that of residue *j* is taken as the vector feature. Additionally, we integrate residue-wise embeddings of the ESM-2^7^ model as scalar features of the nodes. In general directed evolution tasks, a small number of residue mutations are not supposed to significantly alter the structure backbone of the target protein. Hence, at each mutation step, the features extracted from the predicted protein structure remain unchanged in the overall process, while the node scalar features are updated based on ESM-2 embeddings of the mutated sequence. The graph composed of nodes and edges is used to represent the observation output of the environment.

### Design of the mutation action

In the virtual directed evolution environment, the relationship between each action and the corresponding mutational information is:

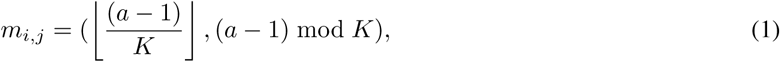

where *m_i,j_* is a tuple indicating that the residue at the *i*-th position will be mutated (by the first term) and that the mutation is directed towards the *j*-th amino acid type in the alphabet (by the second term), *a* is an integer defining the mutation action at the current step, *K* denotes the size of the amino acid alphabet, while ⌊·⌋ and mod refer to the floor and modulo operations, respectively. *a* = 0 indicates that no mutation occurs at the current step.

### Reward model

In most RL tasks, the environment generally provides immediate reward feedback in response to a received action signal. For the directed evolution task, the true reward signal should, in principle, be the real change in the desired property of the mutated protein obtained by experimental measurement. This approach is, however, impractical because of the labor-intensive and time-consuming nature of biological experiments. As an alternative, we choose SPIRED-Fitness^14^, a fast and reliable fitness predictor, as a substitute for the experimental measurement. Pre-trained on 2.5 million DMS data, SPIRED-Fitness performs well on the prediction of a variety of fitness metrics caused by single and double mutations in the zero-shot setting. In this work, we provide a number of models that are initialized with the pre-trained SPIRED-Fitness parameters but are tailored for varying scales of labeled data and distinct prediction tasks. In addition, we designate 5,000 as the recommended threshold: for datasets below this scale, the model undergoes supervised fine-tuning following the procedure outlined in Supplementary Table 27; otherwise, the model fine-tuning follows the procedure specified in Supplementary Table 28.

The Spearman correlation coefficient^53^ *ρ* is usually used to quantify the monotonic relationship between predicted fitness changes and true DMS/experimental labels. This metric is defined as the Pearson correlation coefficient between the rankings of two variables *X* and *Y*:

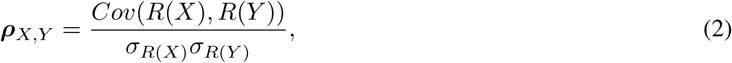

where *Cov* and *σ* denote covariance and standard deviation, respectively, while *R* is the ranking operation.

During training and fine-tuning of our reward models, following SPIRED-Fitness^14^, we adopt the soft Spearman loss^54^ as a differentiable substitute for the Spearman correlation coefficient to constrain the network parameters through gradient descent. In the last step, we employ the MSE loss as an additional constraint to optimize *α* and *β*, two parameters in the last linear layer connecting the difference of MLP predictions on wild-type and mutant sequences and the final output on the fitness change.

### MCTS

During the sampling process in the RL framework, MCTS simulations are performed before each step to select the action to be executed. In RelaVDEP, each node in the search tree corresponds to the graph representation of a protein sequence and stores information that includes the prior probability *P*, the visit count *N*, the average value *Q*, and the reward *R*. The edges connecting node pairs represent state transitions resulting from the action execution, corresponding to mutations in the sequence within the environment. The search process includes three main phases: selection, expansion, and backup.

### Selection

Each simulation starts at the root node *s*_0_, which represents the initial observation of the starting sequence. Following the Upper Confidence Bound (UCB) formula, at the step *k*, an action *a_k_* is selected from the set of legal actions, converting the system from the current node *s_k_*_−1_ to a child node *s_k_*:

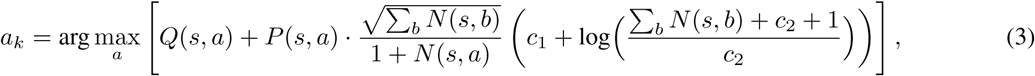

where the value *Q*, the prior *P* and the count *N* are all functions of the state *s* and the action *a*, while *c*_1_ and *c*_2_ are hyperparameters. This process is repeated until an unexpanded leaf node *s_l_* is reached.

In the directed evolution process, each residue has 19 possible mutation choices. Even with restrictions on the number of available positions *L* and the mutation depth *D*, the number of combinations of possible mutants is enormous. Additionally, the number of allowable actions per step can reach thousands ((*L* − *D_k_*) × 19, where *D_k_* is the current depth in MCTS), resulting in a highly sparse probability distribution and an evident disparity between *P* (*s, a*) and *Q*(*s, a*), making it challenging to balance the average reward *R* and the visit count *N* for each node with the default hyperparameters. To address this problem, we set 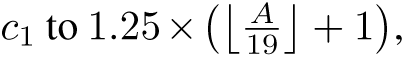 where *A* represents the size of the available action space and ⌊·⌋ denotes the floor function.

### Expansion

When the simulation process reaches a temporary leaf node *s_l_* whose current depth is less than the maximum mutation depth, the network performs a prediction for this state. The dynamics function *g* updates the hidden state *s_l_* of this leaf node and reports the reward *r_l_* for the transition from parent to child. The prediction function *f* estimates the policy *p_l_* and the value *v_l_*, which are stored at the node. Subsequently, the leaf node is expanded based on the set of available actions and the policy *p_l_*. Each new child node is initialized with the information set: {*P* (*s_l_, a*) = *p_l_*, *N* (*s_l_, a*) = 0, *Q*(*s_l_, a*) = 0}. To improve the efficiency of the MCTS, we update available actions by recording the positions of mutated residues along the search path to prevent redundant exploration of residues that have already been mutated.

### Backup

At the end of the simulation, the average value *Q*(*s, a*) and the visit count *N* (*s, a*) for all nodes along the search path are updated as follows:

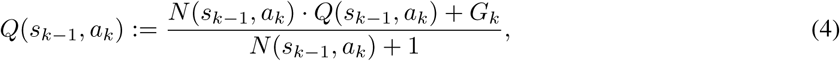

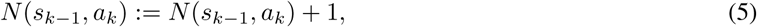

where *G* represents the cumulative discounted reward estimated comprehensively from the rewards (*r*) and values (*v*) as well as a discount (*γ*):

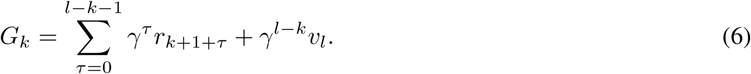

### Data generation

During MCTS simulations, the dynamics and prediction functions always make predictions with the latest network checkpoint. For each sampled mutation trajectory, 600 MCTS simulations are performed by default to select an action at each step. Here, we employ the same method as MuZero^28^ to parameterize the visit count distribution using a temperature parameter *T*:

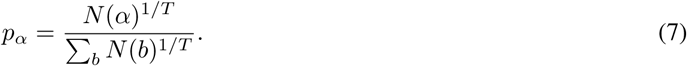

The temperature parameter *T*, initially set to 1.0, is gradually reduced during model training. Specifically, *T* is reduced to 0.5 after 5,000 steps and to 0.25 after 7,500 steps. This ensures an increased level of greediness for action selection as training progresses.

### Network architecture

Observations in many traditional RL environments, such as images or game boards, are typically represented using RGB frames or one-hot encodings. CNN or ResNet (*i.e.* residual convolution neural network) performs well on such problems. However, for inputs with complex topological structures taking the graph representations as in this study, we employ the GNN architecture. The network comprises three components: a representation function *h*, a dynamics function *g*, and a prediction function *f*.

### Representation function

The input of the representation function is the raw graph representation output of the RL environment (Algorithm 1). Initially, the node scalar features in the graph are projected from 1,280 dimensions to 128 dimensions through linear layers. These features are then concatenated with the one-hot encoding of the available action space. Subsequently, one-dimensional convolution is applied to map the features to 128 dimensions, which are then used to update the node scalar features of the graph representation. The features extracted from the predicted structure of the initial sequence remain unchanged during this process.

### Prediction function

The input of the prediction function is the hidden state (Algorithm 2). It will be transformed into an *L*-dimensional embedding through a global network, where *L* corresponds to the number of residues in the protein. The policy network and the value network in the prediction function receive this embedding as input, respectively. The policy network predicts the policy *p*, while the value network predicts the value *v*.

For reward and value predictions, we use an invertible transform 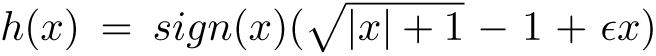 proposed in previous works^28,55^ to convert the discrete predicted outputs of the network into a single value. Additionally, the real reward and value obtained from sampling are also transformed using a similar transform *ϕ* to discretize the data dimension, which are then utilized to train the dynamics state network and the value network.

The dynamics state network, the policy network, and the value network employ the message-passing mechanism. In these networks, messages from neighboring nodes and edges are utilized to update node embeddings at each graph propagation step:

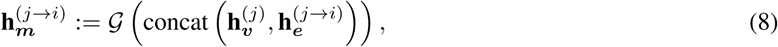

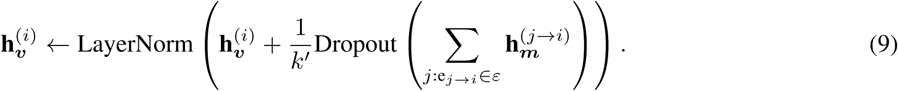

Here, G is a sequence of GVP modules^52^, which are designed to learn the vector-valued and scalar-valued functions for geometric vectors and scalars, respectively. In a GVP module, two linear projections are adopted for the transformation of scalar and vector features. Before transformation, scalar features are concatenated with the *L*_2_ norms of vector features, enabling the extraction of rotation-and translation-invariant information from the vector features. **h**_v_^(*i*)^ and **h**_e_^(*j*→*i*)^ are the embeddings of node *i* and directional edge *j_e_* → *i*, respectively, while **h_m_**^(*j*→*i*)^ represents the message passing from node *j* to node *i*. *k*^′^ is equal to the number of incoming messages *k* unless the protein contains fewer than *k* amino acid residues.

### Dynamics function

The input of the dynamics function includes the hidden state and the action to be executed in this state (Algorithm 3). The hidden state can be either the output of the representation function or the hidden state produced by the previous dynamics function. In the dynamics function, the node scalar features of the hidden state and the one-hot encoding of the action are concatenated, and these features undergo two types of transformation. First, the features are mapped to 128 dimensions via the dynamics state network. These transformed features are used to update the node scalar features in the input graph representation. The updated graph representation is then output as the next hidden state. Second, these features are used directly as node scalar features to update the graph representation. The updated graph representation is then fed to the dynamics reward network, which produces an output representing the predicted reward.

### RelaVDEP training

Training is performed on data in the continuously updated replay buffer (Algorithm 4). Different play trajectories have distinct sampling priority probabilities: 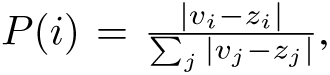 where *v_i_* represents the value obtained from the current step of MCTS, and *z_i_* denotes the *i*-th step return observed in the actual environment. To correct for sampling biases introduced by the prioritized sampling, we use an importance sampling ratio 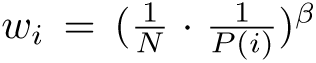 to adjust the loss.

During training, the network is unrolled for *K* virtual steps and aligned to the sampled trajectories. For each observed state in the trajectory, there is a corresponding actual evaluation value *z_t_*, a reward *u_t_*provided by the reward model, and a policy *π_t_* obtained through MCTS, where *π_t_* has a dimension matching the size of the available action space for that state. The network also predicts the evaluation value *v_t_^k^*, reward *r_t_^k^* and policy *p_t_^k^* for each unrolled step *k*.

Consequently, the total loss is computed as

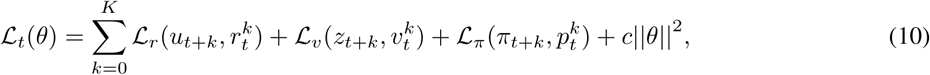

where L*_r_*, L*_v_* and L*_π_* represent the reward loss, the value loss and the policy loss, respectively, and *θ* refers to learnable parameters. In addition, the losses for the unrolled virtual step *k* are adjusted by a factor of 1*/k*.

Moreover, we employ the MuZero Reanalyze strategy^28^ to revisit and re-evaluate play trajectories in the replay buffer. By maintaining the states along the trajectories, we perform another MCTS using the latest network parameters. This process will update the policy and sampling priority probabilities, thereby continually improving the efficiency of sampling from the replay buffer. To improve computational efficiency during training, we utilize Ray^56^ to construct a distributed framework for the entire model, enabling concurrent processing across different threads for sampling, training, replay buffer management, reward prediction, and MuZero Reanalyze. The information transmission among these modules in the distributed framework is shown in Supplementary Figure 7. Specifically, the overall framework engages four NVIDIA RTX 2080Ti (24GB) GPUs, with two GPUs allocated to the sampling process (by six parallel workers), one GPU assigned to the reward model, and one GPU used for the training along with MuZero Reanalyze.

### Initialization and optimization of the mutant library

The workflow (Algorithm 7) begins with the determination of an optimal size *n* of the mutant library (10 ≤ *n* ≤ *N*, where *N* represents the total number of mutants sampled by RelaVDEP). Subsequently, variants with predicted fitness exceeding the wild-type level are selected and clustered based on their DHR^32^ sequence embeddings, with the number of clusters *K* (4 ≤ *K* ≤ 10) determined by systematically evaluating the Silhouette coefficients of candidate clusters. Initialization of the mutant library then proceeds through two sequential phases: 1) The highest-fitness unselected variant from each cluster is iteratively recruited until the vacant positions are fewer than *K*; 2) The remaining positions are filled sequentially by selecting the top-performing variants from the clusters sorted in descending order based on their cluster-averaged predicted fitness values, until the pre-determined library capacity is reached.

Subsequent optimization of this initialized mutant library is formulated as a multi-objective optimization framework designed to simultaneously address fitness and diversity:

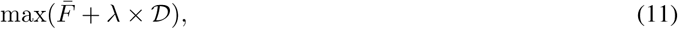

where 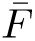 represents the average fitness of the current mutant library, D refers to the sequence diversity of the mutant library, and the hyperparameter *λ* is used to strike a balance between prioritizing high-fitness mutants and replacing existing sequences to enhance diversity. The diversity D is calculated as:

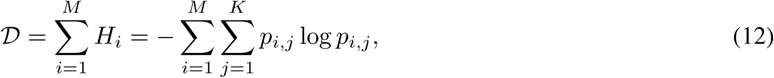

where *M* denotes the number of mutated positions across *L* (*i.e.* sequence length) residue sites observed after aligning all sequences in the mutant library, *H_i_* stands for the information entropy at the *i*-th position, *K* represents the size of the amino acid alphabet, and *p_i,j_* indicates the probability that the residue at the *i*-th position mutates to the *j*-th type of amino acid in the alphabet. Given that the choice of *λ* influences the balance between mean fitness and diversity in the mutant library, this work optimizes its value through parallel computation by systematically evaluating potential optimal libraries constructed with *λ* values increasing from 0.01 to 1.0 in steps of 0.01.

### Preference of mutation sites

The optimized variants recommended by RelaVDEP usually exhibit a notable positional preference over mutational sites, which is quantified as follows:

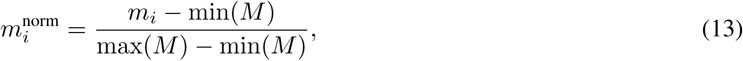

where *m*_i_^norm^ denotes the normalized mutational propensity of the residue at the *i*-th position, *m_i_*represents the raw mutational count at the *i*-th position across all variants within the optimized library, and *M* = {*m_i_* | *i* = 1, 2*,…, L*} refers to the set of raw mutational counts at all individual positions.

### RelaVDEP-guided evolution for GFP

#### Dataset

In order to fine-tune the reward model, we employed the DMS dataset from ProteinGym^13^, which includes the fluorescence intensity data of 51,714 avGFP mutants. Compared to wild-type avGFP, the starting sequence in this dataset has an F64L mutation. This mutation has been reported to improve protein folding stability at 37 ^◦^C^57^. The starting sequence is shown in Supplementary Information 2.3.

#### Supervised fine-tuning of the reward model

Following the procedure outlined in Supplementary Table 28, the DMS dataset was split into a training set and a validation set. Supervised fine-tuning was performed on the GAT module initialized with pre-trained weights from SPIRE-Fitness^14^ and a single-layer MLP with randomly initialized weights. Details regarding the composition of the training and validation sets are provided in the Source Data.

#### Verification of mutants

Mutant GFP fragments were cloned into a pET-28a plasmid using NEBuilder HiFi DNA Assembly. The constructed plasmids were transformed into BL21(DE3) competent cells and verified by Sanger sequencing. For flow cytometry analysis, BL21(DE3) colonies were cultured overnight, and the saturated culture was diluted into fresh medium and grown until OD600 reached 0.5, followed by induction with 0.1 or 0.5 mM IPTG at 16 ^◦^C for 12 hours. Cells were then resuspended and analyzed using a flow cytometer, with at least 10,000 events collected per sample. More details are shown in Supplementary Information 2.4.

#### Improving protein’s cellular abundance with RelaVDEP Dataset

We utilized two DMS datasets from ProteinGym^13^ to perform supervised fine-tuning of two distinctive reward models for the prediction of cellular abundance of NUDT15 and VKOR1 mutants, respectively. These datasets comprise 2,844 and 2,695 single mutational variants, respectively. The starting sequences of the two proteins are shown in Supplementary Information 2.3.

#### Supervised fine-tuning of the reward model

Following the procedure detailed in Supplementary Table 27, we first employed 5-fold cross-validation to determine the optimal layer number for the downstream MLP. Subsequently, the aforementioned dataset was partitioned into a training set and a validation set. We froze the weights of the GAT module, which was initialized with pre-trained weights from SPIRED-Fitness^14^, and performed supervised fine-tuning solely on the downstream MLP, which was randomly initialized. Details regarding the composition of the training and validation sets are provided in the Source Data.

#### Verification of mutants

NUDT15/VKOR1 and their mutants were synthesized and cloned into a pLJM1-eGFP plasmid. As an internal reference, mCherry was also cloned into another pLJM1 plasmid. Plasmids of pLJM1-NUDT15/VKOR1-eGFP and pLJM1-mCherry were then transfected into HEK293T cells using the transfection reagent PEI. At 48 hours after transfection, differences in cellular abundance were measured by flow cytometry using the ratio of eGFP to mCherry signals. More details are shown in Supplementary Information 2.5.

#### Tuning AmeR through RelaVDEP Dataset

We utilized the dataset reported in EvoAI^41^, which includes 82 mutational variants. The starting sequence is shown in Supplementary Information 2.3.

#### Supervised fine-tuning of the reward model

We first employed 10-fold cross-validation to determine the optimal number of MLP layers. Subsequently, the aforementioned dataset was partitioned into a training set and a validation set. The downstream MLP was fine-tuned using the soft Spearman loss, while the parameters *α* and *β* (of the last linear layer) were optimized using the MSE loss. Details of the training-validation partitioning scheme are provided in the Source Data.

#### Verification of mutants

Synthetic AmeR mutant genes were cloned into a p15A-origin/kanamycin-resistant test plasmid using the ClonExpress assembly kit and transformed into *E. coli*. The sequence-verified mutant strains were cultured and diluted. Log-phase cultures were further diluted into fresh medium for incubation. Upon arrest of bacterial growth, fluorescent protein expression was quantified by flow cytometry. Fold repression was calculated as the ratio of corrected signals attained without and with IPTG induction. More details are shown in Supplementary Information 2.6.

#### Improving LCC-ICCG activity through RelaVDEP Dataset

The evolution of LCC-ICCG underwent six rounds of screening. The first round used data derived from previous reports and patents on PETase activity and thermostability of variants derived from two PETase homologs to coarsely tune the reward model. The subsequent five rounds of evolution utilized the data determined in this study. The starting sequence of LCC-ICCG is shown in Supplementary Information 2.3.

#### Supervised fine-tuning of the reward model

Throughout the six rounds of sequence evolution, different datasets and strategies were employed to tune the reward model in each round, as detailed below.

**Round 1**: Coarse-tuning phase. Data on activity (Supplementary Tables 29, 30 & 31) and thermostability (Supplementary Tables 32, 33 & 34) of two PETase homologs (PDB IDs: 4EB0 and 7EOA) collected from a prior work^47^ were utilized, with 20% of the 4EB0 data serving as the validation set and the remainder constituting the training set. Per-epoch training was conducted on data from individual target proteins to enhance the accuracy of the reward model in ranking fitness among variants of the same protein. The reward model for activity prediction was initialized with pre-trained weights inherited from SPIRED-Fitness^14^, while the thermostability prediction filter model inherited pre-trained weights from SPIRED-Stab^14^.

**Round 2**: Hybrid-tuning phase. All reported data formed the training set, while experimentally measured data from the first-round mutants (*p*NP hydrolysis activity and thermostability) generated in this work comprised the validation set. The training protocols for both the activity reward model and the thermostability filter model followed the same protocol as in Round 1.

**Round 3-6**: Precision-tuning phase. Reward models were fine-tuned using exclusively experimentally determined data from this study. Each round of tuning was performed from scratch according to the workflow specified in Supplementary Table 27. The activity prediction model served as the RelaVDEP reward model (weights initialized from SPIRED-Fitness), and the thermostability prediction model functioned as the filter (weights initialized from SPIRED-Stab).

#### Verification of mutants

All mutant gene sequences were amplified via polymerase chain reaction (PCR) and subsequently cloned into a pET15b vector using Gibson assembly. Recombinant proteins were expressed overnight at 16 ^◦^C, followed by cell lysis via sonication. The supernatant was subjected to Ni-NTA chromatography purification and buffer-exchanged into Tris buffer using a desalting column. Enzymatic activities of LCC-ICCG and its mutants were determined using two substrates: *p*NP and PET. The hydrolytic activity of *p*NP was measured by monitoring the absorbance change at 415 nm using a microplate reader, with enzyme activity calculated from the slope of the linear reaction phase. PET depolymerization assays were conducted at 65 ^◦^C, with reaction aliquots collected after 8 hours to quantify the release of TPA via ultra-performance liquid chromatography analysis. All mutant activities were normalized with respect to the reference enzyme LCC-ICCG. Thermostability was assessed by determining the melting temperature (*T_m_*) using Differential Scanning Fluorimetry (DSF, a fluorescence-based thermal shift assay) performed on a quantitative real-time PCR system. More details are shown in Supplementary Information 2.7.

## Data availability

Model parameters are available at https://zenodo.org/doi/10.5281/zenodo.15720582.

## Code availability

All codes of the RelaVDEP pipeline are available at https://github.com/Gonglab-THU/RelaVDEP. Datasets used for model training and fine-tuning are also available at the same repository.

## Author contribution

H. G. proposed the concept and theory. For the RelaVDEP, T. M. and H. G. proposed the initial model design, while T. M. implemented coding, model training and testing. In all sequence evolution projects, T. M. and N. X. performed the virtual sequence optimization, and H. G. supervised the process. For the GFP project, W. W. performed experimental verification, and G.-Q. C. supervised the process. For the NUDT15 and VKOR1 projects, J. Z. performed experimental verification, and L. C. supervised the process. For the AmeR project, Y. S. performed experimental verification, and S. Z. supervised the process. For the LCC-ICCG project, W.-B. Z. proposed the optimization plan, Y. W. performed experimental verification, and W.-B. Z. supervised the process. T. M. and N. X. performed data analysis and generated figures. T. M., N. X. and H. G. wrote the manuscript, and all the other authors made revisions. All authors agreed with the final manuscript.

## Acknowledgements

This work has been supported by the Ministry of Science and Technology of China (#2023YFF1204400 to H. G. and W.-B. Z.), by the National Natural Science Foundation of China (#32171243 to H. G. and #22331003 to W.-B. Z.) and by the Beijing Frontier Research Center for Biological Structure.

## Ethics declarations

### Competing interests

The authors declare no competing interests.

## 1 Supplementary Results

### 1.1 Results for the evolution of GFP

#### 1.1.1 Benchmarking the reward model trained with various proportions of data

**Supplementary Figure 1.**
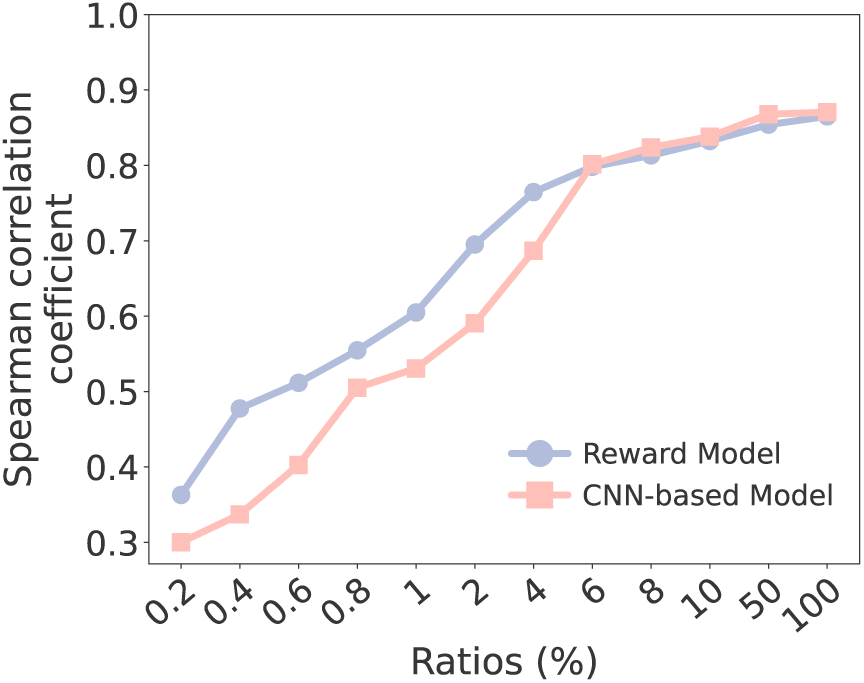
Comparison of fitness predictors. The horizontal axis represents the proportion of data in the training set used for model training, while the vertical axis stands for the Spearman correlation coefficient between model prediction and label in the validation set. Clearly, when the amount of training data is limited (*e.g.*, < 5%), the graph-based reward model proposed in this study (red) shows notable advantages over the CNN-based model adopted by EvoPlay (blue).

#### 1.1.2 Experimental verification

**Supplementary Table 1.**
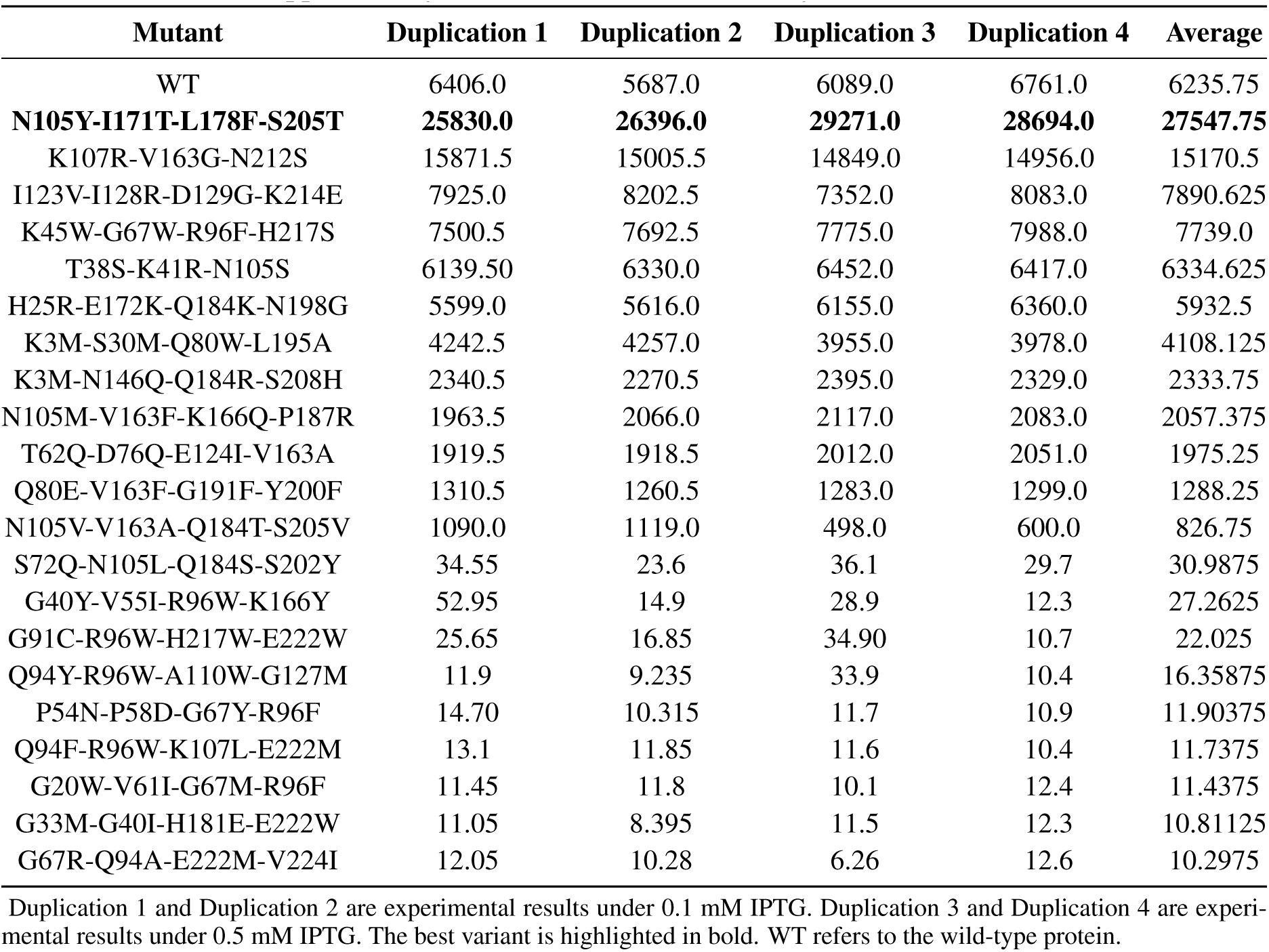
Fluorescence intensity of GFP mutants.

| Mutant | Duplication 1 | Duplication 2 | Duplication 3 | Duplication 4 | Average |
| --- | --- | --- | --- | --- | --- |
| WT | 6406.0 | 5687.0 | 6089.0 | 6761.0 | 6235.75 |
| **N105Y-I171T-L178F-S205T** | **25830.0** | **26396.0** | **29271.0** | **28694.0** | **27547.75** |
| K107R-V163G-N212S | 15871.5 | 15005.5 | 14849.0 | 14956.0 | 15170.5 |
| I123V-I128R-D129G-K214E | 7925.0 | 8202.5 | 7352.0 | 8083.0 | 7890.625 |
| K45W-G67W-R96F-H217S | 7500.5 | 7692.5 | 7775.0 | 7988.0 | 7739.0 |
| T38S-K41R-N105S | 6139.50 | 6330.0 | 6452.0 | 6417.0 | 6334.625 |
| H25R-E172K-Q184K-N198G | 5599.0 | 5616.0 | 6155.0 | 6360.0 | 5932.5 |
| K3M-S30M-Q80W-L195A | 4242.5 | 4257.0 | 3955.0 | 3978.0 | 4108.125 |
| K3M-N146Q-Q184R-S208H | 2340.5 | 2270.5 | 2395.0 | 2329.0 | 2333.75 |
| N105M-V163F-K166Q-P187R | 1963.5 | 2066.0 | 2117.0 | 2083.0 | 2057.375 |
| T62Q-D76Q-E124I-V163A | 1919.5 | 1918.5 | 2012.0 | 2051.0 | 1975.25 |
| Q80E-V163F-G191F-Y200F | 1310.5 | 1260.5 | 1283.0 | 1299.0 | 1288.25 |
| N105V-V163A-Q184T-S205V | 1090.0 | 1119.0 | 498.0 | 600.0 | 826.75 |
| S72Q-N105L-Q184S-S202Y | 34.55 | 23.6 | 36.1 | 29.7 | 30.9875 |
| G40Y-V55I-R96W-K166Y | 52.95 | 14.9 | 28.9 | 12.3 | 27.2625 |
| G91C-R96W-H217W-E222W | 25.65 | 16.85 | 34.90 | 10.7 | 22.025 |
| Q94Y-R96W-A110W-G127M | 11.9 | 9.235 | 33.9 | 10.4 | 16.35875 |
| P54N-P58D-G67Y-R96F | 14.70 | 10.315 | 11.7 | 10.9 | 11.90375 |
| Q94F-R96W-K107L-E222M | 13.1 | 11.85 | 11.6 | 10.4 | 11.7375 |
| G20W-V61I-G67M-R96F | 11.45 | 11.8 | 10.1 | 12.4 | 11.4375 |
| G33M-G40I-H181E-E222W | 11.05 | 8.395 | 11.5 | 12.3 | 10.81125 |
| G67R-Q94A-E222M-V224I | 12.05 | 10.28 | 6.26 | 12.6 | 10.2975 |

Duplication 1 and Duplication 2 are experimental results under 0.1 mM IPTG. Duplication 3 and Duplication 4 are experimental results under 0.5 mM IPTG. The best variant is highlighted in bold. WT refers to the wild-type protein.

#### 1.1.3 Analysis of the identified quadruple mutational variant

**Comparison with known mutations on avGFP** The best avGFP variant identified by the single round of RelaVDEP carries four mutations: N105Y, I171T, L178F and S205T. Upon investigating these mutations, we identified N105Y as the mutation present in a fast-folding mutant, demonstrating increased tolerance to denaturants in the refolding experiment^1^. S205T, located adjacent to the chromophore, is another well-known mutation recognized as contributing to folding enhancement^2^. The rest two mutations, I171T and L178F, were only observed in the DMS data with suboptimal fitness scores (*i.e.* fluorescence intensity). By incorporating all of these four mutations, the quadruple mutational variant identified in this work is undoubtedly novel, different from all variants reported in previous directed evolution and/or DMS studies. Furthermore, the synergistic effect of these four mutations is salient, leading to significantly improved fluorescence intensity over any subsets of them ever explored in prior works. This observation not only highlights RelaVDEP’s high exploration efficiency for the combinatorial optimization of individual beneficial mutations but also underscores the necessity of considering epistatic effects between mutations in protein engineering.

**Comparison with eGFP** We further compared this variant with eGFP (M1_S2insV/F64L/S65T/H231L), a widely used fluorescence dye that has been optimized through extensive protein engineering. Both constructs share the F64L mutation, which is known to improve folding stability^3^. However, eGFP also incorporates a sequence insertion (M1_S2insV) and a chromophore mutation (S65T) at Ser^65^. RelaVDEP’s design is constrained by the need for protein sequence alignment during fitness prediction, which means that its current version is unable to handle insertions or deletions. Moreover, to boost the success rate of our directed evolution task, we deliberately excluded the three chromophore residues as mutable sites, considering that their mutations frequently lead to disastrous results (*e.g.*, complete loss of fluorescence). While this *a priori* decision somewhat limited the maximum potential of our directed evolution, it served as a crucial proof-of-principle for more complex evolutionary challenges, particularly when the domain knowledge is lacking. For instance, our relatively mild strategy could be safely applied to practical tasks like the enzymatic optimization (*e.g.*, LCC-ICCG in this study), by first pinpointing active sites through substrate docking and then only mutating the surrounding residues. Although the fluorescence intensity of the best RelaVDEP-identified mutant is still lower than that of eGFP (Supplementary Figure 2), RelaVDEP has demonstrated the power of effectively accelerating the directed evolution process, in comparison to the serendipitous success on eGFP in the lengthy history of GFP engineering.

**Supplementary Figure 2.**
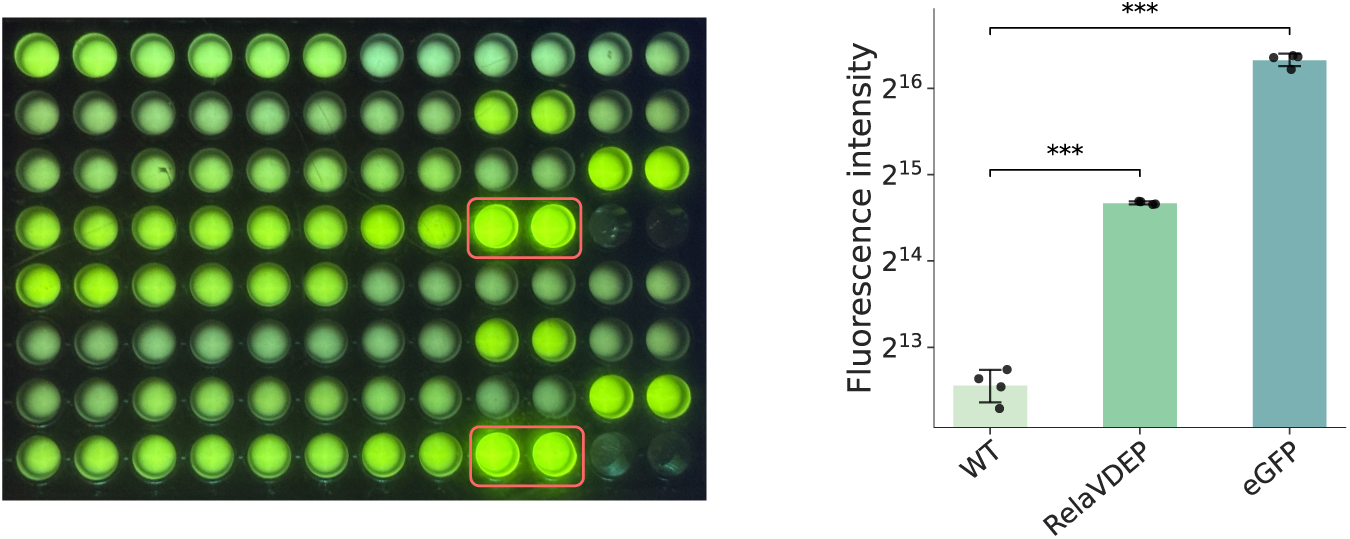
Experimental fluorescence intensity of RelaVDEP mutants and eGFP. The left panel displays the supplementary experiments for eGFP, indicated by the red box, while the right panel presents a comparison of the fluorescence intensity values for avGFP, the best RelaVDEP variant, and eGFP (*n* = 4 repeats). “***” indicates statistical significance in the Student’s t test, with *p*-value < 0.001.

### 1.2 Results for the optimization of protein’s cellular abundance

#### 1.2.1 Examining the mutational preference in cellular abundance optimization

**Supplementary Figure 3.**
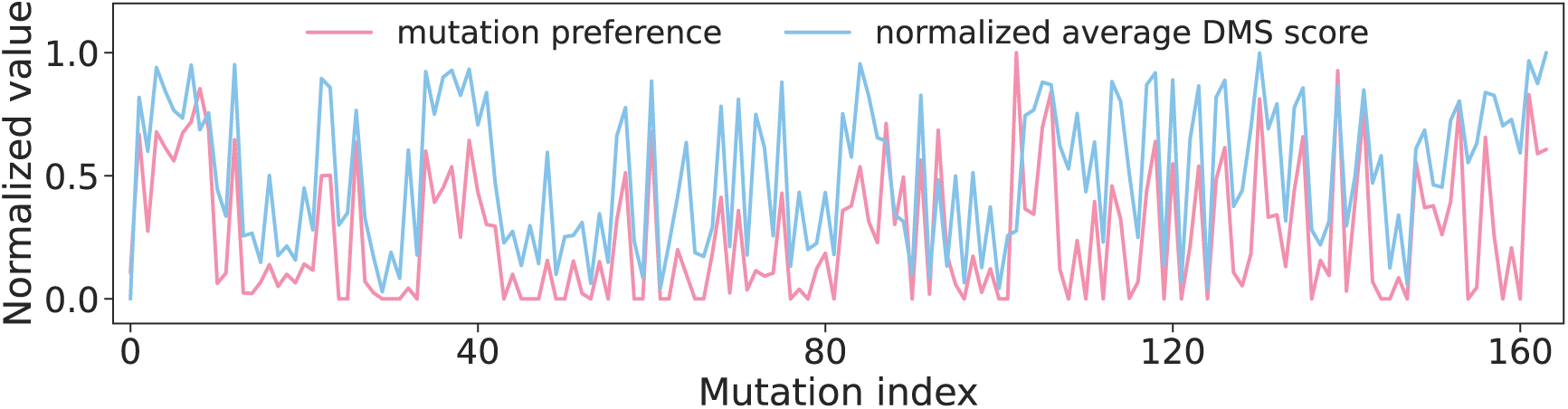
Mutational preference in the RelaVDEP-based evolution of NUDT15. Preferences of mutation sites summarized from the single-round RelaVDEP evolution (red) are compared with the positional abundance scores of single mutational variants in the DMS dataset (blue). The Spearman correlation coefficient between these two indicators is 0.8152.

**Supplementary Figure 4.**
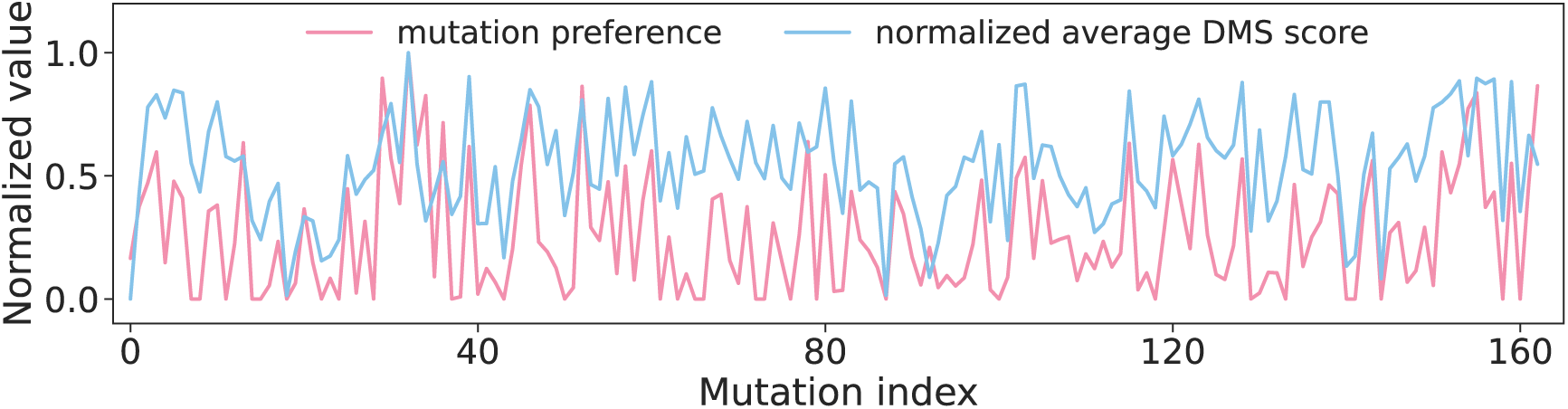
Mutational preference in the RelaVDEP-based evolution of VKOR1. Preferences of mutation sites summarized from the single-round RelaVDEP evolution (red) are compared with the positional abundance scores of single mutational variants in the DMS dataset (blue). The Spearman correlation coefficient between these two indicators is 0.6709.

#### 1.2.2 Experimental verification

**Supplementary Table 2.** Abundance of NUDT15 mutants.

| Mutant | Duplication 1 | Duplication 2 | Duplication 3 | Average |
| --- | --- | --- | --- | --- |
| WT | 1.2159 | 1.1977 | 1.1857 | 1.1998 |
| **S4P-V14I-Q61I** | **1.9761** | **1.9935** | **1.9840** | **1.9845** |
| T2A-K35A-L153F | 1.3464 | 1.3150 | 1.2808 | 1.3141 |
| G13T-A40I-E127N | 1.4312 | 1.4389 | 1.4429 | 1.43768 |
| S37C-Q61L-V103D | 1.5373 | 1.4734 | 1.5354 | 1.5154 |
| Q61A-V123W-P124N | 1.6438 | 1.6212 | 1.6383 | 1.6344 |
| G9I-T21I-E121Q | 1.0973 | 1.1104 | 1.1444 | 1.1174 |
| G9V-Q61R-Y90A | 1.6440 | 1.6522 | 1.6393 | 1.6452 |
| G9I-T21V-E88M | 1.2717 | 1.2577 | 1.2063 | 1.2453 |

All values in this table are experimentally measured eGFP:mCherry ratios. The best variant is highlighted in bold. WT refers to the wild-type protein.

**Supplementary Table 3.** Abundance of VKOR1 mutants.

| Mutant | Duplication 1 | Duplication 2 | Duplication 3 | Average |
| --- | --- | --- | --- | --- |
| WT | 0.3561 | 0.3463 | 0.3465 | 0.3496 |
| **R33D-R35Y-E155F** | **0.4608** | **0.4598** | **0.4447** | **0.4551** |
| R33C-R35T-L42M | 0.2938 | 0.2889 | 0.2786 | 0.2871 |
| K30D-A31Y-R33A | 0.3778 | 0.3754 | 0.3649 | 0.3727 |
| K30E-R35F-V104H | 0.2921 | 0.2880 | 0.2933 | 0.2911 |
| R35E-K152M-H163G | 0.3262 | 0.3178 | 0.3249 | 0.3230 |
| G6R-A14L-K30E | 0.3627 | 0.3676 | 0.3566 | 0.3623 |
| A14L-K30E-S144L | 0.3467 | 0.3319 | 0.3258 | 0.3348 |
| G6R-K30E-Y129H | 0.3292 | 0.3320 | 0.3315 | 0.3309 |

All values in this table are experimentally measured eGFP:mCherry ratios. The best variant is highlighted in bold. WT refers to the wild-type protein.

### 1.3 Results for the evolution of AmeR

#### 1.3.1 Variants identified by single-round RelaVDEP

**Supplementary Table 4.** Fold repression of AmeR mutants (RelaVDEP-Top)

| Mutant | Duplication 1 | Duplication 2 | Duplication 3 | Average |
| --- | --- | --- | --- | --- |
| **D33E-I34V-A51T-S57R-A75T-I80V-G83V-C93R-P94L-Q204R** | 98.6512 | 121.2000 | 111.6316 | 110.4942 |
| **D12V-D33E-I34V-A51T-A53T-S57R-A75T-C93R-P94L-Q204R** | 161.8400 | 404.6000 | 168.5833 | 245.0078 |
| **D12V-D33E-I34V-A51T-A53T-S57R-I80V-C93R-H117R-Q204R** | 86.6829 | 88.8500 | 86.6829 | 87.4053 |
| **D33E-I34V-A51T-A53T-S57R-A75T-I80V-C93R-H117R-A218T** | 116.6000 | 124.9286 | 174.9000 | 138.8095 |
| **D33E-I34V-A51T-S57R-A75T-G83V-C93R-P94L-H117R-Q204R** | 40.0875 | 61.6731 | 50.9048 | 50.8884 |
| **D33E-I34V-A51T-A53T-S57R-A75T-G86R-C93R-H117R-Q204R** | 190.5455 | 419.2000 | 174.6667 | 261.4707 |
| **D12V-D33E-I34V-A51T-A53T-S57R-A75T-C93R-H117R-Q204R** | 161.4583 | 129.1667 | 107.6389 | 132.7546 |
| **D33E-I34V-A36T-A51T-A53T-S57R-I80V-C93R-H117R-Q204R** | 157.5357 | 129.7353 | 133.6667 | 140.3126 |
| **D33E-I34V-A51T-A53T-S57R-A75T-C93R-H117R-Q204R-A218T** | 61.6182 | 80.6905 | 67.7800 | 70.0296 |
| D33E-I34V-A36T-A51T-A53T-S57R-A75T-I80V-C93R-H117R | 28.1308 | 55.7407 | 37.6250 | 40.4989 |
| D33E-I34V-A51T-S57R-I80V-G83V-C93R-P94L-H117R-Q204R | 49.3833 | 84.6571 | 84.6571 | 72.8992 |
| D33E-I34V-A51T-A53T-S57R-G86R-C93R-P94L-H117R-Q204R | 15.4178 | 18.8736 | 21.6053 | 18.6322 |
| **D33E-I34V-A51T-S57R-A75T-C93R-P94L-H117R-D119N-Q204R** | 121.0938 | 387.5000 | 276.7857 | 261.7932 |
| **D33E-I34V-A51T-S57R-A75T-I80V-C93R-H107Y-V188I-Q204R** | 106.0286 | 106.0286 | 97.6579 | 103.2383 |
| **D33E-I34V-A51T-A53T-S57R-A75T-I80V-C93R-K135R-Q204R** | 11.1932 | 14.9242 | 15.6349 | 13.9174 |
| D12V-D33E-I34V-A51T-A53T-S57R-C93R-P94L-H117R-Q204R | 33.4536 | 30.6132 | 33.4536 | 32.5068 |
| D33E-I34V-A36T-A51T-A53T-S57R-I80V-C93R-P94L-H117R | 54.0714 | 49.1558 | 66.4035 | 56.5436 |
| **D33E-I34V-A51T-S57R-A75T-I80V-G83V-P94L-H117R-Q204R** | 90.0513 | 159.6364 | 79.8182 | 109.8353 |
| **D33E-I34V-A51T-A53T-S57R-A75T-C93R-H107Y-H117R-Q204R** | 20.8811 | 28.4381 | 25.3051 | 24.8748 |
| **D12V-D33E-I34V-A51T-A53T-S57R-A75T-C93R-V188I-Q204R** | 37.1803 | 36.0000 | 31.5000 | 34.8934 |
| **D33E-I34V-A51T-S57R-A75T-G83V-C93R-P94L-Q204R-G212S** | 25.3931 | 34.7358 | 28.7656 | 29.6315 |
| **D12V-D33E-I34V-A51T-A53T-S57R-A75T-G86R-C93R-Q204R** | 10.1899 | 12.1970 | 13.4728 | 11.9532 |
| **D12V-D33E-I34V-A51T-A53T-S57R-A75T-C93R-V188I-Q204R** | 104.5294 | 96.0541 | 148.0833 | 116.2223 |
| **D12V-D33E-I34V-A51T-S57R-A75T-G86R-C93R-H117R-Q204R** | 28.7934 | 41.9759 | 38.7111 | 36.4935 |

Fold repression of AmeR mutants was quantified via flow cytometry. Calculated as: (background-subtracted MFI uninduced) / (background-subtracted MFI induced), where MFI denotes the median fluorescence intensity. For mutants exhibiting high-affinity promoter binding (*i.e.* high fold repression), induced MFI approaches background levels, resulting in elevated replicate variability (*i.e.* standard deviation). Mutants highlighted in bold share a motif composed of 7 common mutations: D33E, I34V, A51T, S57R, A75T, G83V, and Q204R.

**Supplementary Table 5.**
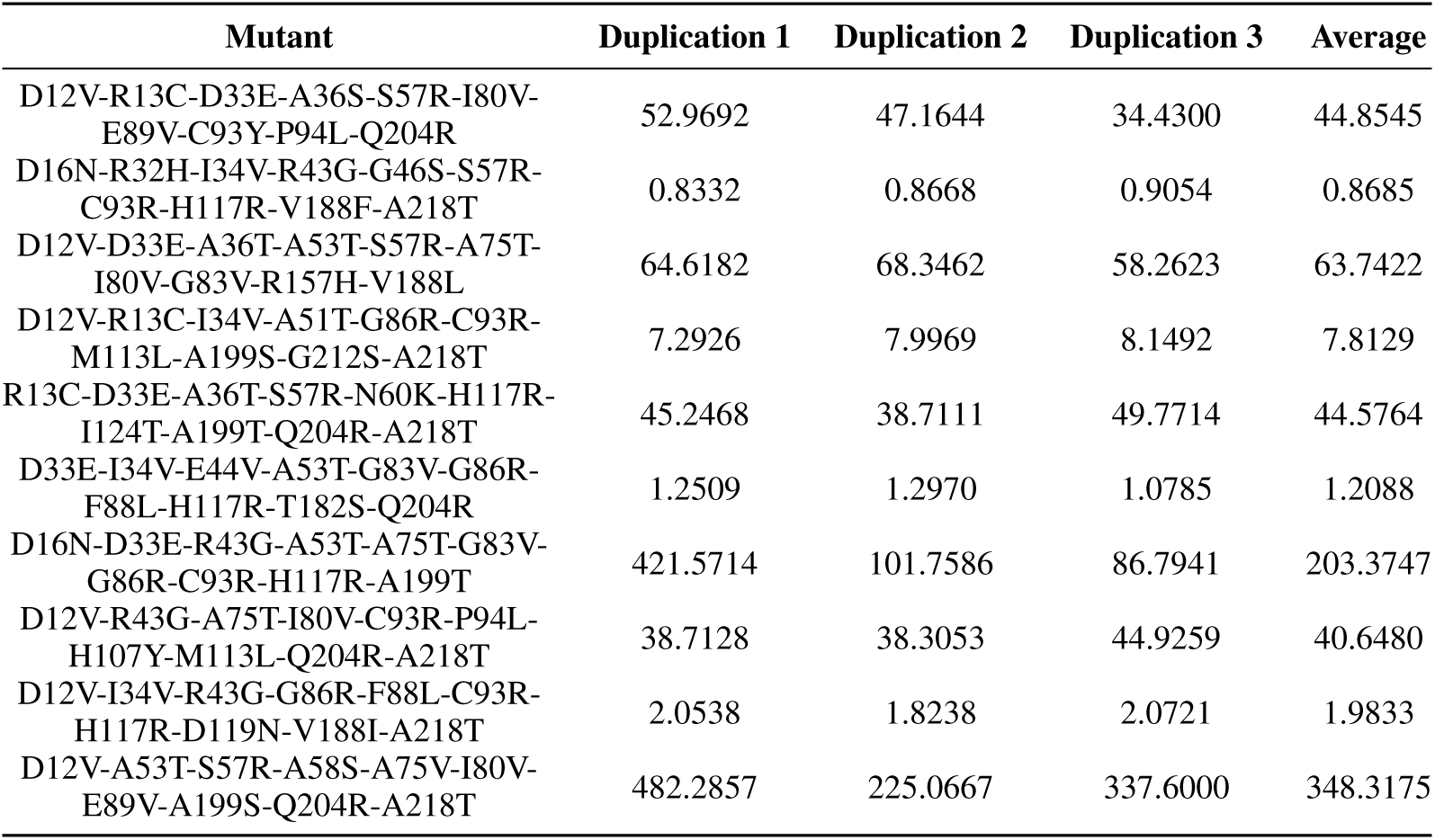
Fold repression of AmeR mutants (RelaVDEP-Mid)

**Supplementary Table 6.** Fold repression of AmeR mutants (RelaVDEP-Bot)

| Mutant | Duplication 1 | Duplication 2 | Duplication 3 | Average |
| --- | --- | --- | --- | --- |
| R13S-D16N-R32H-I34V-E44V-A58S-N60K-E89V-D119N-V188L | 1.0696 | 1.0365 | 1.0365 | 1.0475 |
| R13S-D16H-A36S-R43S-G83V-F88L-I124K-R125W-T182S-V206I | 34.0459 | 41.2333 | 37.4848 | 37.5880 |
| R32H-R43S-G83V-G86R-F88L-K135R-R157H-V188F-A199S-G212S | 4.9933 | 5.6234 | 4.6223 | 5.0797 |
| R13S-D16H-R32H-A36S-A75V-E89V-H107Y-T182S-V206I-A218T | 1.3019 | 1.4933 | 1.3284 | 1.3746 |
| D12V-R13C-D16N-R32H-R43S-G46S-F88L-H117R-R157H-T182S | 1.7935 | 1.9887 | 1.8611 | 1.8811 |
| R13C-R32H-R43G-D54H-H107Y-D119H-I124T-K135R-V188F-A199T | 10.8664 | 11.7121 | 12.1371 | 11.5719 |
| A36T-E44V-A53T-A58S-N60K-I80V-H107Y-K135R-R157H-V188F | 3.5081 | 3.9479 | 3.5526 | 3.6695 |
| R43S-G46S-D54H-A58S-G83V-P94L-R125W-R157H-T182S-V188L | 0.8632 | 0.7819 | 0.7517 | 0.7989 |
| R32H-A36T-A58S-G86R-C93Y-D119H-I124T-K135R-R157H-V188L | 3.9597 | 4.1929 | 3.9053 | 4.0193 |
| D16H-R32H-R43S-E44V-A75V-R125W-R157H-V188F-Q204H-V206I | 0.9959 | 0.9921 | 0.9685 | 0.9855 |

#### 1.3.2 Variants suggested by the random approach

**Supplementary Table 7.** Fold repression of AmeR mutants (Random-Top)

| Mutant | Duplication 1 | Duplication 2 | Duplication 3 | Average |
| --- | --- | --- | --- | --- |
| D33E-I34V-A51T-D54H-S57R-I80V-G83V-C93R-V188I-Q204R | 15.4524 | 19.0882 | 19.0882 | 17.8763 |
| D12V-I34V-A51T-A53T-S57R-A75T-I80V-P94L-H117R-G212S | 41.8193 | 61.9821 | 61.9821 | 55.2612 |
| D12V-I34V-G46S-A51T-S57R-I80V-C93R-Q204R-G212S-A218T | 31.8919 | 59.0000 | 49.1667 | 46.6862 |
| D33E-I34V-A51T-A53T-S57R-A75V-I80V-T182S-V188I-Q204R | 31.1359 | 35.6333 | 35.6333 | 34.1342 |
| D16N-I34V-A51T-A53T-S57R-A75T-I80V-G86R-C93R-K135R | 70.4386 | 108.5135 | 108.5135 | 95.8219 |
| D33E-I34V-R43S-A51T-A53T-S57R-G86R-C93R-A199S-Q204R | 138.4545 | 240.4737 | 240.4737 | 206.4673 |
| I34V-A51T-A53T-S57R-A75V-I80V-G86R-P94L-D119H-I124K | 115.1515 | 211.1111 | 211.1111 | 179.1246 |
| D16H-I34V-A51T-S57R-A75T-I80V-C93R-V188L-Q204R-G212S | 22.2128 | 23.5489 | 23.5489 | 23.1035 |
| D33E-I34V-A51T-A53T-S57R-I80V-P94L-M113L-D119H-A218T | 78.2292 | 85.3409 | 89.4048 | 84.3249 |
| D33E-I34V-A51T-S57R-I80V-F88L-H117R-D119N-K135R-Q204R | 35.8876 | 45.6286 | 45.6286 | 42.3816 |

**Supplementary Table 8.** Fold repression of AmeR mutants (Random-Mid)

| Mutant | Duplication 1 | Duplication 2 | Duplication 3 | Average |
| --- | --- | --- | --- | --- |
| D12V-N60K-A75V-G83V-G86R-C93Y-M113L-I124K-K135R-T182S | 21.3742 | 23.2267 | 20.4941 | 21.6983 |
| R13S-E44V-S57R-A58S-G83V-C93R-I124T-R125W-K135R-G212S | 52.6456 | 56.9726 | 55.4533 | 55.0238 |
| R13C-R43G-D54H-S57R-E89V-C93R-I124T-R157H-T182S-G212S | 1.1132 | 1.1087 | 0.9843 | 1.0687 |
| R43S-E44V-A51T-A53T-N60K-F88L-P94L-H117R-K135R-V188F | 31.6435 | 31.3707 | 31.3707 | 31.4616 |
| D12V-R13C-D16E-D33E-A36T-E89V-M113L-I124T-T182S-Q204H | 63.5833 | 84.7778 | 66.9298 | 71.7636 |
| R32H-D33E-E44V-A51T-E89V-H107Y-I124T-T182S-G212S-A218T | 44.5679 | 72.2000 | 76.8085 | 64.5255 |
| R32H-A36S-E44V-G46S-S57R-I80V-H107Y-M113L-V188I-G212S | 0.8806 | 0.8909 | 0.8038 | 0.8584 |
| R43G-S57R-A58S-G83V-F88L-P94L-M113L-K135R-V188F-A199S | 27.8760 | 30.4746 | 33.2963 | 30.5489 |
| D12V-D16N-A51T-D54H-A58S-A75V-F88L-I124T-T182S-G212S | 10.9080 | 10.7841 | 11.4798 | 11.0573 |
| R32H-A36S-R43S-A51T-A53T-F88L-H117R-T182S-G212S-A218T | 5.3247 | 6.3077 | 5.7400 | 5.7908 |

**Supplementary Table 9.**
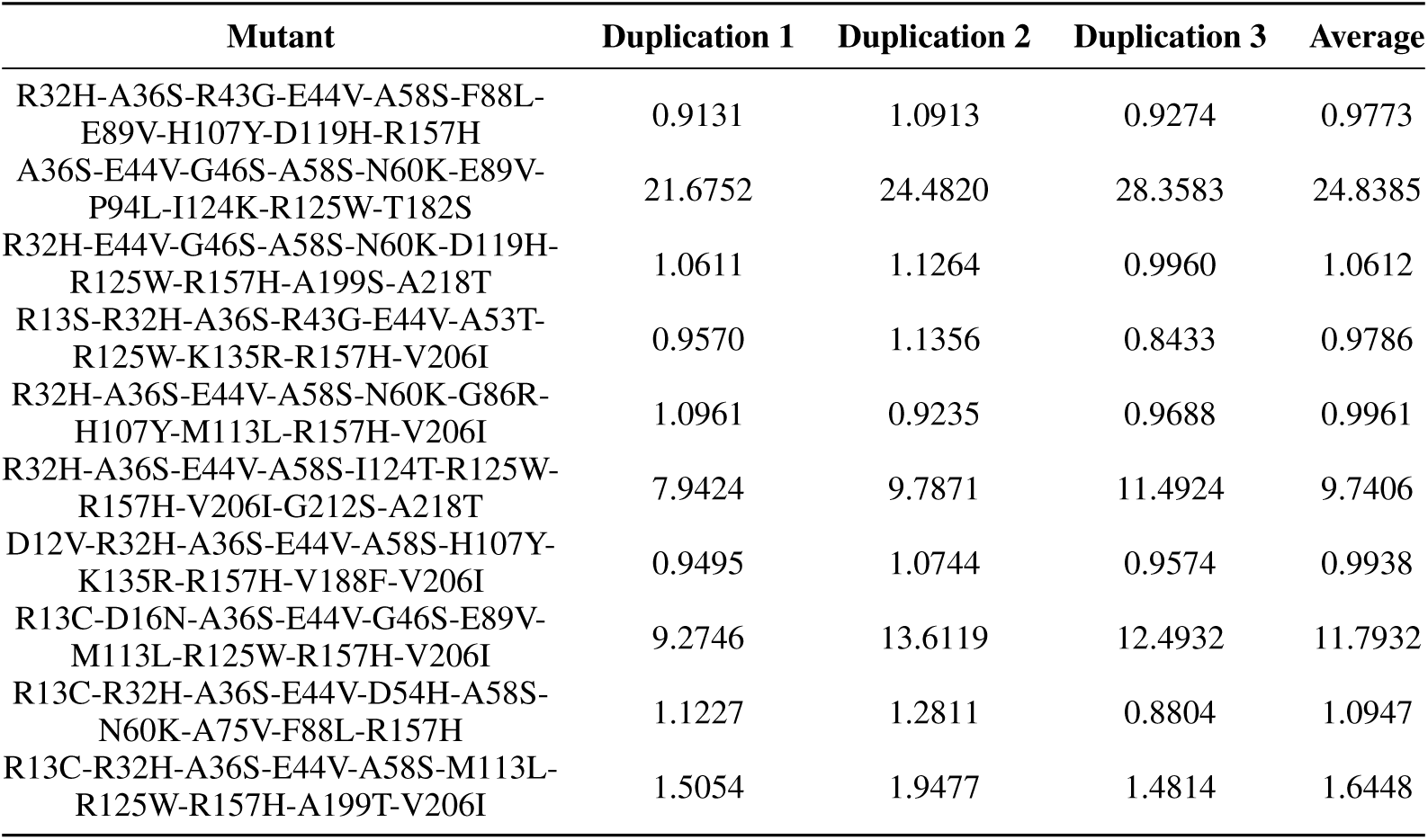
Fold repression of AmeR mutants (Random-Bot)

### 1.4 Results for the evolution process of LCC-ICCG

#### 1.4.1 Hydrolytic reactions catalyzed by LCC

**Supplementary Figure 5.**
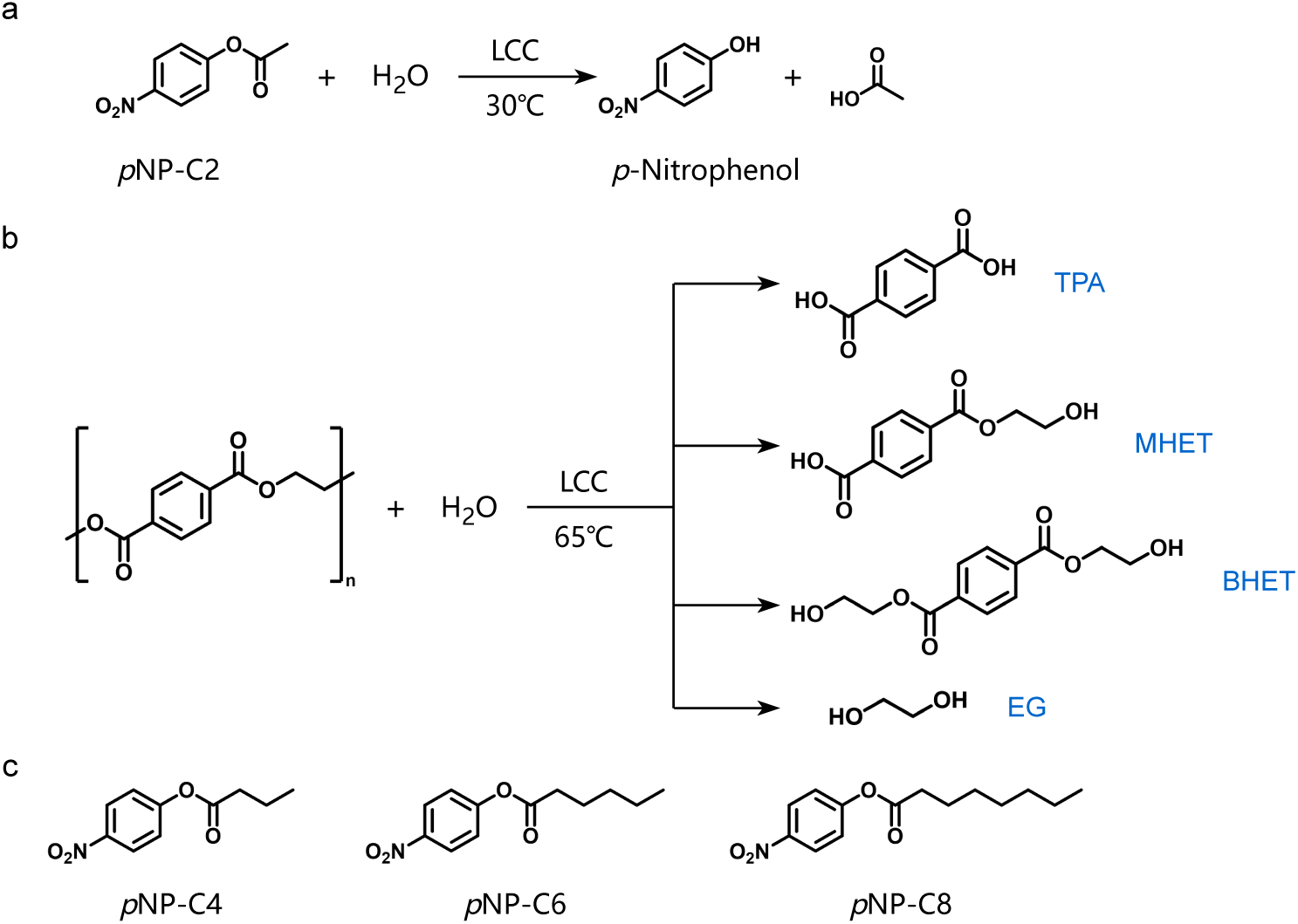
Schematic diagram of reactions catalyzed by LCC. **a)** Hydrolysis of *p*NP-C2, which is also abbreviated as *p*NP in this study. **b)** Hydrolytic products of PET. With improvement in catalytic efficiency, the enzyme tends to achieve thorough depolymerization of PET, creating a higher proportion of TPA rather than MHET and BHET. **c)** *p*NP esters with longer acyl groups than *p*NP-C2 in panel (**a**).

#### 1.4.2 Variation of MLP layers along the optimization rounds

**Supplementary Figure 6.**
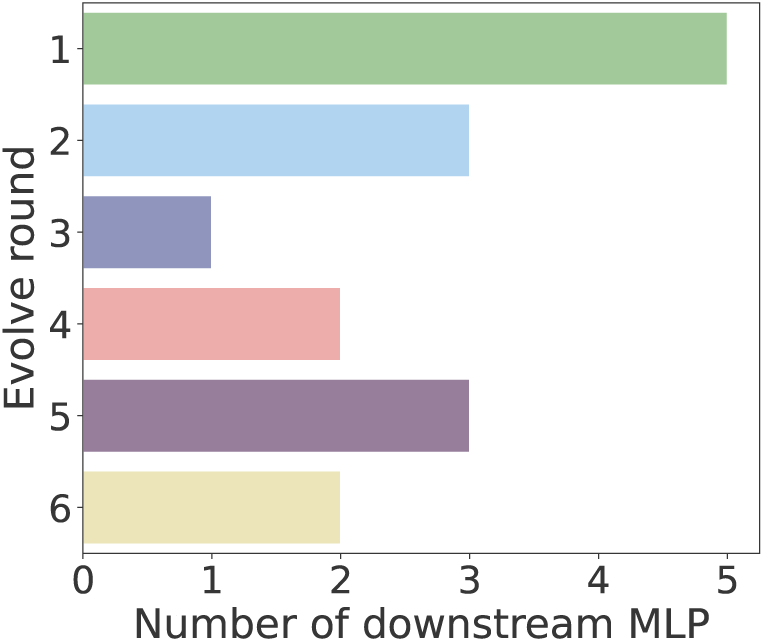
MLP layers of the reward model during the evolution process. The MLP layer configuration within the reward model changes dynamically across the six rounds of RelaVDEP-based sequence evolution.

#### 1.4.3 Experimental verification on *p*NP activity

The enzymatic activity of *p*NP is expressed in units of U/mg, where one unit (U) is defined as the amount of enzyme that catalyzes the conversion of 1 µmol of substrate per minute. The activity was determined by measuring the absorbance of the product *p*-nitrophenol at 415 nm. A standard curve was established by correlating absorbance values with known concentrations of the product, enabling the conversion of absorbance measurements to product concentration. Minor deviations may arise from the need to freshly dilute both the enzyme and the substrate for each assay. Additionally, the substrate undergoes spontaneous hydrolysis in aqueous solution, generating a background reaction rate. All enzymatic activities reported in this study were corrected for this background signal. Mutants exhibiting no detectable activities might yield slightly negative values after background subtraction. Such values were set as zero during the normalization process.

**Supplementary Table 10.** *p*NP activity of LCC-ICCG mutants (Round 1)

| Mutant | Duplication 1 | Duplication 2 | Duplication 3 | Average | Normalized |
| --- | --- | --- | --- | --- | --- |
| WT | 20.1879 | 20.4434 | 22.1268 | 20.9194 | 1.0000 |
| G128E-A172G-T176L-V177F | 0.0215 | -0.1352 | 0.1086 | -0.0017 | 0.0000 |
| Y60H-T153L-E173P-H183S | 0.5033 | 0.3872 | 0.6485 | 0.5130 | 0.0245 |
| T153Y-A178T-P179R-I208G | 0.0000 | 0.0000 | 0.0000 | 0.0000 | 0.0000 |
| G128A-A174G-V177H-A178I | 0.8691 | 0.7413 | 1.0316 | 0.8807 | 0.0421 |
| G128K-S206G-A209F-I217C | 1.7862 | 1.6585 | 1.7804 | 1.7417 | 0.0833 |
| T153A-V170G-A174R-V177Q | 3.5335 | 3.6090 | 3.9921 | 3.7115 | 0.1774 |
| G128L-L152I-A172G-H183Q | 10.5807 | 10.3776 | 10.8304 | 10.5962 | 0.5065 |
| T153K-A172C-V177P-P179H | 1.2406 | 1.1593 | 1.2232 | 1.2077 | 0.0577 |
| Y60I-T176Y-V177G-I217N | 1.3973 | 1.2696 | 1.2696 | 1.3122 | 0.0627 |
| T153W-A172D-A174Y-H183Y | 0.1667 | 0.1028 | 0.1609 | 0.1434 | 0.0069 |
| P154K-V170A-A174T-P179I | 0.6873 | 0.4830 | 0.5480 | 0.5728 | 0.0274 |
| T153L-P154F-S206P-A209M | 2.7028 | 2.5774 | 2.7446 | 2.6749 | 0.1279 |
| V170L-E173V-V180Q-I208T | 7.9272 | 7.3607 | 7.5000 | 7.5960 | 0.3631 |
| G128W-A174W-V177T-A178M | 0.0093 | -0.1625 | -0.0836 | -0.0789 | 0.0000 |
| G128F-V170H-E173G-I208P | 0.0650 | -0.1068 | -0.0093 | -0.0170 | 0.0000 |
| Y60F-E173S-A174H-H183V | 10.7294 | 10.3811 | 11.0108 | 10.7071 | 0.5118 |
| E173L-H183F-S206H-I217S | 4.2957 | 3.7848 | 3.9520 | 4.0108 | 0.1917 |
| H129V-A174P-S206L-P210I | 0.0604 | -0.3251 | 0.0046 | -0.0867 | 0.0000 |
| Y60S-T176W-V180I-S206M | 2.6981 | 1.8808 | 2.4009 | 2.3266 | 0.1112 |

**Supplementary Table 11.**
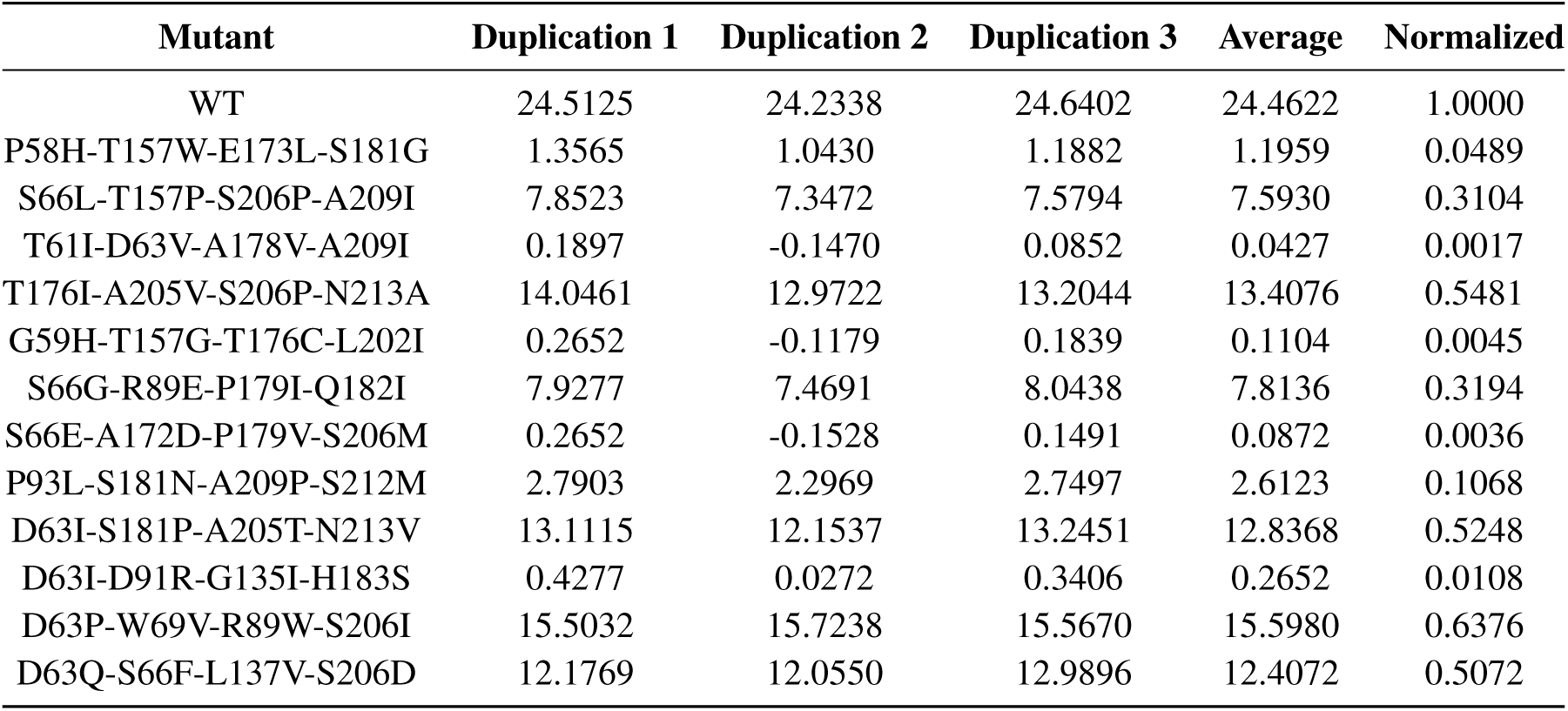
*p*NP activity of LCC-ICCG mutants (Round 2)

**Supplementary Table 12.** *p*NP activity of LCC-ICCG mutants (Round 3)

| Mutant | Duplication 1 | Duplication 2 | Duplication 3 | Average | Normalized |
| --- | --- | --- | --- | --- | --- |
| WT | 20.7237 | 20.7237 | 21.4203 | 20.9559 | 1.0000 |
| D63L-P154F | 1.6834 | 1.7415 | 1.9737 | 1.7995 | 0.0859 |
| D91P-S206M | 2.9025 | 2.8444 | 3.1927 | 2.9799 | 0.1422 |
| A62P-D91W-E173R | -0.1741 | -0.2902 | 0.0000 | -0.1548 | 0.0000 |
| T61F-V170G-A209F | 9.2299 | 9.2879 | 10.3328 | 9.6169 | 0.4589 |
| D63L-E173L-N213A | 14.3963 | 14.7446 | 15.1509 | 14.7639 | 0.7045 |
| D91M-K159T-S206F | 20.5495 | 21.0720 | 23.0457 | 21.5557 | 1.0286 |
| W69F-G132I-H183W | 0.2322 | -0.1161 | 0.0000 | 0.0387 | 0.0018 |
| S57E-V180Q-A209P | 0.2322 | -0.1741 | 0.0580 | 0.0387 | 0.0018 |
| D91P-H156L-K159H-V180D | 0.2322 | -0.0580 | 0.1161 | 0.0967 | 0.0046 |
| A62D-R89L-R96V-N204K | 0.4063 | 0.0580 | 0.2322 | 0.2322 | 0.0111 |
| M131S-H156L-S181T-N213P | 4.0054 | 4.7020 | 4.9342 | 4.5472 | 0.2170 |
| P58F-D91M-H156D-V180Q | -0.0580 | -0.1741 | 0.0580 | -0.0580 | 0.0000 |
| I169M-S181Q-I185W-N213P | 19.7949 | 20.7237 | 21.4783 | 20.6656 | 0.9861 |
| L67F-D91P-A172T-S206M | -0.2902 | -0.2902 | 0.0000 | -0.1935 | 0.0000 |
| A62L-D63T-V170F-H183R | 0.6385 | 0.5805 | 0.8127 | 0.6772 | 0.0323 |
| A62P-G134R-A174I-H183P | -0.1161 | -0.1741 | 0.0580 | -0.0774 | 0.0000 |

**Supplementary Table 13.**
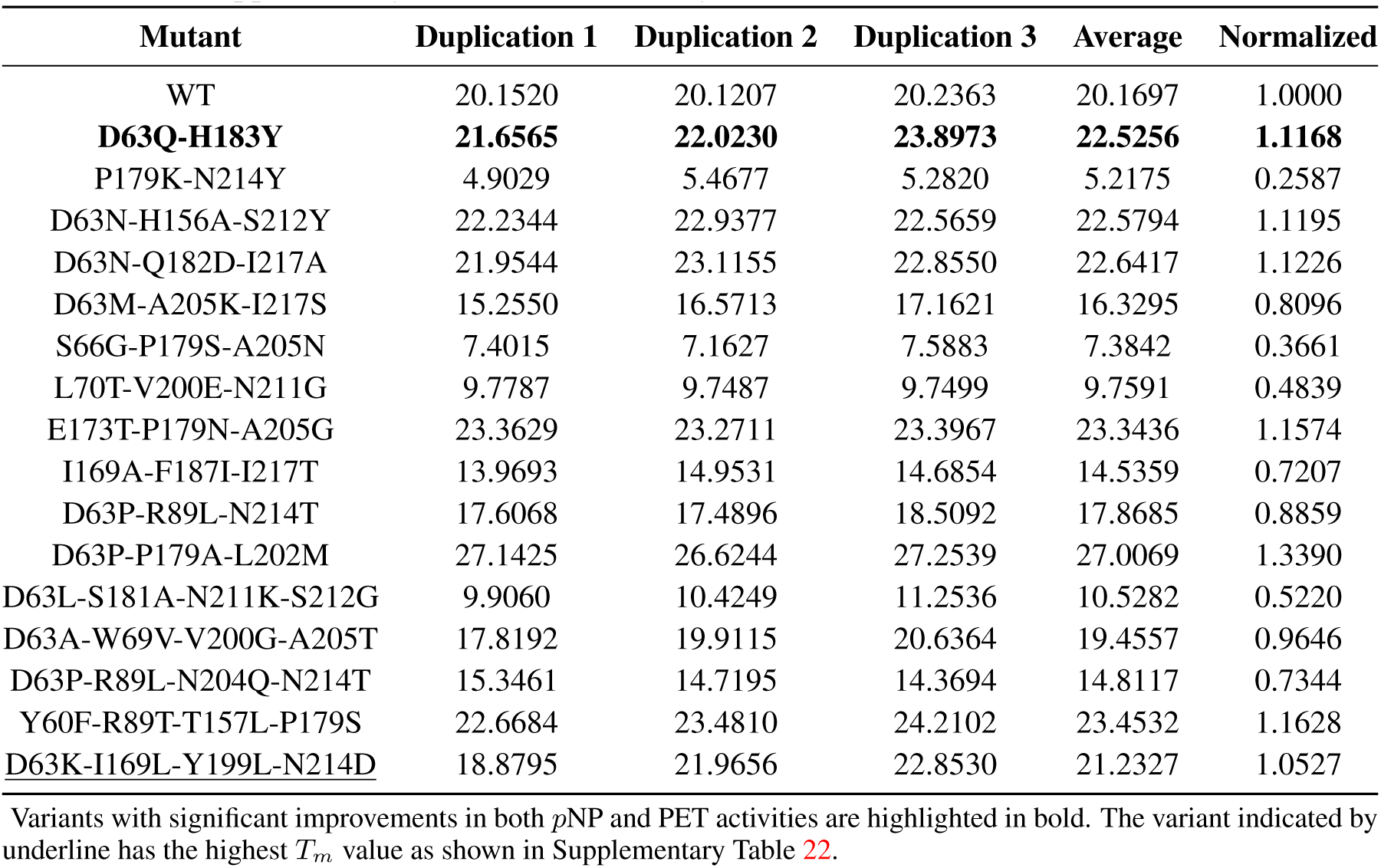
*p*NP activity of LCC-ICCG mutants (Round 4)

| Mutant | Duplication 1 | Duplication 2 | Duplication 3 | Average | Normalized |
| --- | --- | --- | --- | --- | --- |
| WT | 20.1520 | 20.1207 | 20.2363 | 20.1697 | 1.0000 |
| **D63Q-H183Y** | **21.6565** | **22.0230** | **23.8973** | **22.5256** | **1.1168** |
| P179K-N214Y | 4.9029 | 5.4677 | 5.2820 | 5.2175 | 0.2587 |
| D63N-H156A-S212Y | 22.2344 | 22.9377 | 22.5659 | 22.5794 | 1.1195 |
| D63N-Q182D-I217A | 21.9544 | 23.1155 | 22.8550 | 22.6417 | 1.1226 |
| D63M-A205K-I217S | 15.2550 | 16.5713 | 17.1621 | 16.3295 | 0.8096 |
| S66G-P179S-A205N | 7.4015 | 7.1627 | 7.5883 | 7.3842 | 0.3661 |
| L70T-V200E-N211G | 9.7787 | 9.7487 | 9.7499 | 9.7591 | 0.4839 |
| E173T-P179N-A205G | 23.3629 | 23.2711 | 23.3967 | 23.3436 | 1.1574 |
| I169A-F187I-I217T | 13.9693 | 14.9531 | 14.6854 | 14.5359 | 0.7207 |
| D63P-R89L-N214T | 17.6068 | 17.4896 | 18.5092 | 17.8685 | 0.8859 |
| D63P-P179A-L202M | 27.1425 | 26.6244 | 27.2539 | 27.0069 | 1.3390 |
| D63L-S181A-N211K-S212G | 9.9060 | 10.4249 | 11.2536 | 10.5282 | 0.5220 |
| D63A-W69V-V200G-A205T | 17.8192 | 19.9115 | 20.6364 | 19.4557 | 0.9646 |
| D63P-R89L-N204Q-N214T | 15.3461 | 14.7195 | 14.3694 | 14.8117 | 0.7344 |
| Y60F-R89T-T157L-P179S | 22.6684 | 23.4810 | 24.2102 | 23.4532 | 1.1628 |
| <u>D63K-I169L-Y199L-N214D</u> | 18.8795 | 21.9656 | 22.8530 | 21.2327 | 1.0527 |

**Supplementary Table 14.** ***p***NP activity of LCC-ICCG mutants (Round 5)

| Mutant | Duplication 1 | Duplication 2 | Duplication 3 | Average | Normalized |
| --- | --- | --- | --- | --- | --- |
| WT | 21.0797 | 20.2176 | 23.2482 | 21.5152 | 1.0000 |
| **A174K-N204R** | **27.6665** | **26.2660** | **30.0062** | **27.9796** | **1.3005** |
| N204F-S212M | 25.4376 | 22.6569 | 26.2865 | 24.7937 | 1.1524 |
| Y199R-N213P | 25.1556 | 21.1639 | 23.4815 | 23.2670 | 1.0814 |
| S181Q-S212P | 17.3473 | 15.1907 | 16.1133 | 16.2171 | 0.7537 |
| R96D-G135H-L137S | 0.0000 | 0.0000 | 0.0000 | 0.0000 | 0.0000 |
| D63N-A205E-I217P | 20.6731 | 19.1444 | 19.5457 | 19.7877 | 0.9197 |
| T176Q-V180L-N213D | 31.0914 | 29.6541 | 33.6640 | 31.4698 | 1.4627 |
| A174W-Y199I-N213E | 21.7301 | 20.8695 | 24.3648 | 22.3215 | 1.0375 |
| L70M-G171H-T176D | 0.0000 | 0.0000 | 0.0000 | 0.0000 | 0.0000 |
| S66D-K159Q-Y199Q-N213P | 7.4832 | 9.9369 | 9.4282 | 8.9494 | 0.4160 |
| S181F-I185Q-A205S-N213H | 24.1717 | 26.8109 | 27.7018 | 26.2281 | 1.2191 |
| R89L-L137D-P179F-I217A | 0.0000 | 0.0000 | 0.0000 | 0.0000 | 0.0000 |
| D63L-N204A-A205M-I217S | 14.6502 | 14.4440 | 17.8322 | 15.6421 | 0.7270 |

Variants with significant improvements in both *p*NP and PET activities are highlighted in bold.

**Supplementary Table 15.** ***p***NP activity of LCC-ICCG mutants (Round 6)

| Mutant | Duplication 1 | Duplication 2 | Duplication 3 | Average | Normalized |
| --- | --- | --- | --- | --- | --- |
| WT | 19.9361 | 21.6263 | 22.1008 | 21.2211 | 1.0000 |
| I169W-N213E | 19.3570 | 19.6448 | 18.7968 | 19.2662 | 0.9079 |
| I169H-E201H | 11.1509 | 11.3078 | 11.7673 | 11.4087 | 0.5376 |
| T176N-Q182A | 21.9507 | 21.0359 | 21.5875 | 21.5247 | 1.0143 |
| **T157P-T176N** | **30.2772** | **28.5782** | **30.0847** | **29.6467** | **1.3970** |
| T157Q-N213D | 30.8413 | 29.9032 | 31.4897 | 30.7447 | 1.4488 |
| V150Y-T176H-Y199R | 1.5546 | 1.7470 | 1.7340 | 1.6785 | 0.0791 |
| T176D-Y199L-N211Q | 21.2859 | 21.6618 | 23.1328 | 22.0268 | 1.0380 |
| N211D-S212E-N214Q | 6.7540 | 6.9558 | 6.4651 | 6.7249 | 0.3169 |
| V150F-K159L-I169F | 25.3869 | 24.5838 | 27.2257 | 25.7321 | 1.2126 |
| T157A-S212K-N213M | 25.5657 | 24.8683 | 26.7695 | 25.7345 | 1.2127 |
| L70F-T176S-A178D-N211A | 0.8481 | 0.7038 | 0.9117 | 0.8212 | 0.0387 |
| G135M-V150Q-A205E-S212R | 1.7011 | 1.9224 | 1.9925 | 1.8720 | 0.0882 |
| A174Q-H183I-S212R-N213R | 10.7224 | 11.9294 | 12.8178 | 11.8232 | 0.5571 |
| I169M-S181G-I185M-N213F | 21.2046 | 21.9079 | 22.9275 | 22.0133 | 1.0373 |
| I169Y-T176P-S206D-A209F | 8.3113 | 9.3924 | 10.0296 | 9.2444 | 0.4356 |

Variants with significant improvements in both *p*NP and PET activities are highlighted in bold.

#### 1.4.4 Experimental verification on PET activity

**Supplementary Table 16.** PET activity of LCC-ICCG mutants (Round 1)

| Mutant | Duplication 1 | Duplication 2 | Duplication 3 | Average | Normalized |
| --- | --- | --- | --- | --- | --- |
| WT | 24.6191 | 26.3319 | 24.7372 | 25.2294 | 1.0000 |
| G128E-A172G-T176L-V177F | 0.0330 | 0.0000 | 0.0000 | 0.0110 | 0.0004 |
| Y60H-T153L-E173P-H183S | 0.0543 | 0.0702 | 0.0766 | 0.0670 | 0.0027 |
| T153Y-A178T-P179R-I208G | 0.0000 | 0.0000 | 0.0000 | 0.0000 | 0.0000 |
| G128A-A174G-V177H-A178I | 0.0191 | 0.0085 | 0.0149 | 0.0142 | 0.0006 |
| G128K-S206G-A209F-I217C | 0.0457 | 0.0468 | 0.0489 | 0.0472 | 0.0019 |
| T153A-V170G-A174R-V177Q | 9.7191 | 9.1691 | 10.4638 | 9.7840 | 0.3878 |
| G128L-L152I-A172G-H183Q | 15.1894 | 13.5553 | 12.6585 | 13.8011 | 0.5470 |
| T153K-A172C-V177P-P179H | 0.4085 | 0.8479 | 0.6872 | 0.6479 | 0.0257 |
| Y60I-T176Y-V177G-I217N | 0.1340 | 0.1660 | 0.1266 | 0.1422 | 0.0056 |
| T153W-A172D-A174Y-H183Y | 0.0000 | 0.0000 | 0.0000 | 0.0000 | 0.0000 |
| P154K-V170A-A174T-P179I | 0.0298 | 0.0170 | 0.0362 | 0.0277 | 0.0011 |
| T153L-P154F-S206P-A209M | 0.0670 | 0.0723 | 0.0670 | 0.0688 | 0.0027 |
| V170L-E173V-V180Q-I208T | 7.5000 | 8.6723 | 8.4319 | 8.2014 | 0.3251 |
| G128W-A174W-V177T-A178M | 0.0000 | 0.0000 | 0.0000 | 0.0000 | 0.0000 |
| G128F-V170H-E173G-I208P | 0.0000 | 0.0000 | 0.0000 | 0.0000 | 0.0000 |
| Y60F-E173S-A174H-H183V | 17.0436 | 16.2362 | 14.9255 | 16.0684 | 0.6369 |
| E173L-H183F-S206H-I217S | 6.3149 | 6.2723 | 5.4074 | 5.9982 | 0.2377 |
| H129V-A174P-S206L-P210I | 0.0543 | 0.0468 | 0.0479 | 0.0496 | 0.0020 |
| Y60S-T176W-V180I-S206M | 3.4670 | 3.4032 | 3.4160 | 3.4287 | 0.1359 |

**Supplementary Table 17.** PET activity of LCC-ICCG mutants (Round 2)

| Mutant | Duplication 1 | Duplication 2 | Duplication 3 | Average | Normalized |
| --- | --- | --- | --- | --- | --- |
| WT | 24.6152 | 24.1729 | 22.7615 | 23.8499 | 1.0000 |
| P58H-T157W-E173L-S181G | 1.2163 | 1.0727 | 2.2615 | 1.5168 | 0.0636 |
| S66L-T157P-S206P-A209I | 11.9902 | 12.0372 | 11.8582 | 11.9619 | 0.5015 |
| T61I-D63V-A178V-A209I | 0.6463 | 0.6002 | 0.5399 | 0.5954 | 0.0250 |
| T176I-A205V-S206P-N213A | 18.2385 | 17.3351 | 17.0887 | 17.5541 | 0.7360 |
| G59H-T157G-T176C-L202I | 0.0346 | 0.0257 | 0.0284 | 0.0296 | 0.0012 |
| S66G-R89E-P179I-Q182I | 0.0585 | 0.1746 | 0.0381 | 0.0904 | 0.0038 |
| S66E-A172D-P179V-S206M | 0.0000 | 0.0293 | 0.0213 | 0.0168 | 0.0007 |
| P93L-S181N-A209P-S212M | 0.5736 | 0.7642 | 0.5629 | 0.6336 | 0.0266 |
| D63I-S181P-A205T-N213V | 7.6720 | 7.8307 | 8.3457 | 7.9495 | 0.3333 |
| D63I-D91R-G135I-H183S | 0.0000 | 0.0000 | 0.0222 | 0.0074 | 0.0003 |
| D63P-W69V-R89W-S206I | 13.2234 | 13.1445 | 12.7004 | 13.0228 | 0.5460 |
| D63Q-S66F-L137V-S206D | 5.9521 | 5.9628 | 6.6082 | 6.1743 | 0.2589 |

**Supplementary Table 18.** PET activity of LCC-ICCG mutants (Round 3)

| Mutant | Duplication 1 | Duplication 2 | Duplication 3 | Average | Normalized |
| --- | --- | --- | --- | --- | --- |
| WT | 24.7957 | 22.0931 | 23.0980 | 23.3289 | 1.0000 |
| D63L-P154F | 0.0000 | 0.0000 | 0.0000 | 0.0000 | 0.0000 |
| D91P-S206M | 0.5451 | 0.5382 | 0.4287 | 0.5040 | 0.0216 |
| A62P-D91W-E173R | 0.0000 | 0.0000 | 0.0000 | 0.0000 | 0.0000 |
| T61F-V170G-A209F | 13.6880 | 12.9147 | 10.7238 | 12.4422 | 0.5333 |
| D63L-E173L-N213A | 19.0517 | 15.0445 | 14.6688 | 16.2550 | 0.6968 |
| D91M-K159T-S206F | 13.4391 | 13.1809 | 11.5594 | 12.7265 | 0.5455 |
| W69F-G132I-H183W | 0.0000 | 0.0000 | 0.0000 | 0.0000 | 0.0000 |
| S57E-V180Q-A209P | 0.0000 | 0.0000 | 0.0000 | 0.0000 | 0.0000 |
| D91P-H156L-K159H-V180D | 0.0000 | 0.0000 | 0.0000 | 0.0000 | 0.0000 |
| A62D-R89L-R96V-N204K | 0.0000 | 0.0000 | 0.0000 | 0.0000 | 0.0000 |
| M131S-H156L-S181T-N213P | 0.2812 | 0.4057 | 0.6143 | 0.4337 | 0.0186 |
| P58F-D91M-H156D-V180Q | 0.0000 | 0.0000 | 0.0000 | 0.0000 | 0.0000 |
| I169M-S181Q-I185W-N213P | 22.9972 | 20.6809 | 22.7121 | 22.1300 | 0.9486 |
| L67F-D91P-A172T-S206M | 0.0000 | 0.0000 | 0.0000 | 0.0000 | 0.0000 |
| A62L-D63T-V170F-H183R | 2.7464 | 2.6311 | 2.5816 | 2.6530 | 0.1137 |
| A62P-G134R-A174I-H183P | 0.0000 | 0.0000 | 0.0000 | 0.0000 | 0.0000 |

**Supplementary Table 19.** PET activity of LCC-ICCG mutants (Round 4)

| Mutant | Duplication 1 | Duplication 2 | Duplication 3 | Average | Normalized |
| --- | --- | --- | --- | --- | --- |
| WT | 22.7872 | 25.1702 | 25.4695 | 24.4757 | 1.0000 |
| **D63Q-H183Y** | **27.5305** | **29.5716** | **28.0638** | **28.3887** | **1.1599** |
| P179K-N214Y | 2.3816 | 1.8752 | 2.6993 | 2.3187 | 0.0947 |
| D63N-H156A-S212Y | 15.1135 | 15.4369 | 11.2426 | 13.9310 | 0.5692 |
| D63N-Q182D-I217A | 16.0326 | 15.1631 | 13.7504 | 14.9820 | 0.6121 |
| D63M-A205K-I217S | 7.9418 | 8.0397 | 8.2525 | 8.0780 | 0.3300 |
| S66G-P179S-A205N | 8.4298 | 9.4071 | 9.5957 | 9.1442 | 0.3736 |
| L70T-V200E-N211G | 0.1078 | 0.1177 | 0.1603 | 0.1286 | 0.0053 |
| E173T-P179N-A205G | 13.5418 | 17.0000 | 15.7475 | 15.4298 | 0.6304 |
| I169A-F187I-I217T | 18.9702 | 16.1390 | 17.0780 | 17.3957 | 0.7107 |
| D63P-R89L-N214T | 16.3078 | 15.5887 | 14.0000 | 15.2988 | 0.6251 |
| D63P-P179A-L202M | 17.3376 | 15.3901 | 16.9929 | 16.5735 | 0.6771 |
| D63L-S181A-N211K-S212G | 4.6411 | 4.5617 | 4.4809 | 4.5612 | 0.1864 |
| D63A-W69V-V200G-A205T | 10.5759 | 9.5801 | 10.2184 | 10.1248 | 0.4137 |
| D63P-R89L-N204Q-N214T | 14.8738 | 13.8851 | 12.5050 | 13.7546 | 0.5620 |
| Y60F-R89T-T157L-P179S | 16.6851 | 16.6567 | 14.1461 | 15.8293 | 0.6467 |
| <u>D63K-I169L-Y199L-N214D</u> | 14.1929 | 13.8270 | 13.0397 | 13.6865 | 0.5592 |

**Supplementary Table 20.** PET activity of LCC-ICCG mutants (Round 5)

| Mutant | Duplication 1 | Duplication 2 | Duplication 3 | Average | Normalized |
| --- | --- | --- | --- | --- | --- |
| WT | 27.7411 | 21.4872 | 23.2461 | 24.1582 | 1.0000 |
| **A174K-N204R** | **32.1326** | **23.6255** | **22.3376** | **26.0319** | **1.0776** |
| N204F-S212M | 26.4262 | 23.9638 | 24.4092 | 24.9331 | 1.0321 |
| Y199R-N213P | 13.9963 | 11.5587 | 10.0730 | 11.8760 | 0.4916 |
| S181Q-S212P | 6.0406 | 5.1147 | 3.8711 | 5.0088 | 0.2073 |
| R96D-G135H-L137S | 0.0000 | 0.0000 | 0.0000 | 0.0000 | 0.0000 |
| D63N-A205E-I217P | 4.5860 | 4.2782 | 3.9952 | 4.2865 | 0.1774 |
| T176Q-V180L-N213D | 22.5738 | 20.3957 | 18.7475 | 20.5723 | 0.8516 |
| A174W-Y199I-N213E | 26.8291 | 23.6266 | 22.7130 | 24.3896 | 1.0096 |
| L70M-G171H-T176D | 0.0000 | 0.0000 | 0.0000 | 0.0000 | 0.0000 |
| S66D-K159Q-Y199Q-N213P | 0.3252 | 0.3699 | 0.4046 | 0.3665 | 0.0152 |
| S181F-I185Q-A205S-N213H | 19.2972 | 17.9064 | 16.5809 | 17.9281 | 0.7421 |
| R89L-L137D-P179F-I217A | 0.0000 | 0.0000 | 0.0000 | 0.0000 | 0.0000 |
| D63L-N204A-A205M-I217S | 5.0440 | 4.8392 | 4.9645 | 4.9492 | 0.2049 |

Variants with significant improvements in both *p*NP and PET activities are highlighted in bold.

**Supplementary Table 21.** PET activity of LCC-ICCG mutants (Round 6)

| Mutant | Duplication 1 | Duplication 2 | Duplication 3 | Average | Normalized |
| --- | --- | --- | --- | --- | --- |
| WT | 25.3000 | 23.6865 | 22.0092 | 23.6652 | 1.0000 |
| I169W-N213E | 0.0858 | 0.1709 | 0.1358 | 0.1309 | 0.0055 |
| I169H-E201H | 0.0000 | 0.0000 | 0.0000 | 0.0000 | 0.0000 |
| T176N-Q182A | 19.5043 | 22.4950 | 19.3702 | 20.4565 | 0.8644 |
| **T157P-T176N** | **29.0362** | **28.6446** | **29.9123** | **29.1977** | **1.2338** |
| T157Q-N213D | 23.2496 | 16.8433 | 22.7326 | 20.9418 | 0.8849 |
| V150Y-T176H-Y199R | 0.0000 | 0.0000 | 0.0000 | 0.0000 | 0.0000 |
| T176D-Y199L-N211Q | 12.6050 | 15.1035 | 11.5908 | 13.0998 | 0.5535 |
| N211D-S212E-N214Q | 0.0277 | 0.0404 | 0.0232 | 0.0304 | 0.0013 |
| V150F-K159L-I169F | 0.0652 | 0.0141 | 0.0601 | 0.0465 | 0.0020 |
| T157A-S212K-N213M | 10.2738 | 15.4241 | 15.2468 | 13.6482 | 0.5767 |
| L70F-T176S-A178D-N211A | 0.0000 | 0.0000 | 0.0000 | 0.0000 | 0.0000 |
| G135M-V150Q-A205E-S212R | 0.0000 | 0.0000 | 0.0000 | 0.0000 | 0.0000 |
| A174Q-H183I-S212R-N213R | 9.2418 | 8.0837 | 8.4574 | 8.5943 | 0.3632 |
| I169M-S181G-I185M-N213F | 14.5759 | 13.3348 | 13.8645 | 13.9251 | 0.5884 |
| I169Y-T176P-S206D-A209F | 0.0000 | 0.0000 | 0.0000 | 0.0000 | 0.0000 |

Variants with significant improvements in both *p*NP and PET activities are highlighted in bold.

#### 1.4.5 Thermostability of RelaVDEP-sampled LCC-ICCG variants

**Supplementary Table 22.**
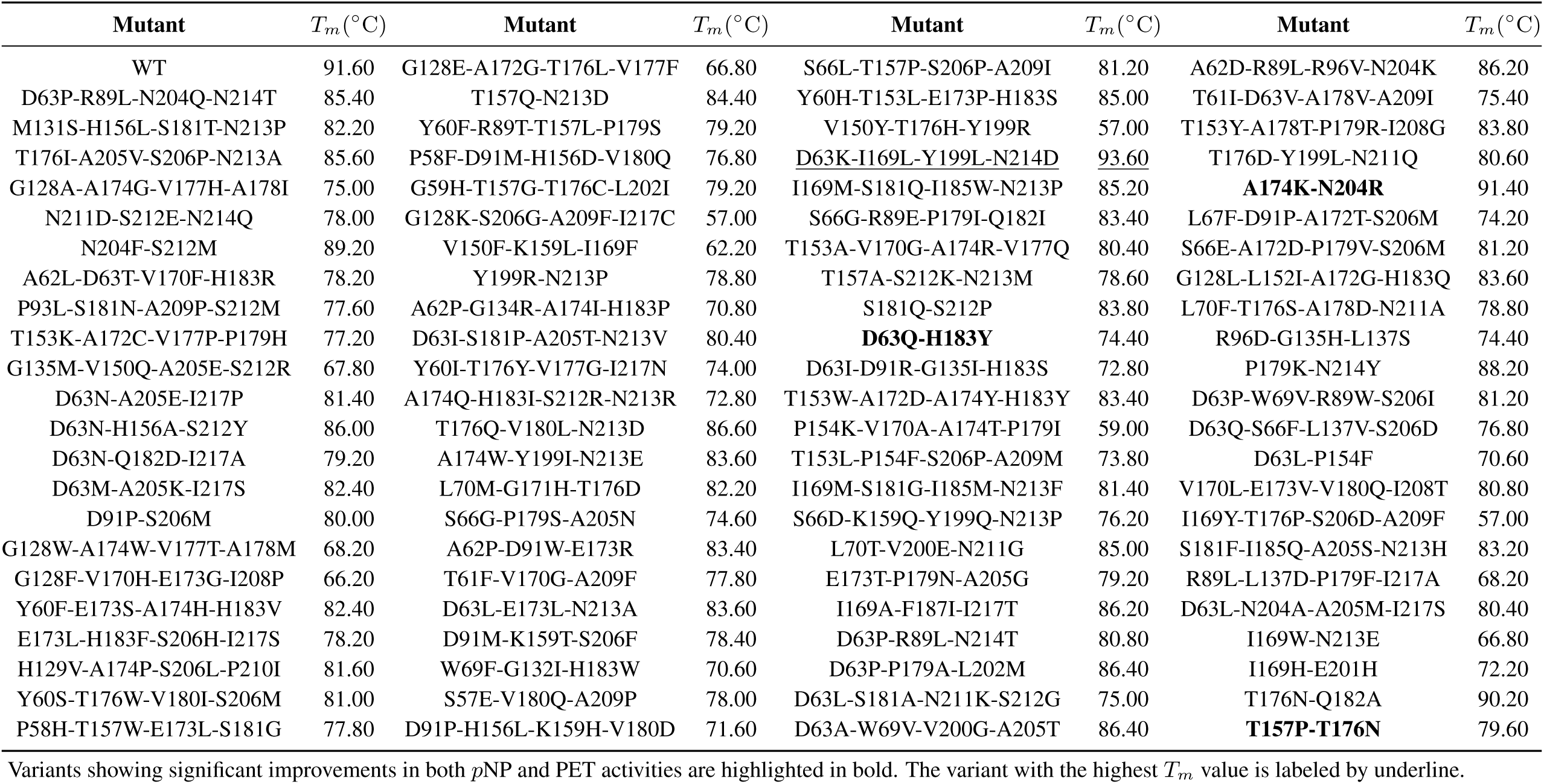
Thermostability of all LCC-ICCG mutants.

| Mutant | $T_m$ (°C) | Mutant | $T_m$ (°C) | Mutant | $T_m$ (°C) | Mutant | $T_m$ (°C) |
| --- | --- | --- | --- | --- | --- | --- | --- |
| WT | 91.60 | G128E-A172G-T176L-V177F | 66.80 | S66L-T157P-S206P-A209I | 81.20 | A62D-R89L-R96V-N204K | 86.20 |
| D63P-R89L-N204Q-N214T | 85.40 | T157Q-N213D | 84.40 | Y60H-T153L-E173P-H183S | 85.00 | T61I-D63V-A178V-A209I | 75.40 |
| M131S-H156L-S181T-N213P | 82.20 | Y60F-R89T-T157L-P179S | 79.20 | V150Y-T176H-Y199R | 57.00 | T153Y-A178T-P179R-I208G | 83.80 |
| T176I-A205V-S206P-N213A | 85.60 | P58F-D91M-H156D-V180Q | 76.80 | D63K-I169L-Y199L-N214D | 93.60 | T176D-Y199L-N211Q | 80.60 |
| G128A-A174G-V177H-A178I | 75.00 | G59H-T157G-T176C-L202I | 79.20 | I169M-S181Q-I185W-N213P | 85.20 | **A174K-N204R** | 91.40 |
| N211D-S212E-N214Q | 78.00 | G128K-S206G-A209F-I217C | 57.00 | S66G-R89E-P179I-Q182I | 83.40 | L67F-D91P-A172T-S206M | 74.20 |
| N204F-S212M | 89.20 | V150F-K159L-I169F | 62.20 | T153A-V170G-A174R-V177Q | 80.40 | S66E-A172D-P179V-S206M | 81.20 |
| A62L-D63T-V170F-H183R | 78.20 | Y199R-N213P | 78.80 | T157A-S212K-N213M | 78.60 | G128L-L152I-A172G-H183Q | 83.60 |
| P93L-S181N-A209P-S212M | 77.60 | A62P-G134R-A174I-H183P | 70.80 | S181Q-S212P | 83.80 | L70F-T176S-A178D-N211A | 78.80 |
| T153K-A172C-V177P-P179H | 77.20 | D63I-S181P-A205T-N213V | 80.40 | **D63Q-H183Y** | 74.40 | R96D-G135H-L137S | 74.40 |
| G135M-V150Q-A205E-S212R | 67.80 | Y60I-T176Y-V177G-I217N | 74.00 | D63I-D91R-G135I-H183S | 72.80 | P179K-N214Y | 88.20 |
| D63N-A205E-I217P | 81.40 | A174Q-H183I-S212R-N213R | 72.80 | T153W-A172D-A174Y-H183Y | 83.40 | D63P-W69V-R89W-S206I | 81.20 |
| D63N-H156A-S212Y | 86.00 | T176Q-V180L-N213D | 86.60 | P154K-V170A-A174T-P179I | 59.00 | D63Q-S66F-L137V-S206D | 76.80 |
| D63N-Q182D-I217A | 79.20 | A174W-Y199I-N213E | 83.60 | T153L-P154F-S206P-A209M | 73.80 | D63L-P154F | 70.60 |
| D63M-A205K-I217S | 82.40 | L70M-G171H-T176D | 82.20 | I169M-S181G-I185M-N213F | 81.40 | V170L-E173V-V180Q-I208T | 80.80 |
| D91P-S206M | 80.00 | S66G-P179S-A205N | 74.60 | S66D-K159Q-Y199Q-N213P | 76.20 | I169Y-T176P-S206D-A209F | 57.00 |
| G128W-A174W-V177T-A178M | 68.20 | A62P-D91W-E173R | 83.40 | L70T-V200E-N211G | 85.00 | S181F-I185Q-A205S-N213H | 83.20 |
| G128F-V170H-E173G-I208P | 66.20 | T61F-V170G-A209F | 77.80 | E173T-P179N-A205G | 79.20 | R89L-L137D-P179F-I217A | 68.20 |
| Y60F-E173S-A174H-H183V | 82.40 | D63L-E173L-N213A | 83.60 | I169A-F187I-I217T | 86.20 | D63L-N204A-A205M-I217S | 80.40 |
| E173L-H183F-S206H-I217S | 78.20 | D91M-K159T-S206F | 78.40 | D63P-R89L-N214T | 80.80 | I169W-N213E | 66.80 |
| H129V-A174P-S206L-P210I | 81.60 | W69F-G132I-H183W | 70.60 | D63P-P179A-L202M | 86.40 | I169H-E201H | 72.20 |
| Y60S-T176W-V180I-S206M | 81.00 | S57E-V180Q-A209P | 78.00 | D63L-S181A-N211K-S212G | 75.00 | T176N-Q182A | 90.20 |
| P58H-T157W-E173L-S181G | 77.80 | D91P-H156L-K159H-V180D | 71.60 | D63A-W69V-V200G-A205T | 86.40 | **T157P-T176N** | 79.60 |

Variants showing significant improvements in both pNP and PET activities are highlighted in bold. The variant with the highest $T_m$ value is labeled by underline.

#### 1.4.6 Catalytic analysis on *p*NP esters of varying lengths

**Supplementary Table 23.** *p*NP-C4 activity of LCC-ICCG mutants.

| Mutant | Duplication 1 | Duplication 2 | Duplication 3 | Average | Normalized |
| --- | --- | --- | --- | --- | --- |
| WT | 27.1725 | 33.4157 | 38.8950 | 33.1610 | 1.0000 |
| A174K-N204R | 37.1330 | 37.2545 | 59.4053 | 44.5976 | 1.3449 |
| T176Q-V180L-N213D | 29.7699 | 37.5539 | 45.4376 | 37.5871 | 1.1335 |
| S181F-I185Q-A205S-N213H | 37.3695 | 42.5360 | 50.6416 | 43.5157 | 1.3123 |
| T157P-T176N | 45.2114 | 47.6899 | 51.3300 | 48.0771 | 1.4498 |
| T157Q-N213D | 36.7622 | 45.4775 | 43.8018 | 42.0138 | 1.2670 |
| V150F-K159L-I169F | 24.3350 | 19.4254 | 27.8133 | 23.8579 | 0.7195 |
| T157A-S212K-N213M | 46.7509 | 54.2058 | 50.4442 | 50.4670 | 1.5219 |

**Supplementary Table 24.** *p*NP-C6 activity of LCC-ICCG mutants.

| Mutant | Duplication 1 | Duplication 2 | Duplication 3 | Average | Normalized |
| --- | --- | --- | --- | --- | --- |
| WT | 29.7154 | 31.5490 | 31.2833 | 30.8492 | 1.0000 |
| A174K-N204R | 33.9286 | 41.4427 | 53.8336 | 43.0683 | 1.3961 |
| T176Q-V180L-N213D | 28.8602 | 35.0961 | 53.5168 | 39.1577 | 1.2693 |
| S181F-I185Q-A205S-N213H | 34.4263 | 38.5399 | 56.4367 | 43.1343 | 1.3982 |
| T157P-T176N | 48.2711 | 61.2496 | 63.6434 | 57.7214 | 1.8711 |
| T157Q-N213D | 55.9404 | 82.5533 | 84.9000 | 74.4646 | 2.4138 |
| V150F-K159L-I169F | 35.6048 | 42.0329 | 40.5699 | 39.4025 | 1.2773 |
| T157A-S212K-N213M | 64.6045 | 78.1490 | 92.0606 | 78.2714 | 2.5372 |

**Supplementary Table 25.** *p*NP-C8 activity of LCC-ICCG mutants.

| Mutant | Duplication 1 | Duplication 2 | Duplication 3 | Average | Normalized |
| --- | --- | --- | --- | --- | --- |
| WT | 33.9267 | 35.9483 | 32.0128 | 33.9626 | 1.0000 |
| A174K-N204R | 29.9834 | 33.5917 | 29.7143 | 31.0964 | 0.9156 |
| T176Q-V180L-N213D | 33.2745 | 37.1975 | 31.5873 | 34.0197 | 1.0017 |
| S181F-I185Q-A205S-N213H | 40.8529 | 44.9323 | 38.0301 | 41.2718 | 1.2152 |
| T157P-T176N | 42.1998 | 48.0208 | 38.3251 | 42.8486 | 1.2616 |
| T157Q-N213D | 42.8746 | 50.1719 | 44.2392 | 45.7619 | 1.3474 |
| V150F-K159L-I169F | 20.5556 | 21.7691 | 24.4630 | 22.2626 | 0.6555 |
| T157A-S212K-N213M | 54.3258 | 55.8957 | 61.1963 | 57.1392 | 1.6824 |

### 1.5 Summarization on the time consumption of the virtual DE

**Supplementary Table 26.** The time consumption (hours) of different stages.

| DE task | Stage 1 | Stage 2 | Stage 3 | Stage 4 | Total |
| --- | --- | --- | --- | --- | --- |
| GFP | - | 28.56 | 12.87 | 0.12 | 41.55 |
| NUDT15 | 11.3 | 1.52 | 11.27 | 0.07 | 24.16 |
| VKOR1 | 9.8 | 1.40 | 10.82 | 0.05 | 22.07 |
| AmeR | 0.85 | 0.14 | 14.57 | 0.09 | 15.65 |
| LCC-ICCG round 1 | 0.32 | 0.07 | 17.2 | 0.15 | 17.74 |
| LCC-ICCG round 2 | 0.28 | 0.12 | 12.95 | 0.18 | 13.53 |
| LCC-ICCG round 3 | 0.13 | 0.03 | 14.65 | 0.22 | 15.03 |
| LCC-ICCG round 4 | 0.17 | 0.03 | 13.00 | 0.24 | 13.44 |
| LCC-ICCG round 5 | 0.24 | 0.05 | 12.03 | 0.30 | 12.62 |
| LCC-ICCG round 6 | 0.30 | 0.05 | 14.78 | 0.27 | 15.40 |

Stage 1: Cross-validation for hyperparameter optimization of the reward model.
Stage 2: Supervised fine-tuning of the reward model.
Stage 3: Training and sampling of the RL framework.
Stage 4: Construction of the mutant library.

## 2 Supplementary Methods

### 2.1 Fine-tuning process of the reward model

**Supplementary Table 27.** Fine-tuning strategy for small datasets.

|  | Stage 1 | Stage 2 | Stage 3 |
| --- | --- | --- | --- |
| **Data splitting** | 7:1:2 | 8:2 | 8:2 |
| **Initial learning rate** | 1e-4 | 1e-4 | 1e1 |
| **Loss function** | Soft Spearman loss | Soft Spearman loss | MSE loss |
| **Training parameters** | $N$ | MLPs | $\alpha/\beta$ |

$N$ is a hyperparameter denoting the number of MLP layers. $\alpha$ and $\beta$ are two learnable parameters in the last linear layer, which connects the difference of MLP predictions on wild-type and mutant sequences and the final output of the fitness change.

**Supplementary Table 28.** Fine-tuning strategy for large datasets.

|  | Stage 1 | Stage 2 |
| --- | --- | --- |
| **Data splitting** | 8:2 | 8:2 |
| **Initial learning rate** | 5e-3 | 1e2 |
| **Loss function** | Soft Spearman loss | MSE loss |
| **Training parameters** | GAT module + MLPs | $\alpha/\beta$ |

$\alpha$ and $\beta$ are two learnable parameters in the last linear layer, which connects the difference of MLP predictions on wild-type and mutant sequences and the final output of the fitness change.

The strategy for fine-tuning the reward model depends on the size of the dataset, as shown in Supplementary Tables 27 and 28. In both strategies, the soft Spearman loss^4^ is first employed to optimize parameters or hyperparameters in the GAT module and MLPs by constraining the agreement in ranking between predictions and labels, while the MSE loss is eventually adopted to tune the two learnable parameters *α* and *β* in the last linear layer in order to adjust the magnitude of output to match that of the original data label without breaking the relative ranking that has already been optimized.

### 2.2 Distributed architecture of RelaVDEP

**Supplementary Figure 7.**
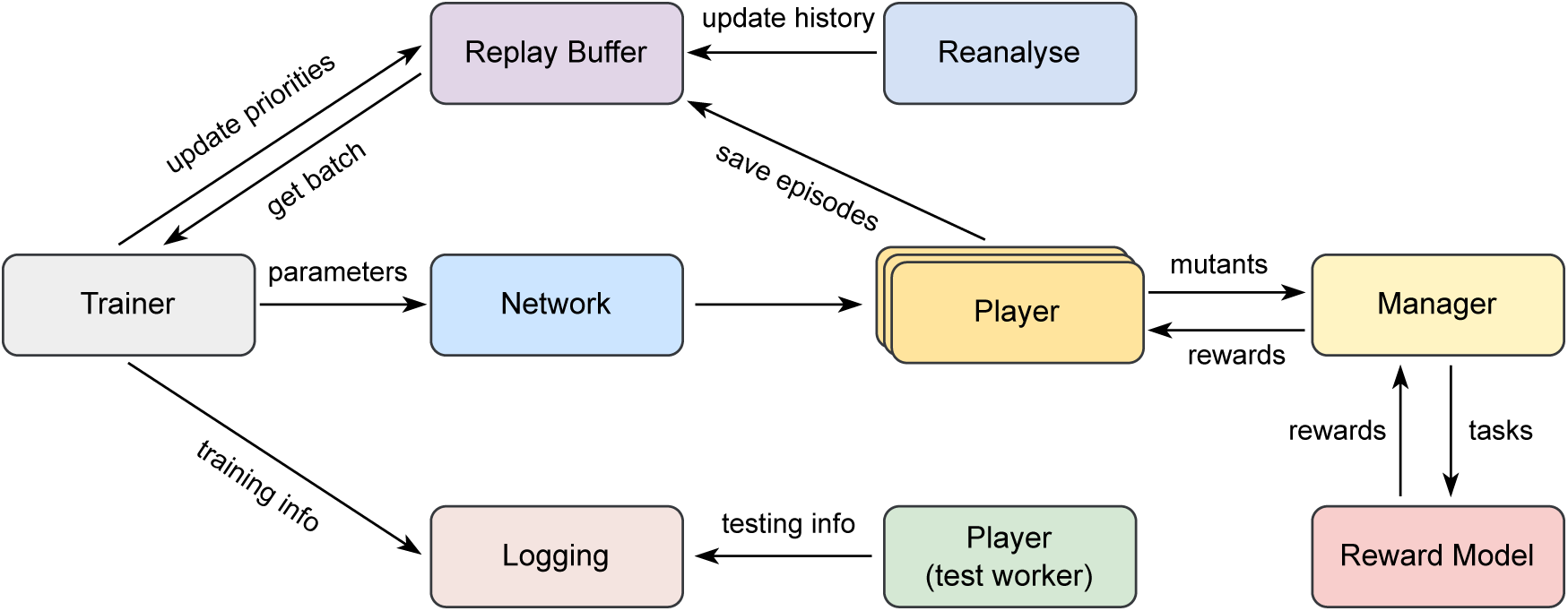
RelaVDEP architecture in the distributed computation framework.

Leveraging the Ray^5^ distributed computing framework, we parallelized RelaVDEP’s workflow by deploying its modular components across dedicated threads for concurrent training and sampling operations. The “Trainer” module continuously updates network parameters, while the “Player” modules utilize the latest parameters to sample mutant variants, storing resultant episodes in a shared “Replay Buffer” for subsequent training. A dedicated “Test” worker evaluates model performance under exploitation-dominated search regimes (*i.e.* high utilization and low exploration). To address the requirements of the “Reward Model” during inference, we decoupled it from individual environments and implemented a shared singleton instance for mutant fitness prediction. The “Manager” module orchestrates this workflow by: (1) collecting mutant sequences from multiple “Players” at varying time steps; (2) serializing task inputs to the centralized “Reward Model”; and (3) propagating predicted rewards back to their respective environments. Simultaneously, a “Logging” module monitors and records real-time operational metrics across all components.

### 2.3 Starting sequences for optimization targets in this study

GFP (length=238):

MSKGEELFTGVVPILVELDGDVNGHKFSVSGEGEGDATYGKLTLKFICTTGKLPVPWPTLVTTLSYGVQCFSRYPDHMKQHDFFKSAMPEGYVQERTIFFKDDGNYKTRAEVKFEGDTLVNRIELKGIDFKEDGNILGHKLEYNYNSHNVYIMADKQKNGIKVNFKIRHNIEDGSVQLADHYQQNTPIGDGPVLLPDNHYLSTQSALSKDPNEKRDHMVLLEFVTAAGITHGMDELYK

NUDT15 (length=164):

MTASAQPRGRRPGVGVGVVVTSCKHPRCVLLGKRKGSVGAGSFQLPGGHLEFGETWEECAQRETWEEAALHLKNVHFASVVNSFIEKENYHYVTILMKGEVDVTHDSEPKNVEPEKNESWEWVPWEELPPLDQLFWGLRCLKEQGYDPFKEDLNHLVGYKGNHL

VKOR1 (length=163):

MGSTWGSPGWVRLALCLTGLVLSLYALHVKAARARDRDYRALCDVGTAISCSRVFSSRWGRGFGLVEHVLGQDSILNQSNSIFGCIFYTLQLLLGCLRTRWASVLMLLSSLVSLAGSVYLAWILFFVLYDFCIVCITTYAINVSLMWLSFRKVQEPQGKAKRH

AmeR (length=219):

MNKTIDQVRKGDRKSDLPVRRRPRRSAEETRRDILAKAEELFRERGFNAVAIADIASALNMSPANVFKHFSSKNALVDAIGFGQIGVFERQICPLDKSHAPLDRLRHLARNLMEQHHQDHFKHIRVFIQILMTAKQDMKCGDYYKSVIAKLLAEIIRDGVEAGLYIATDIPVLAETVLHALTSVIHPVLIAQEDIGNLATRCDQLVDLIDAGLRNPL AK

LCC-ICCG (length=258):

SNPYQRGPNPTRSALTADGPFSVATYTVSRLSVSGFGGGVIYYPTGTSLTFGGIAMSPGYTADASSLAWLGRRLASHGFVVLVINTNSRFDGPDSRASQLSAALNYLRTSSPSAVRARLDANRLAVAGHSMGGGGTLRIAEQNPSLKAAVPLTPWHTDKTFNTSVPVLIVGAEADTVAPVSQHAIPFYQNLPSTTPKVYVELCNASHIAPNSNNAAISVYTISWMKLWVDNDTRYRQFLCNVNDPALCDFRTNNRHCQ

### 2.4 Experimental details for the evolution task of GFP

#### 2.4.1 Construction of mutants

All mutant fragments were assembled into the kanamycin-resistant PET-28a plasmid vector using the NEBuilder HiFi DNA Assembly (NEB) method. The constructed plasmids were transformed into BL21(DE3) competent cells (from Tsingke Biotech Co., Ltd.) for subsequent experiments. The synthetic mutant fragments were ordered from Dynegene Technologies. The plasmid was extracted using the Tiangen Biotech extraction kit, and Sanger sequencing was completed by RuiBiotech.

#### 2.4.2 Fluorescence intensity verification

The identified BL21(DE3) strain was used in flow cytometry experiments. Individual colonies were inoculated into 4 mL of Luria-Bertani (LB) liquid medium with 50 µg/ml kanamycin and cultured overnight. Then, the saturated culture was inoculated into 20 mL of LB liquid medium with 50 µg/ml kanamycin at a 1:100 ratio and cultured for 6 hours at 37 ^◦^C and 200 rpm. When the optical density at 600 nm reached 0.5, the culture was then inoculated into deep well plates containing 50 µg/ml kanamycin LB liquid medium at a 1:500 ratio and cultured at 37 ^◦^C for 2 hours. In the induction group, 0.1mM or 0.5mM IPTG was added to the culture medium for induction, and the bacteria were cultured at 16 ^◦^C for 12 hours to obtain the culture for flow cytometry detection.

For flow cytometry analysis, 10 µL of the culture was added to 200 µL of phosphate-buffered saline (PBS) and placed in a 96-well plate. The BD Fortessa flow cytometer was used to measure the average fluorescence intensity of single cells, which represented the fluorescence protein expression levels in different samples. At least 10,000 samples were collected, and the background fluorescence of the cells was subtracted from all experimental groups before analysis using the FlowJo v10 software.

### 2.5 Experimental details for the evolution tasks of NUDT15 and VKOR1

#### 2.5.1 Construction of mutants

All mutant fragments were assembled onto the ampicin-resistant pLJM1-eGFP plasmid vector using the Seamless Cloning Kit (Beyotime). As an internal reference, mCherry was also cloned into the pLJM1 plasmid. The constructed plasmids were transformed into DH5*α* competent cells (Tiangen) for subsequent experiments. The mutant synthesis fragments were ordered by Qingke Biotech. The clone plasmids were extracted using the Tiangen DNA Extraction Kit, and the Sanger sequencing was completed by Qingke Company. The plasmids after identification were used for cell transfection experiments.

#### 2.5.2 Determination of abundance in cell lines

On day 1, HEK293T cells were seeded into a 6-well plate at a density of 500,000 cells per well. On day 2, 2 µg of pLJM1-NUDT15/VKOR1-eGFP and their mutants were diluted in 125 µL of OptiMEM with 1 µg of pLJM1mCherry. 9 µL of transfection reagent PEI (Yeasen Biotech) was diluted in 125 µL of OptiMEM in another tube. The PEI mixture was then added to the DNA mixture and incubated at room temperature for 15 min. Transfection mixtures were then added dropwise to HEK293T cells. 6 hours after transfection, the complete culture medium was replaced in 6-well plates. 48 hours after transfection, HEK293T cells were digested with trypsin and resuspended with PBS. The expression levels of fluorescent proteins were quantified using a BD Aria III FACS machine and at least 10,000 cells were collected for each sample. FlowJo v10 software was used to measure the median fluorescence intensity. Differences in abundance were characterized by the ratio of eGFP/mCherry signal.

### 2.6 Experimental details for the evolution task of AmeR

#### 2.6.1 Mutant strain preparation

Synthetic AmeR mutant genes were obtained from Tsingke Bioscience. All mutant fragments were cloned into the test plasmid (p15A replication origin, kanamycin resistance) using the ClonExpress assembly Kit (Vazyme). The constructed plasmids were transformed into *E. coli* DH5*α* competent cells (HT Health). The plasmid DNA was purified using the Tiangen Plasmid Mini Kit and verified by Sanger sequencing (Genewiz). Sequence-verified DH5*α* strains were used for subsequent experiments.

#### 2.6.2 Induced expression assay

Single colonies of mutant strains were cultured overnight in LB medium. The saturated cultures were diluted 1:50 in fresh LB medium containing 50 µg/mL kanamycin and incubated at 37 ^◦^C with 1000 rpm shaking for 1.5 hours. Then, 1 µL of this log-phase culture was diluted 1:500 into 500 µL fresh LB medium with 50 µg/mL kanamycin (induced group: supplemented with 1 mM IPTG; uninduced group: no IPTG). Both groups were incubated at 37 ^◦^C with 1000 rpm shaking for 5 hours before flow cytometry.

#### 2.6.3 Flow cytometry and data analysis

A 5 µL aliquot of bacterial culture was added to 195 µL PBS containing 2 mg/L kanamycin in 96-well U-bottom plates to arrest cell growth. Fluorescent protein expression was quantified using a Beckman Coulter Cytoflex S flow cytometer. FlowJo software (v10) was used to gate cells (≥ 10,000 events/sample) and calculate the median fluorescence intensity (MFI) for the induced and uninduced groups. Background fluorescence (MFI of bacteria harboring the empty backbone plasmid) was subtracted from all samples. Fold repression was calculated as: (Background-subtracted MFI of uninduced group) / (Background-subtracted MFI of induced group)

### 2.7 Experimental details for the evolution task of LCC-ICCG

#### 2.7.1 MLP filter construction

The experimentally determined hydrolysis activity data were used to train an MLP-based filter referencing previous research^6^. All protein sequences were encoded into embeddings using the ESM-C model^7^, followed by dimensionality reduction to a one-dimensional relative activity value through the MLP architecture. Multiple independent filters were trained on distinct cross-validation datasets, with ensemble-averaged predictions from these parallel models serving as a criterion to eliminate low-activity mutants during the screening process.

#### 2.7.2 DNA construction

All oligonucleotide primers were ordered from RuiBiotech. The gene of LCC-ICCG was ordered from Beijing RuiBiotech. Co., Ltd. The gene sequences encoding multi-site mutants were constructed through PCR amplification and Gibson assembly. LCC-ICCG and all mutants were cloned into pET-15b vectors. All DNA sequences were confirmed by direct sequencing.

#### 2.7.3 Protein expression and purification

All of the plasmids were transformed into *E. coli* BL21(DE3) competent cells. The single colony was cultured on 2×YT plates containing 100 µg/mL ampicillin, and then inoculated into 5 mL of 2×YT broth with the same antibiotics and grown in a shaker (37 ^◦^C, 220 rpm). The cultures were inoculated into 100 mL of fresh 2×YT broth containing the corresponding antibiotics. When OD600 reached 0.8 1.0, IPTG was added to a concentration of 0.5 mM to induce the expression of the target protein. The cultures were then shaken at 16 ^◦^C for 12 hours. Cells were harvested by centrifugation (4000 rpm × 20 min, 4 ^◦^C). The harvested cell pellets were resuspended in 15 mL lysis buffer (50 mM NaH_2_PO_4_, 300 mM NaCl, 10 mM imidazole, pH 8.0) and lysed by ultrasonication. The supernatant was then collected after centrifugation (13000 rpm × 45 min, 4 ^◦^C), and mixed with Ni-NTA resin (GE Healthcare. Inc.). The mixture was incubated at 4 ^◦^C for 30 min, then loaded to an empty column and washed by wash buffer (50 mM NaH_2_PO_4_, 300 mM NaCl, 20 mM imidazole, pH 8.0) for five resin volumes, and finally eluted by elution buffer (50 mM NaH_2_PO_4_, 300 mM NaCl, 250 mM imidazole, pH 8.0). The eluted products were collected and subsequently buffer-exchanged into Tris buffer (20 mM Tris, 150 mM NaCl, pH 8.0) using a desalting column.

#### 2.7.4 Catalytic activity assay

The enzymatic activities of LCC-ICCG and its mutants were determined using *p*-nitrophenyl acetate (*p*NP-C2) and polyethylene terephthalate (PET) as substrates, respectively. For *p*NP-C2 hydrolysis, protein samples were diluted to 250 nM with potassium buffer (100 mM K_2_HPO_4_, pH 8.0). The concentrated stock of the substrate is a 10 mM acetonitrile solution of *p*NP-C2. The stock solution was diluted to 1.11 mM with potassium buffer before the test. 20 µL 250 nM protein samples were then mixed with 180 µL 1.11 mM *p*NP-C2 (in potassium buffer, sigma) in a 96-well microplate, and the absorbance at 415 nm was immediately measured in kinetic mode at room temperature using the EnSpire multimode plate reader (PerkinElmer Inc.). The absorbance of *p*-nitrophenol with varying amounts (0, 20, 40, 60, 80, 100, 120, 200 nmol) at 415 nm was also measured to get the standard curve, which was used to calculate the amounts of *p*-nitrophenol that were generated during the reaction. A linear range of samples plot was used to calculate the activity. For PET depolymerization, the activity was quantitatively determined by measuring the concentration of terephthalic acid (TPA) released in the reaction system. The amorphous Gf-PET film (Goodfellow, 250 µm thickness, product number ES301445, *ϕ*5mm, approximately 5 mg) was soaked in 100 µL of 100 mM potassium phosphate buffer (100 mM K_2_HPO_4_, pH 8.0) with 1.5 µM of purified enzyme at 65 ^◦^C for 8 hours. In ultra-performance liquid chromatography (UPLC) analysis, the mobile phase consisted of 70% buffer A and 30% buffer B (Buffer A: 0.1% formic acid in distilled water; Buffer B: acetonitrile). The flow rate was 0.8 mL/min and the separation was carried out at 25 ^◦^C with detection performed at 254 nm. The absorbance of TPA with varying amounts (0.1, 0.2, 0.5, 1.0, 2.0, 5.0 mmol) at 254 nm was also measured to get the standard curve, which was used to calculate the amounts of TPA generated during the reaction. The enzymatic activity was calculated by quantifying the TPA concentration in the reaction system through UPLC peak area analysis calibrated against a standard curve.

#### 2.7.5 Determination of apparent melting temperature

To determine the apparent melting temperature (*T_m_*), a fluorescence-based thermal stability assay was employed. A 1 µL 50-fold diluted SYPRO Orange dye (Molecular Probes, Life Technologies, USA) was added to a 20 µL protein solution. The mixture was placed in a thin-walled 96-well PCR plate, sealed with optical-quality sealing tape. The mixture was heated in a CFX 96 real-time polymerase chain reaction (PCR) system (BioRad, Hercules, CA, USA) from 50 ^◦^C to 100 ^◦^C with a heating rate of 1.2 ^◦^C/sec.

### 2.8 PETase activity reported in literature and patents

In the following tables, we show the catalytic activity and thermostability data of mutants derived from two PETases (PDBID: 4EBO and 7EOA) collected in a previous investigation^8^ from the literature and patents. We used these data to train the reward model during the first round of RelaVDEP-based sequence optimization for LCC-ICCG.

**Supplementary Table 29.**
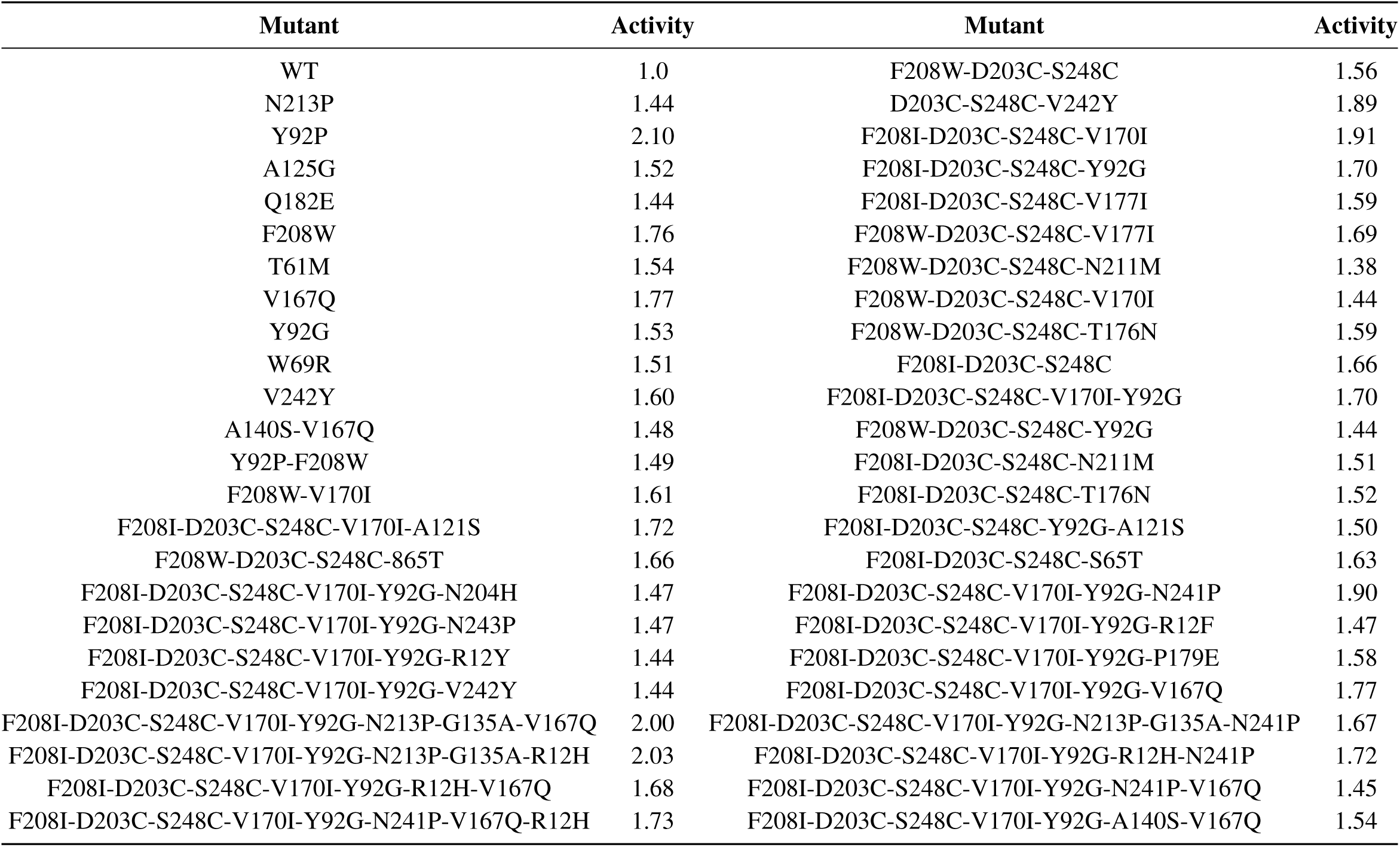
Reported PETase activity data (template: 4EB0)

**Supplementary Table 30.** Reported PETase activity data (template: 7EOA)

| Mutant | Activity | Mutant | Activity | Mutant | Activity |
| --- | --- | --- | --- | --- | --- |
| WT | 1.0 | W190M | 0.04 | W190H | 0.56 |
| H191L | 0.89 | F243D | 1.14 | F243G | 1.54 |
| H191S | 1.20 | H191Y | 1.13 | F243S | 1.16 |
| W104L | 1.26 | H218S-F222I-W104L | 2.75 | W104C | 1.12 |
| W104D | 1.76 | H218S-F222I-W104D | 2.12 | H218S-F222I | 2.25 |
| W104G | 1.39 | H218S-F222I-W104G | 2.50 | H218S-F222I-S64P | 1.91 |
| H164S | 0.22 | H218S-F222I-H164S | 0.27 | H218S-F222I-L102V | 1.78 |
| H164Q | 0.56 | H218S-F222I-H164Q | 0.51 | H218S-F222I-F127T | 1.65 |
| M166L | 0.06 | H218S-F222I-M166L | 0.11 | H218S-F222I-I178H | 2.13 |
| M166D | 0.09 | H218S-F222I-M166D | 0.26 | H218S-F222I-Q202V | 2.31 |
| W190L | 0.01 | H218S-F222I-W190L | 0.09 | H218S-F222I-D238K | 2.22 |
| W190S | 0.27 | H218S-F222I-W190S | 0.29 | H218S-F222I-S146V | 2.02 |
| W190D | 0.01 | H218S-F222I-W190D | 0.05 | H218S-F222I-S287A | 2.33 |
| H191M | 0.75 | H218S-F222I-H191M | 0.33 | H218S-F222I-A209R | 2.51 |
| H191D | 0.72 | H218S-F222I-H191D | 0.82 | H218S-F222I-T192Y | 2.16 |
| F243I | 1.60 | H218S-F222I-F243I | 2.81 | H218S-F222I-W104H | 2.56 |
| F243T | 1.61 | H218S-F222I-F243T | 2.68 | H218S-F222I-W104S | 2.45 |
| F243N | 1.51 | H218S-F222I-F243N | 1.53 | H218S-F222I-W104C | 1.98 |
| H218S | 2.00 | H218S-F222I-S57P | 2.27 | H218S-F222I-T85E | 2.41 |
| W104S | 1.11 | H218S-F222I-H112W | 2.20 | H218S-F222I-W104R | 1.00 |
| W104H | 1.20 | H218S-F222I-H164L | 0.10 | H218S-F222I-H164E | 0.02 |
| W104R | 0.24 | H218S-F222I-H164F | 0.34 | H218S-F222I-M166S | 0.01 |
| H164L | 0.15 | H218S-F222I-M166F | 0.22 | H218S-F222I-W190M | 0.06 |
| H164E | 0.21 | H218S-F222I-W190H | 0.97 | H218S-F222I-H191L | 0.25 |
| H164F | 0.41 | H218S-F222I-H191S | 0.42 | H218S-F222I-H191Y | 1.10 |
| M166S | 0.12 | H218S-F222I-F243S | 1.40 | H218S-F222I-F243G | 2.94 |
| M166F | 0.11 | H218S-F222I-F243D | 1.48 | H218S-F222I-A251C-A281C | 2.27 |

**Supplementary Table 31.** Reported PETase activity data (template: 7EOA)

| Mutant | Activity |
| --- | --- |
| H218S-F222I-D238K-A251C-A281C | 2.21 |
| H218S-F222I-D238K-A251C-A281C-A209R | 2.17 |
| H218S-F222I-D238K-A251C-A281C-A209R-W104G-F243G | 3.01 |
| H218S-F222I-D238K-A251C-A281C-A209R-W104G-F243I | 3.21 |
| H218S-F222I-D238K-A251C-A281C-A209R-W104G-F243T | 2.68 |
| H218S-F222I-D238K-A251C-A281C-A209R-W104H-F243G | 3.32 |
| H218S-F222I-D238K-A251C-A281C-A209R-W104H-F243I | 3.20 |
| H218S-F222I-D238K-A251C-A281C-A209R-W104H-F243T | 3.25 |
| H218S-F222I-D238K-A251C-A281C-A209R-W104L-F243G | 3.46 |
| H218S-F222I-D238K-A251C-A281C-A209R-W104L-F243I | 3.24 |
| H218S-F222I-D238K-A251C-A281C-A209R-W104L-F243T | 3.55 |
| H218S-F222I-D238K-A251C-A281C-A209R-W104S-F243G | 3.49 |
| H218S-F222I-D238K-A251C-A281C-A209R-W104S-F243I | 3.29 |
| H218S-F222I-D238K-A251C-A281C-A209R-W104S-F243T | 3.17 |

**Supplementary Table 32.** Reported PETase thermostability data (template: 4EB0)

| Mutant | $T_m(^{\circ}\text{C})$ | Mutant | $T_m(^{\circ}\text{C})$ |
| --- | --- | --- | --- |
| F208I-D203C-S248C-V170I | 92.10 | F208W-D203C-S248C-N211M | 98.10 |
| F208I-D203C-S248C-Y92G | 94.00 | F208I-D203C-S248C-V177I | 91.50 |
| F208W-D203C-S248C-Y92G | 98.00 | F208W-D203C-S248C-V177I | 95.90 |
| F208I-D203C-S248C-N211M | 94.50 | F208W-D203C-S248C-N211M | 98.10 |
| F208W-D203C-S248C-V170I | 96.40 | F208I-D203C-S248C-V170I-Y92G | 94.60 |
| F208I-D203C-S248C-T176N | 90.30 | F208I-D203C-S248C-V170I-A121S | 92.20 |
| F208W-D203C-S248C-T176N | 95.00 |  |  |

**Supplementary Table 33.** Reported PETase thermostability data (template: 7EOA)

| Mutant | $T_m$ (°C) | Mutant | $T_m$ (°C) | Mutant | $T_m$ (°C) |
| --- | --- | --- | --- | --- | --- |
| H191Y | 87.00 | H218S | 80.50 | H218S-F222I-H191L | 76.00 |
| H191M | 86.50 | H191D | 96.00 | H218S-F222I-H191Y | 77.00 |
| F243S | 94.00 | F243T | 90.50 | H218S-F222I-W104L | 72.00 |
| F243N | 94.50 | F243G | 92.00 | H218S-F222I-W104S | 72.50 |
| W104L | 85.50 | F243I | 92.50 | H218S-F222I-F243I | 81.00 |
| W104S | 85.00 | F243D | 97.50 | H218S-F222I-T85E | 84.50 |
| W104C | 87.00 | H218S-F222I | 85.00 | H218S-F222I-L102V | 80.00 |
| W104H | 88.50 | H218S-F222I-W104C | 72.50 | H218S-F222I-H112W | 85.00 |
| W104D | 84.50 | H218S-F222I-W104H | 75.50 | H218S-F222I-F127T | 84.00 |
| W104R | 85.00 | H218S-F222I-W104D | 73.50 | H218S-F222I-S146V | 84.50 |
| W104G | 85.50 | H218S-F222I-W104R | 74.50 | H218S-F222I-I178H | 83.50 |
| H164L | 82.00 | H218S-F222I-W104G | 71.50 | H218S-F222I-H191D | 88.00 |
| H164S | 90.50 | H218S-F222I-H164L | 72.50 | H218S-F222I-T192Y | 85.00 |
| H164E | 84.50 | H218S-F222I-H164S | 79.00 | H218S-F222I-Q202V | 76.00 |
| H164Q | 91.00 | H218S-F222I-H164E | 75.00 | H218S-F222I-A209R | 87.00 |
| H164F | 83.00 | H218S-F222I-H164Q | 80.50 | H218S-F222I-D238K | 93.50 |
| M166L | 91.50 | H218S-F222I-H164F | 74.00 | H218S-F222I-S287A | 80.00 |
| M166S | 93.50 | H218S-F222I-M166L | 79.00 | H218S-F222I-H191S | 86.00 |
| M166D | 92.50 | H218S-F222I-M166S | 87.00 | H218S-F222I-S64P | 85.50 |
| M166F | 95.00 | H218S-F222I-M166D | 83.00 | H218S-F222I-F243S | 83.00 |
| W190L | 88.00 | H218S-F222I-M166F | 84.00 | H218S-F222I-F243T | 84.00 |
| W190M | 89.50 | H218S-F222I-W190L | 78.00 | H218S-F222I-F243D | 86.50 |
| W190S | 84.50 | H218S-F222I-W190M | 78.50 | H218S-F222I-F243N | 85.00 |
| W190H | 84.00 | H218S-F222I-W190S | 76.50 | H218S-F222I-F243G | 81.00 |
| W190D | 84.00 | H218S-F222I-W190H | 76.50 | H218S-F222I-S57P | 85.00 |
| H191L | 79.50 | H218S-F222I-W190D | 75.00 |  |  |
| H191S | 91.00 | H218S-F222I-H191M | 78.00 |  |  |

**Supplementary Table 34.** Reported PETase thermostability data (template: 7EOA)

| Mutant | $T_m(^{\circ}\text{C})$ |
| --- | --- |
| H218S-F222I-A251C-A281C | 88.00 |
| H218S-F222I-D238K-A251C-A281C | 95.50 |
| H218S-F222I-D238K-A251C-A281C-A209R | 97.00 |
| H218S-F222I-D238K-A251C-A281C-A209R-W104G-F243G | 83.00 |
| H218S-F222I-D238K-A251C-A281C-A209R-W104G-F243I | 83.00 |
| H218S-F222I-D238K-A251C-A281C-A209R-W104G-F243T | 86.00 |
| H218S-F222I-D238K-A251C-A281C-A209R-W104H-F243G | 86.00 |
| H218S-F222I-D238K-A251C-A281C-A209R-W104H-F243I | 85.00 |
| H218S-F222I-D238K-A251C-A281C-A209R-W104H-F243T | 87.50 |
| H218S-F222I-D238K-A251C-A281C-A209R-W104L-F243G | 82.50 |
| H218S-F222I-D238K-A251C-A281C-A209R-W104L-F243I | 82.00 |
| H218S-F222I-D238K-A251C-A281C-A209R-W104L-F243T | 84.00 |
| H218S-F222I-D238K-A251C-A281C-A209R-W104S-F243G | 83.00 |
| H218S-F222I-D238K-A251C-A281C-A209R-W104S-F243I | 83.50 |
| H218S-F222I-D238K-A251C-A281C-A209R-W104S-F243T | 85.00 |

## 3 Pseudo Codes and Algorithms

### Algorithm 1

Representation Network

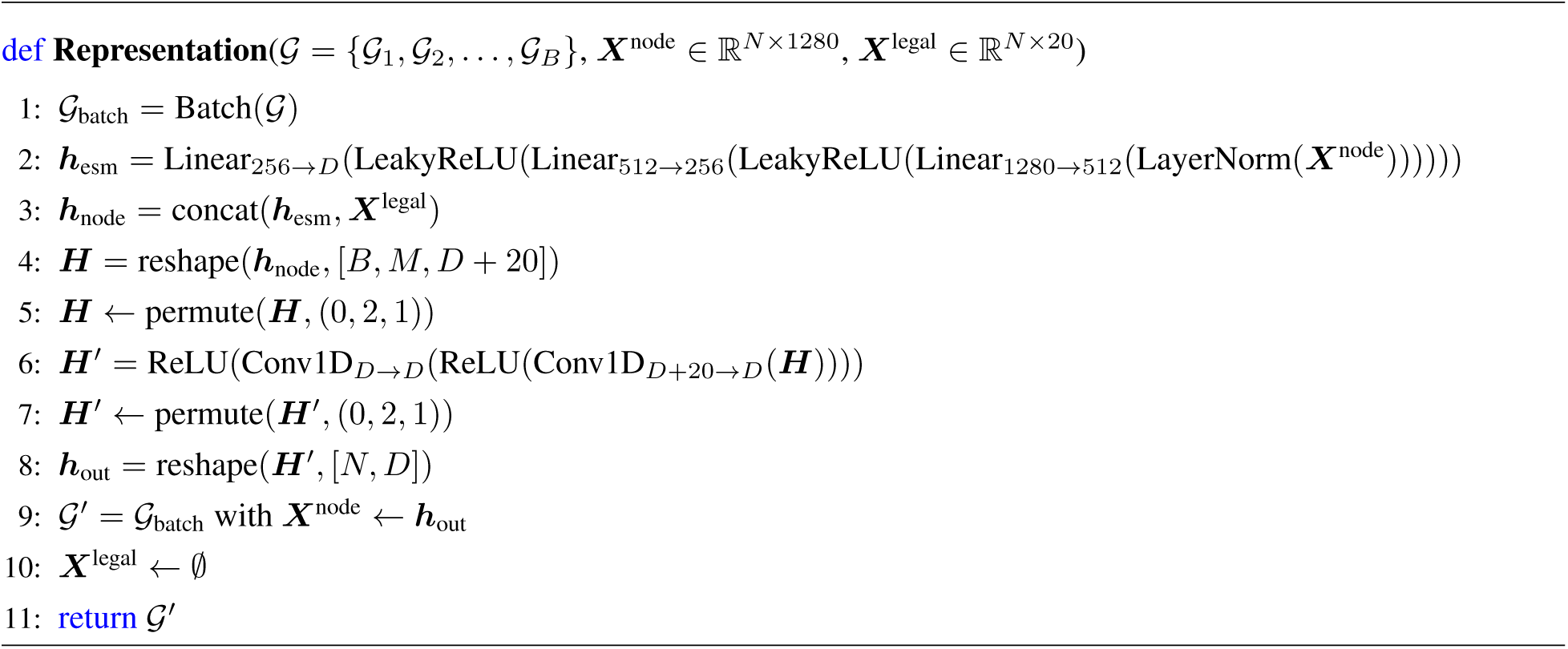

### Algorithm 2

Prediction Network

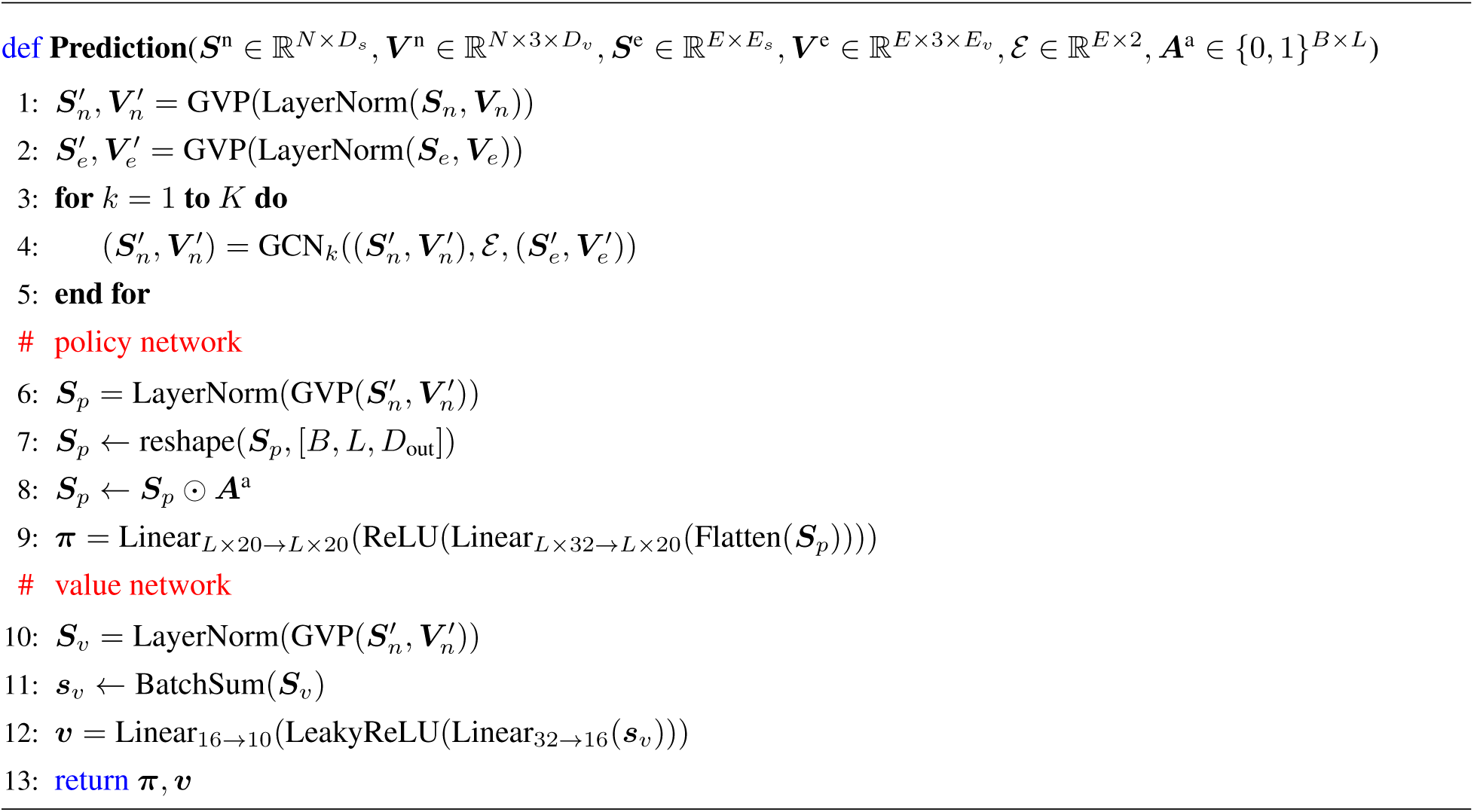

### Algorithm 3

Dynamics Network

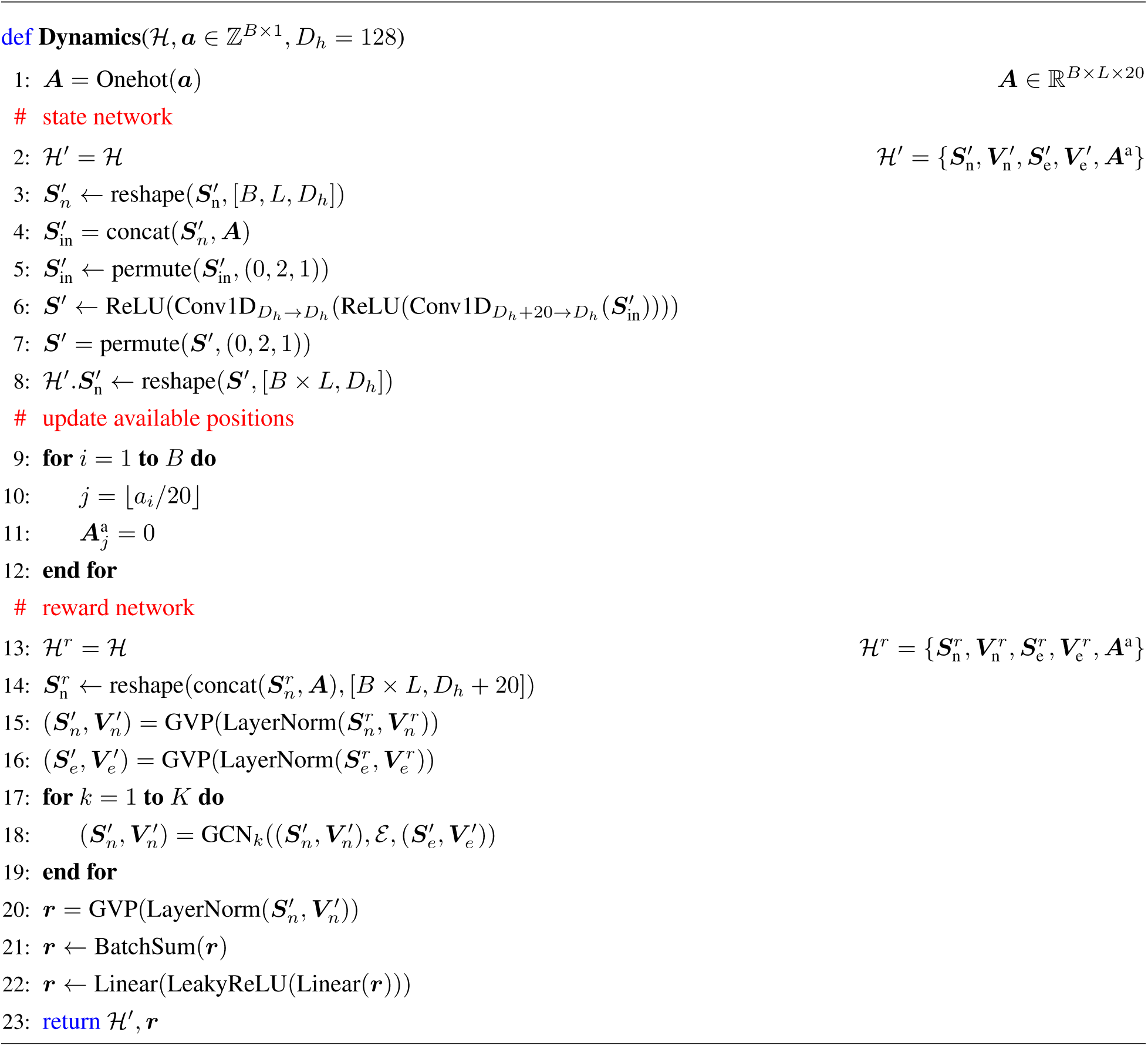

### Algorithm 4

Training loop

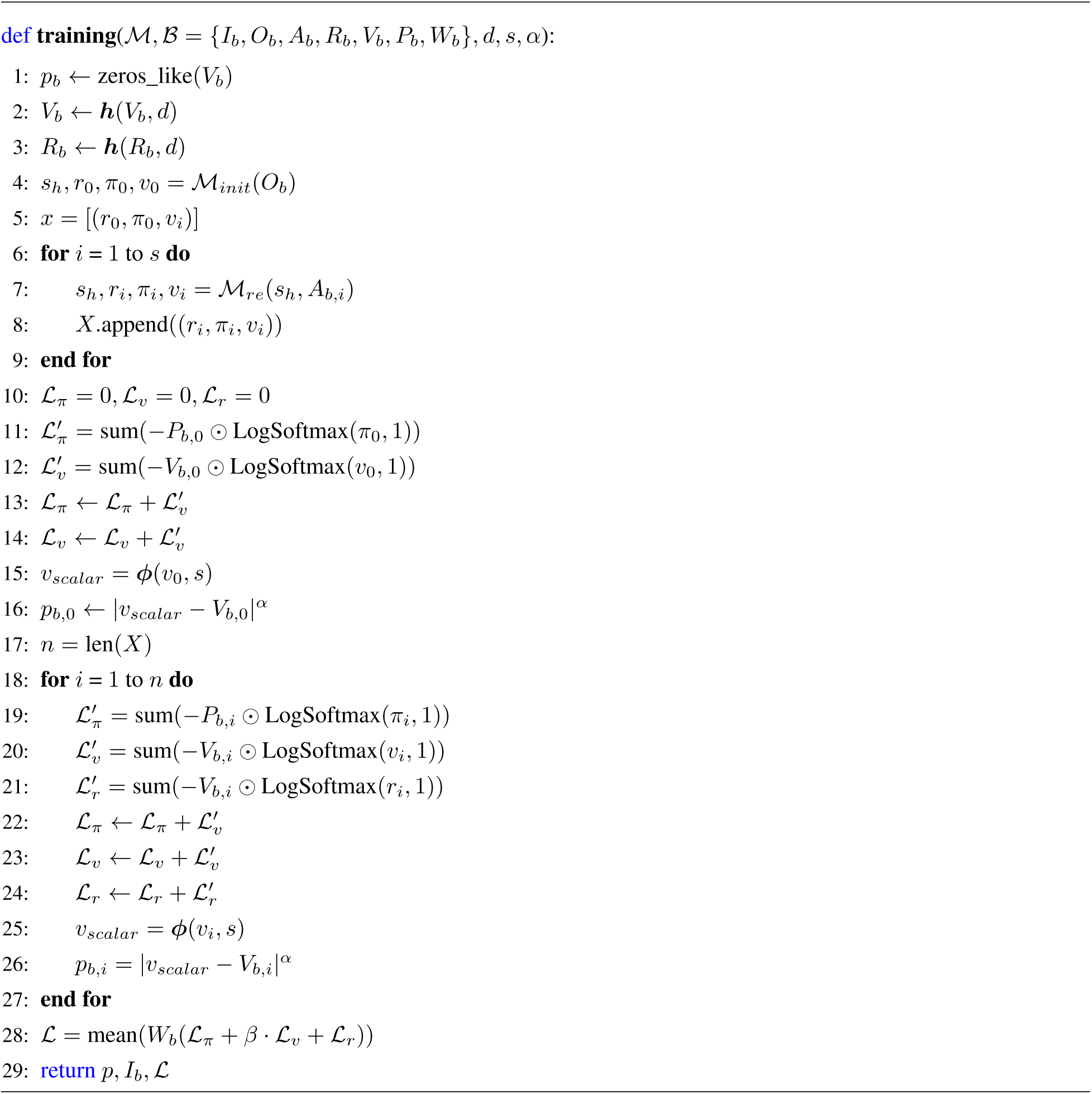

### Algorithm 5

Calculate the mutation frequency of input mutants

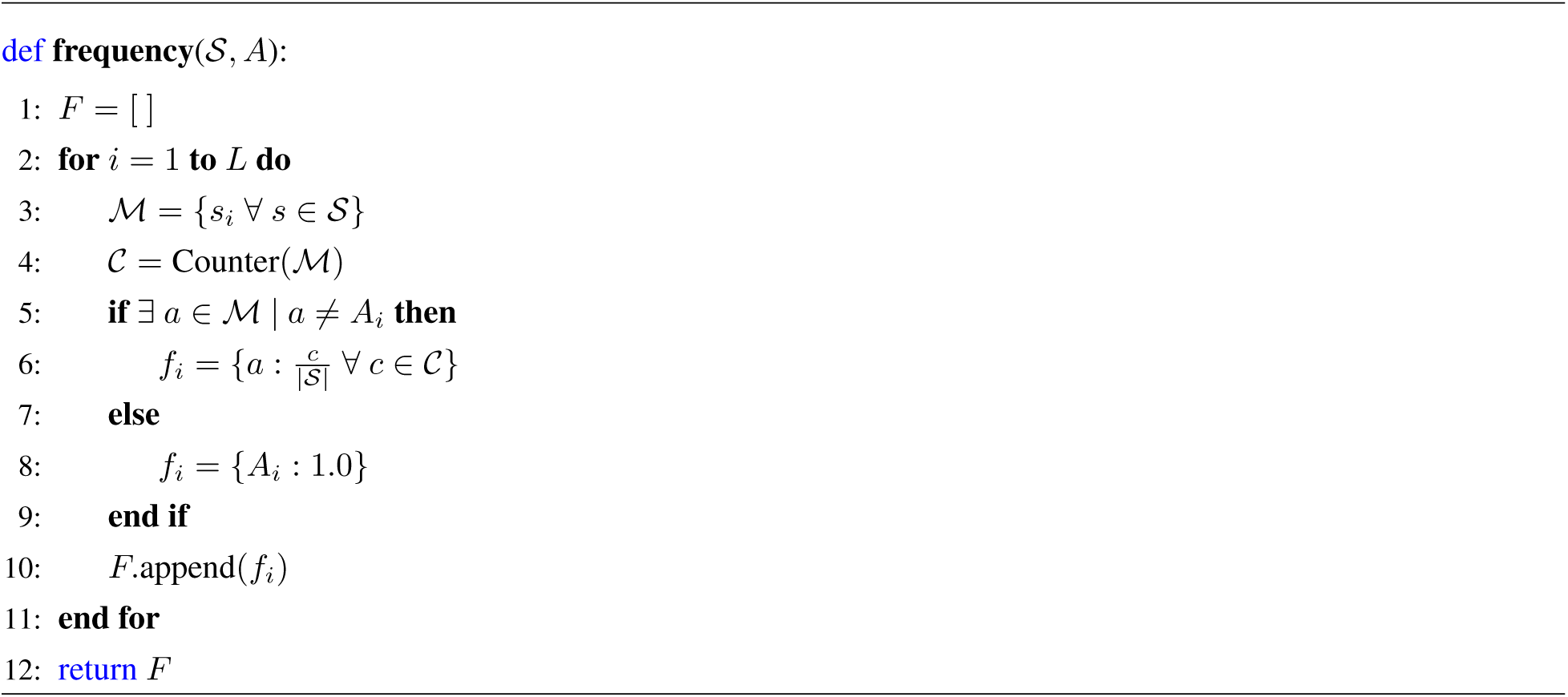

### Algorithm 6

Calculate the target to be optimized

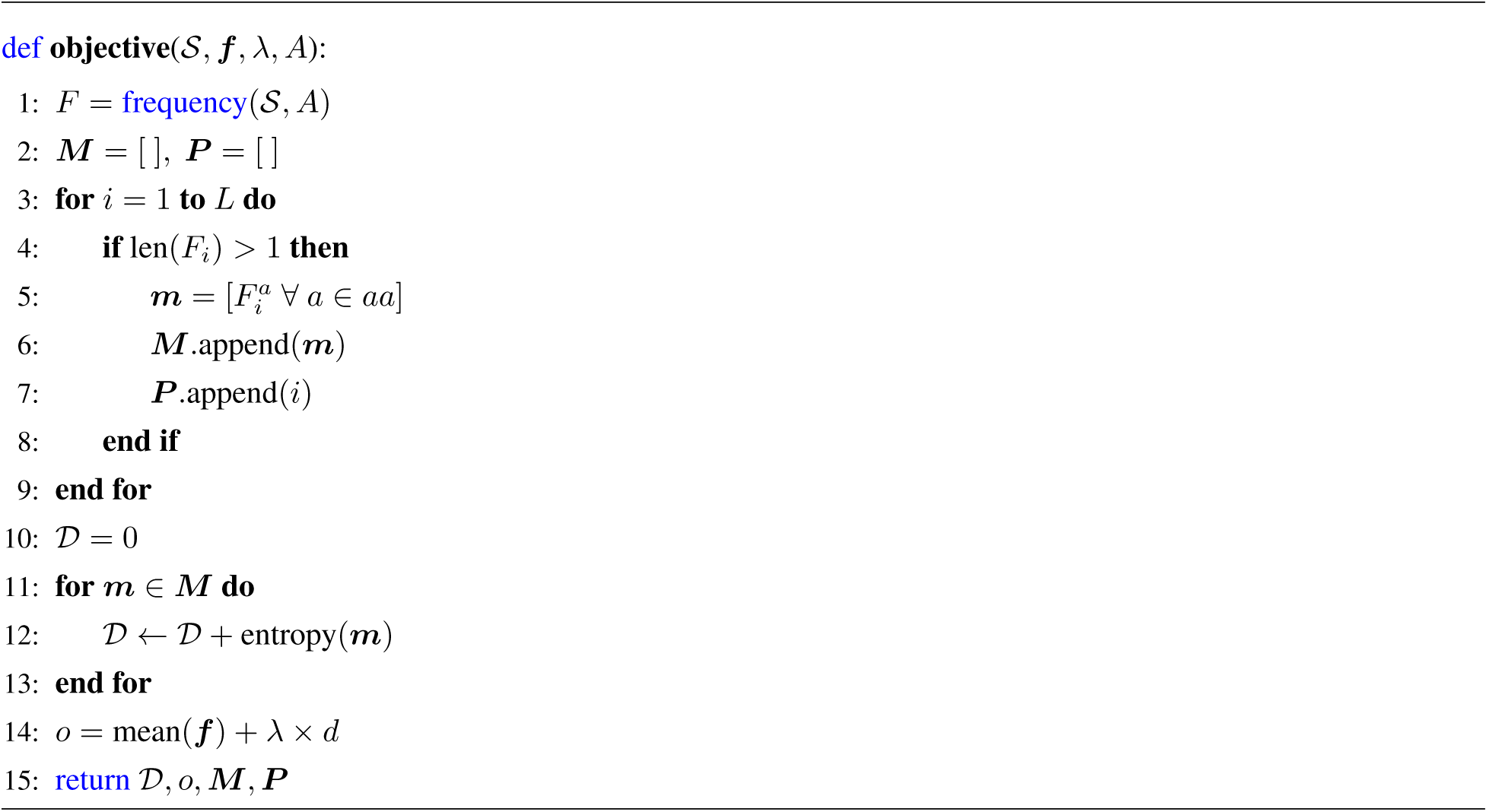

### Algorithm 7

Construct optimized mutant library

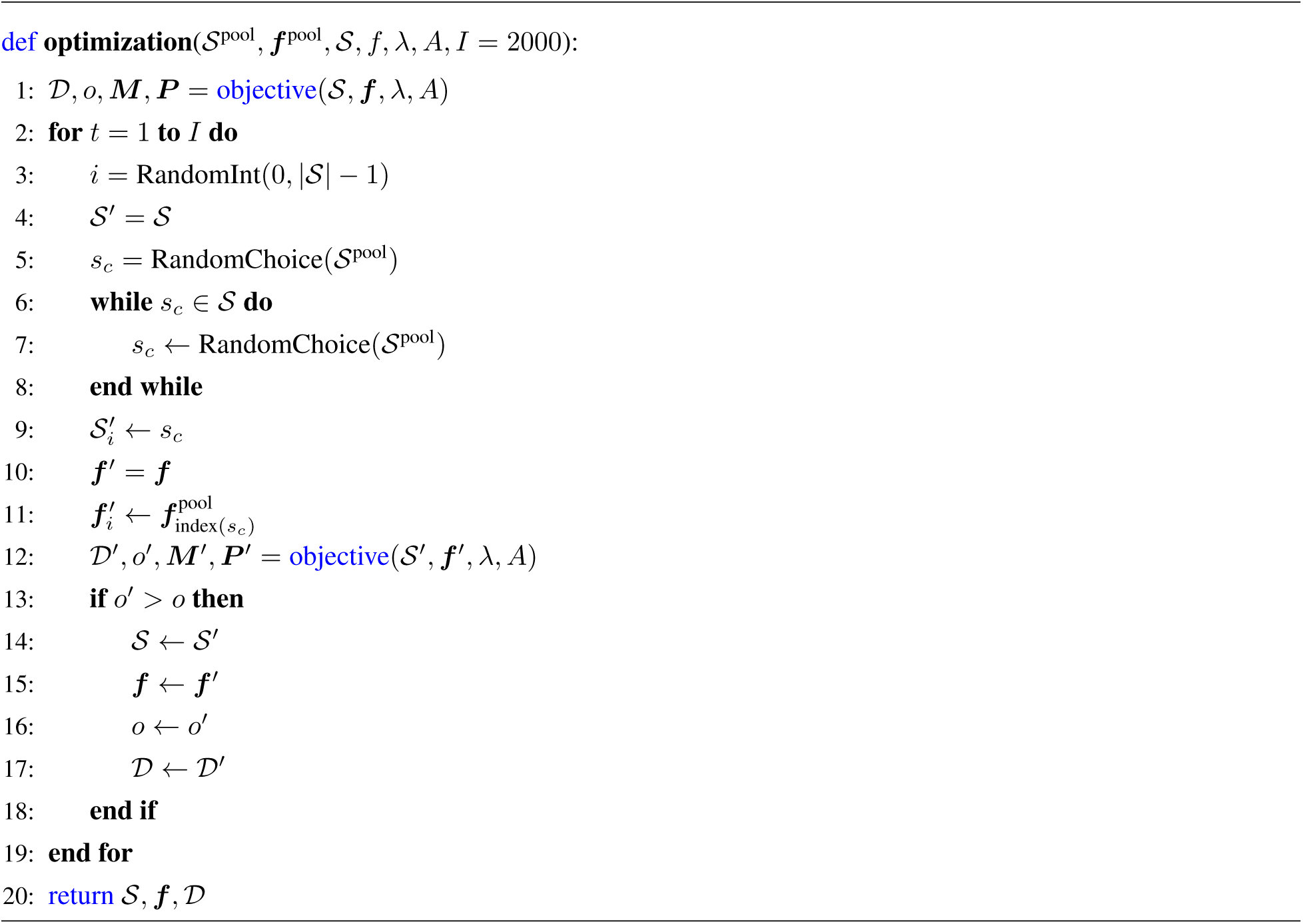

## Notes

### Competing Interest Statement

The authors have declared no competing interest.

